# Identification of spatially variable genes with graph cuts

**DOI:** 10.1101/491472

**Authors:** Ke Zhang, Wanwan Feng, Peng Wang

## Abstract

Single-cell gene expression data with positional information are critical to dissect mechanisms and architectures of multicellular organisms, but the potential is limited by current data analysis strategies. Here, we present scGCO (single-cell graph cuts optimization), a method based on fast optimization of Markov Random Fields with graph cuts, to identify spatially viable genes. Extensive benchmarking demonstrated that scGCO delivers superior performance with optimal segmentation of spatial patterns, and can process millions of cells in a timely manner owing to its linear scalability.

## Introduction

Systematic assessment of the spatial context of gene expression is a cornerstone in understanding mechanistic functionality and molecular organization of tissues and organs^1^. Currently, two main classes of experimental approaches have been established to measure spatial transcriptomics. Utilizing probes for individual RNA molecules to directly quantify gene expression *in situ*, image-based single-cell spatial transcriptomics, such as seqFISH^2^ and MERFISH^3^, can measure hundreds of genes in an entire tissue section. On the other hand, by combining single-cell RNA-seq data with prerecorded coordinate information, spatial gene expression can be generated for hundreds of cells at the genome-scale^4,5^.

The central task of analyzing spatial transcriptomics is to identify genes with spatially viable expression patterns (hereafter referred to as spatial genes). The first generation of methods mainly identify spatial genes by comparing gene expression among arbitrarily selected regions using procedures such as ANOVA^2,4^. However, the boundaries of selected regions are not rigorously defined, which could limit the detection power of subsequent statistical methods. More importantly, scientific discovery of novel spatial regions is not possible. Recently, two methods based on Gaussian process^6^ or marked point process^7^ were developed to specifically identify spatial genes. However, benchmarking showed that these methods reported a substantially lower number of spatial genes than methods directly comparing preselected regions using the same data^4,6,7^. Moreover, these methods scale poorly with a cubic or quadratic growth rate in the number of cells^6,7^. This may substantially limit their utilities, as the spatial transcriptomics scales beyond hundreds of cells.

Here, we present a novel algorithm, single-cell graph cuts optimization (scGCO), to identify spatial genes. A crucial insight of scGCO is that identifying spatial genes is analogous to identifying objects from an image, also known as image segmentation, which is a classical problem in computer vision that can be solved optimally with graph cuts algorithms^8^. Consistent with the theoretical advantages of graph cuts, scGCO demonstrated superior performance against existing methods over a wide range of spatial transcriptomics data and can scale to millions of cells. We have made scGCO available as a python package to allow optimal analysis of spatial transcriptomics data.

## Results

### Overview of scGCO algorithm

To apply graph cuts to spatial gene expression data, scGCO first performs Delaunay triangulation on spatial coordinates of cells to generate a sparse graph representation of cell locations (Fig. 1a, 1b). The graph can then be analyzed by graph cuts algorithms to identify cuts that minimize the energy of the underlying Markov random fields (MRFs), where resulting subgraphs correspond to clusters of cells with similar expression values (Fig. 1c). The identified spatial patterns can then be visualized by Voronoi tessellation, and statistical significance of identified spatial genes can be evaluated with a homogeneous spatial Poisson process (Fig. 1d).

**Figure 1.**
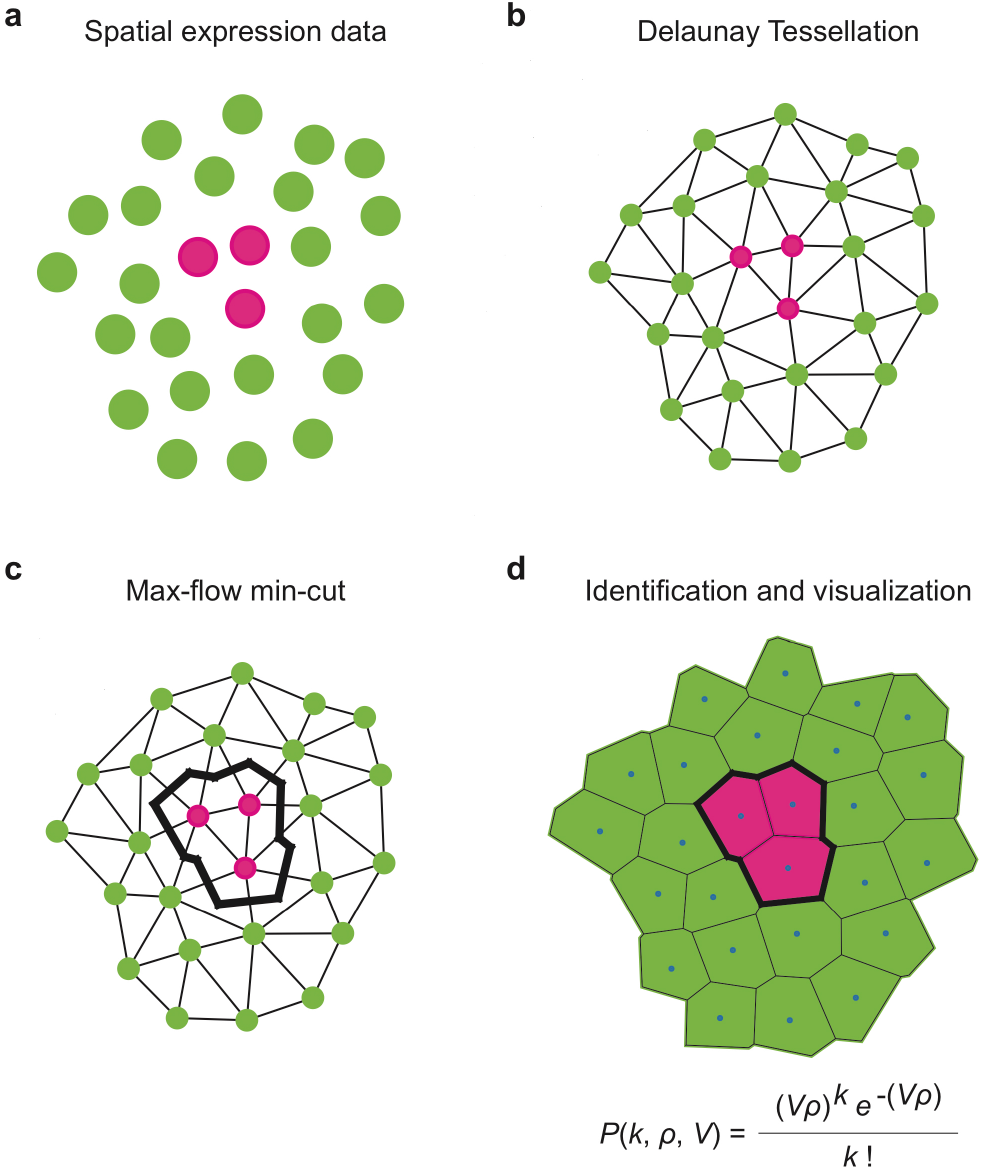
Overview of scGCO for spatial gene identification. (**a**) An example of spatial gene expression data. Each dot represents a cell and is placed according to its spatial coordinates. The gene expression level is high in magenta colored cells and low in turquoise colored cells. (**b**) Transforming the spatial gene expression data into a graph representation via Delaunay tessellation. (**c**) The classical max-flow min-cut algorithm cuts the spatial gene expression graph into two subgraphs, one consisting of cells overexpressing the gene and one consisting of cells underexpressing the gene. (**d**) Voronoi tessellation produces a visual representation of the identified spatial expression pattern. Thicker lines highlight the subgraph boundaries identified by graph cuts. The statistical significance of spatial distribution of gene expression is determined by a homogeneous spatial Poisson distribution.

### scGCO provides sensitive and robust identification of spatial genes

We first applied scGCO to spatial transcriptomics data from mouse olfactory bulb (MOB)^4^. In the original study, Ståhl et al. directly compared the expression of cells in the granular cell layer (GCL) against cells in the glomerular layer (GL), and reported 170 differentially expressed genes^4^. Because MOB consists of 5 different layers^4^, hundreds to thousands of genes could be differentially expressed between these regions, and hence are spatially viable if we assume that each pair of regions generates a similar number of differentially expressed genes to that of GCL vs. GL.

Two recently published methods, spatialDE^5^ and trendSceek^6^, were especially designed to identify spatial genes. Because trendSceek can only identify < 100 genes in two out of the twelve replicates of MOB data^7^, we focused on the comparison with spatialDE. We first applied scGCO to replicate 11 of the MOB data, which spatialDE analyzed extensively in their study^6^. Strikingly, scGCO identified 16-fold more spatial genes (1,131 genes, FDR < 0.01) than spatialDE (67 genes, FDR < 0.05), and reproduced a majority of spatialDE identified genes (59 of the 67) (Fig. 2a). Because biological functions are carried out by modules or networks of genes that are highly correlated, we expect that the spatial genes should also share similar spatial patterns. Indeed, genes identified by scGCO formed four tight clusters when projected onto a low-dimensional space via t-distributed stochastic neighbor embedding (t-SNE)^9^ (Fig. 2b). Moreover, direct visualization of spatial gene expression patterns of representative genes confirmed that a distinct spatial pattern is associated with each cluster (Fig. 2c). To exhaustively validate the predictions, we plotted and visually examined all 1,131 genes identified by scGCO and confirmed that the vast majority of identified genes indeed display valid spatial patterns that resemble representative genes from each cluster (Supplementary File 1). Finally, five out of the top ten enriched gene sets are neuron-related, confirming that the large number of spatial genes identified by scGCO demonstrate significant biological relevance (Supplementary Fig. 1).

**Figure 2.**
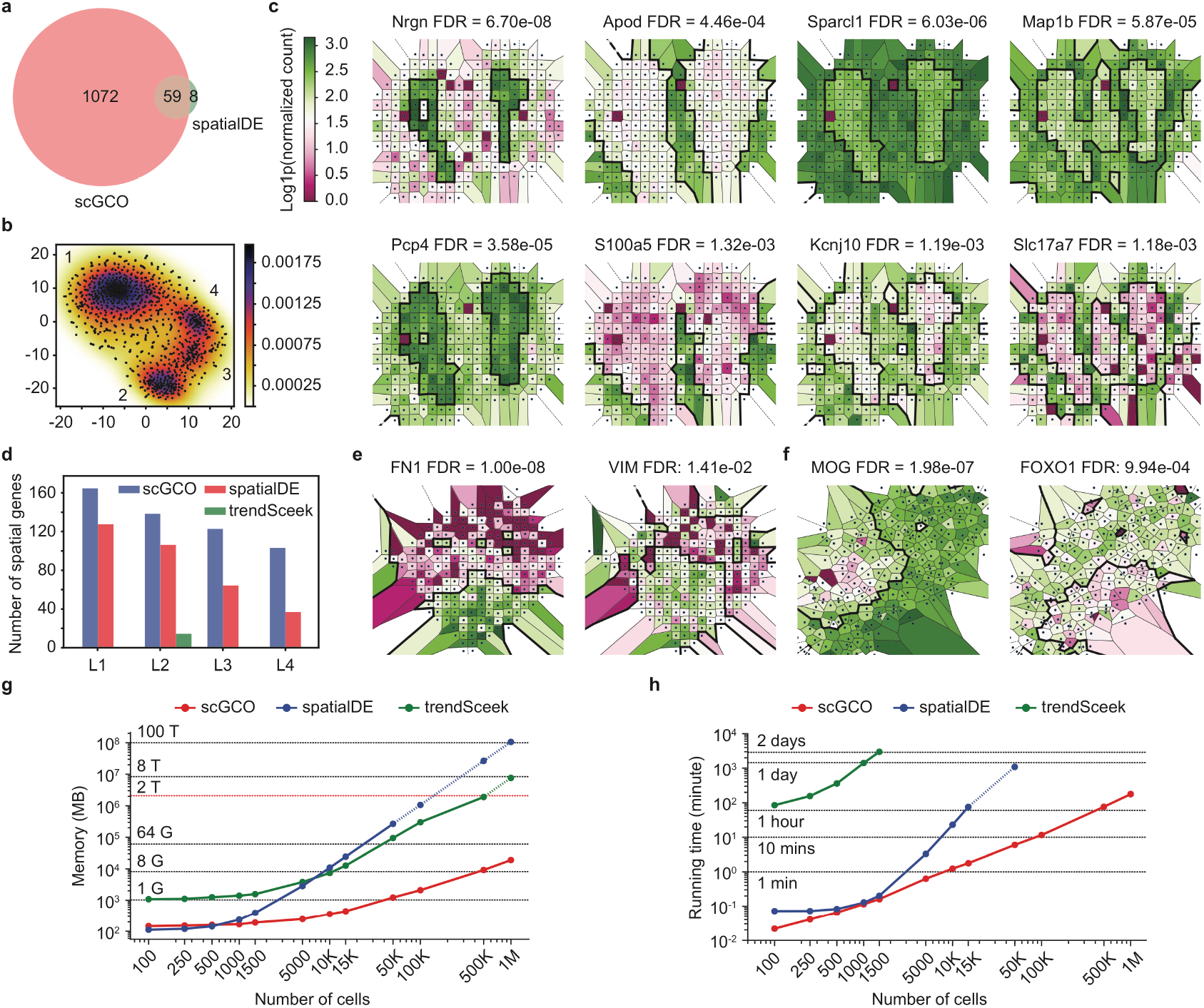
scGCO delivers superior performance in analyzing spatial gene expression data. (**a**) Venn diagram showing the set relationship among spatial genes identified by scGCO (FDR < 0.01) and spatialDE (FDR < 0.05) using mouse olfactory bulb data (replicate 11). (**b**) t-SNE for all significant spatial genes identified by scGCO shown in **a**. Numbers indicate indexes of identified clusters. (**c**) Representative Voronoi diagrams showing spatial expression patterns. Two examples per column were shown for each cluster in **b**. Polygons representing cells are colored according associated gene expression levels. The boundaries for segments of significant spatial expression patterns are depicted by thicker black lines. (**d**) Bar charts showing the number of significant spatial genes identified by scGCO, spatialDE and trendSceek, for breast cancer data (FDR < 0.05). (**e**) Representative Voronoi diagrams showing spatial expression patterns identified in breast cancer data by scGCO (layer 2). (**f**) Representative Voronoi diagrams showing spatial expression patterns identified in mouse hippocampus seqFISH data by scGCO (field 43). (**g**) Graph showing the memory requirement of scGCO, spatialDE and trendSceek in the number of cells (100 genes). Dotted line indicates memory extrapolated from measured data. (**h**) Graph showing the running time of scGCO, spatialDE and trendSceek in the number of cells (100 genes). Dotted line indicates running time extrapolated from measured data.

We next analyzed all 12 replicates of the MOB data. Similar to the results for replicate 11, scGCO consistently identified substantially more spatial genes than spatialDE and trendSceek in all replicates (Supplementary Fig. 2). Reassuringly, four clusters with the minor cluster detectable in nine replicates were consistently recovered by t-SNE analysis (Supplementary Fig. 3). Direct visualization confirmed the validity of the identified spatial patterns in all replicates, and the results of two replicates (1 and 10) with a large number of identified genes are provided in supplementary materials (supplementary Fig. 4 and 5, supplementary Files 2 and 3). In contrast, genes identified by spatialDE formed fewer clusters, and each cluster contained many fewer genes (supplementary Fig. 6). Importantly, scGCO also reproduced majority of the genes differentially expressed between GCL vs. GL layers, confirming that scGCO could identify spatial genes beyond direct region comparison (Supplementary Fig. 7).

We next investigated robustness of the algorithms by comparing genes that were reproducibly identified across all 12 biological replicates. ScGCO consistently reproduced more spatial genes and had a smaller percentage of unreproducible genes than spatialDE (35% v.s. 46%) (Supplementary Fig. 8). Moreover, the reproducible genes identified by scGCO are highly enriched with neuron-related gene ontologies, further confirming the validity of identified spatial genes (Supplementary Fig. 8).

### scGCO is applicable to a wide variety of spatial transcriptomics data

We next applied scGCO to spatial gene expression data from breast cancer biopsies, which were generated using the same protocol as the MOB data^4^. As expected, scGCO consistently identified more spatial genes than both spatialDE and trendSceek (Fig. 2d, supplementary Fig. 9). Interestingly, genes identified by scGCO consistently formed three clusters using t-SNE across all four replicates, while genes identified by spatialDE failed to maintain consistent clustering patterns, suggesting that scGCO is not only more sensitive but also more robust (Supplementary Fig. 10). Indeed, scGCO consistently reproduced more spatial genes than spatialDE when biological repeats were compared, and had a lower percentage of unreproducible genes (46.1% vs. 57.6%). Reassuringly, reproducible genes identified by scGCO are enriched with metastasis-related GO terms such as focal adhesion, confirming their biological relevance (Fig. 2e, Supplementary Fig. 11).

We next tested scGCO using seqFISH data from mouse hippocampus. The hippocampus data contain 21 fields with variable quality, and consequently, the number of identified spatial genes ranged from single digits to over two hundred (Supplementary Fig. 12). Despite this variation, scGCO and spatialDE demonstrated robust performance and identified spatial genes in all 21 samples, while trendSceek only identified spatial genes in 15 samples (Fig. 2f, Supplementary Fig. 12, 13). Moreover, scGCO consistently identified more spatial genes than spatialDE in 15 out of 21 samples, and outnumbered trendSceek in 14 out of 21 samples, further demonstrating scGCO’s superior performance.

Finally, we extended the analysis to MERFISH data^3^. ScGCO identified 139 spatial genes, which is comparable to trendSceek (140) and is higher than spatialDE (91). Interestingly, spatial genes identified by the three methods displayed a near perfect overlap, supporting a comparable performance (Supplementary Fig. 14). However, only 150 genes were identified by all three methods combined. Hence, the similarity is likely to be a consequence of a lack of spatial genes, rather than a valid indicator of the algorithms’ performance.

### scGCO scales linearly with the number of cells

Spatial gene expression is now being measured for millions of cells^10^; hence, it is essential that analysis methods demonstrate scalabilities that meet these challenges. We first compared the memory requirement of scGCO, spatialDE and trendSceek using simulated data with cell numbers up to a million. Consistent with previous algorithm analyses results^6,7^, memory footprints of spatialDE and trendSceek grow quadratically with the number of cells. Importantly, both algorithms are unpractical to scale to 1 million cells, because they require about 8 T and 106 T memory, respectively (Fig. 2g). In contrast, scGCO demonstrates a minimal memory requirement that grows near linearly with the number of cells and can process 1 million cells using only 19 GB memory (Fig. 2g). The low memory footprint of scGCO is expected because scGCO uses a graphical representation of spatial information of cells that is intrinsically sparse, because cells only make contact with a few neighboring cells.

We next compared the running times of scGCO, spatialDE and trendSceek using the same simulated data. For cell numbers less than 5,000, scGCO and spatialDE deliver excellent running time and can perform analysis in minutes using a typical desktop computer (Fig. 2h). TrendSceek is not competitive and requires orders of magnitude longer running time under the same test conditions (Fig. 2h). Importantly, the running time of spatialDE and trendSceek is cubic or quadratic in the number of cells^6,7^, and both methods are unpractical to scale to millions of cells (Fig. 2h). In sharp contrast, scGCO’s running time is linear in the number of cells, which is consistent with benchmarks of graph cuts^11^. As a result, scGCO can analyze 1,000,000 cells in less than 3 hours using a typical desktop computer (Fig. 2h), demonstrating unparalleled scalability.

## Discussion

Single-cell sequencing technology is enjoying a rapid revolution, and data are now being generated for millions of cells in a single experiment^10^. This astronomical amount of data poses a great challenge for analysis methods, which are essential to fully realize values for single-cell data. By employing powerful graph cuts algorithms for spatial gene analysis, our method delivers excellent scalability and can process millions of cells in a reasonable time using modest hardware. Moreover, the graph cuts algorithm has demonstrated excellent performance in 3-D object recognition^12^ and can be accelerated by GPU^13^. Hence, our method could readily scale to 3-D single-cell spatial transcriptomics data.

By posing spatial gene identification as an image-processing problem, our method delivers a powerful visual presentation of identified spatial patterns and could be valuable for a broad spectrum of researchers. In contrast to existing methods, graph cuts do not rely on assumptions of data distribution and theoretically can identify any pattern of spatial distribution. Moreover, while the hyperparameters in Gaussian process or marked point process typically settle for a local optimum, for bilabel image segmentation, which is equivalent to identifying spatial genes that are over-or underexpressed in specific regions, graph cuts guarantee to find the global optimal solution^14,15^. Consequently, our method consistently demonstrates superior performance across a wide range of spatial gene expression data types. Taken together, we expect scGCO to become the method of reference for spatial gene expression analyses.

## Materials and methods

### Graph and Voronoi diagram representation of spatial gene expression data

To apply the graph cuts algorithm to spatial gene expression data, we first performed Delaunay triangulation on the spatial coordinates of the cells. The graph produced by Delaunay triangulation has the nice property that only authentic neighbors are connected by edges in the graph because no cells are allowed in the triangle connecting three cells. Hence, Delaunay triangulation captures essential information of cell-cell interactions with a sparse graph. After spatial gene expression patterns have been identified by graph cuts, we performed the dual operation of Delaunay triangulation to generate Voronoi diagrams, which has been broadly used to model cells^16^. To highlight the boundaries of cell clusters identified by graph cuts, edges in the Delaunay triangulation connecting cells with different predicted labels are identified, and Voronoi polygon edges intersecting these identified edges in Delaunay triangulation are highlighted, providing a direct visual representation of spatial gene expression patterns.

### Markov random fields model

A Markov random field (MRF) is an undirected graphical model capturing conditional independence among a set of random variables. According to the Hammersley-Clifford Theorem, the joint distribution *p*(*X*) of an MRF can be written as a product of positive potential functions *ψ_c_*(*x_c_*) over the maximal cliques of the graph:

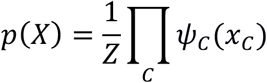

where *Z* is the partition function that normalizes the distribution *p*(*X*), which is the sum of potential functions over all maximal cliques. The positive potential functions allow the joint distribution of an MRF to be conveniently written as a Gibbs distribution:

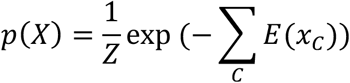

where *E*(*x_c_*) > 0 is the energy associated with the variables in clique *c*. Thus, minimizing the total energy function is equivalent to the maximum a posteriori estimation of *p*(*X*).

Studies analyzing spatial expression of genes demonstrated that the spatial distribution of expression values forms patches, where adjacent cells tend to display comparable levels of gene expression^4^. Thus, patches of cells in which a gene displays similar gene expression levels are analogous to objects in an image. Consequently, we adopt the classical energy formulation for image segmentation in computer vision to describe the spatial distribution of gene expression in single cells:

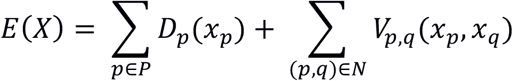

where *N* is the set of adjacent cells that interact directly in the graphical representation of single cell spatial gene expression data. In the context of single cell spatial gene expression analysis, *D_p_*(*x_p_*) is a data penalty function of assigning a particular gene expression classification *x* to cell *p*, and *V_p,q_*(*x_p_, x_q_*) is the interaction energy of assigning a particular pair of gene expression classifications to a pair of cells interacting directly. Essentially, assigning gene expression classifications is analogous to assigning pixel labels in image segmentation. Although *V_p,q_*(*x_p_, x_q_*) can take many forms, a common requirement is that the interaction energy penalizes the assignment that adjacent cells are with different classifications, which is crucial to identify patches of cells with similar gene expression patterns.

### Minimizing MRFs energy with graph cuts

When the classification of cells is limited to two classes, or two labels in an image segmentation problem, a crucial advantage of the above energy formulation of MRFs is that powerful min-cut/max-flow algorithms for graph cuts can be used to minimize the above energy functions, which provides fast, globally optimal solutions for two-label problems^15^. For multilabel problems, global minimization of the energy function is NP-hard^17^. In scGCO, we adopt the alpha-expansion algorithm developed by Boykov et al., which iteratively applies 2-label graph cuts to expand each label until the algorithm converges^17^. The algorithm runs in low polynomial time and guarantees that the solution is within a known factor of the global minimum^17^.

The above graph cuts algorithm can be applied to energy minimization of MRFs if and only if the interaction energy is regular^18^:

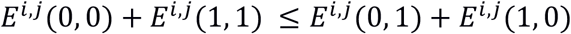

The regularity of interaction energy guarantees a duality between energy states of MRFs and label configurations of the corresponding graph, where the minimal energy state matches the maximum flow of the graph, hence allowing the application of graph cuts to solve energy minimization of MRFs. In our implementation, we used a topological interaction energy that has greater penalties when the classification of adjacent cells is further away. Specifically, the interaction energy *S* is a symmetric matrix whose entries were:

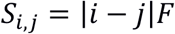

where *F* is a smooth factor that controls the size of the penalty and *S_i,j_* is the interaction energy for adjacent cells with classification *i* and *j* respectively.

### Statistical significance of identified spatial genes

We modeled the spatial gene expression patterns as homogeneous spatial Poisson processes, which describe the random distribution of points in 2-D plane. For points with a density *ρ*, the probability of finding exactly *k* such points in a region *V* can be determined from Poisson distribution:

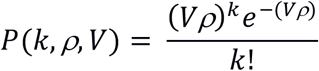

In the setting of spatial gene expression analysis, the graph cuts algorithm will separate cells into distinct segments according to gene expression classification predicted by the MRF model. *V* is the number of cells in a segment determined by graph cuts. Although all cells in the same segment have the same predicted classification, the cells’ true classifications determined from their gene expression levels may be different. In the analyzed segment, *k* is the number of cells with a particular true classification, and *ρ* is the density of cells of corresponding true classification in the entire sample. For each candidate gene, we analyzed all possible classifications (all *k*, *p* pairs) in all segments identified by graph cuts, and reported the best result as the p-value for the gene. For genome-scale analyses, multiple test correction was performed with Benjamini– Hochberg procedure.

### Gene expression classification via Gaussian mixture modeling

For each gene we performed Gaussian mixture modeling (GMM) on its gene expression vector to identify the underlying Gaussian distribution components. We then assigned each cell a gene expression classification according to the GMM classification of the gene’s expression level in the cell. The classifications were ordered by corresponding gene expression levels so that cells with larger difference in gene expression levels have greater difference in their classifications. This setup ensures that adjacent cells with larger expression difference are associated with larger classification differences, which will generate larger penalties in energies of associated MRFs. This energy formulation favors graph cuts that put cells with similar classifications in the same sub-graph.

To determine the best number of components for GMM, we generated GMM with component numbers from 2 to 10. We then calculated Bayesian information criterion (BIC) for each GMM and selected the GMM with best BIC as final GMM for downstream analysis.

### Data sets and data preprocessing

We downloaded the spatial transcriptomics data reported by Ståhl et al. from the Spatial Transcriptomics Research website (http://www.spatialtranscriptomicsresearch.org/datasets/doi-10-1126science-aaf2403)^4^. We used all 12 replicates for the mouse olfactory bulb, and all four layers for the breast cancer data. For mouse hippocampus seqFISH data^2^, we downloaded the data from https://ars.els-cdn.com/content/image/1-s2.0-S0896627316307024-mmc6.xlsx. We used all 21 fields provided by the authors for analysis. The MERFISH data was downloaded from the Zhuang lab website (http://zhuang.harvard.edu/MERFISHData/data_for_release.zip)^3^. We used “Replicate 6” similar to spatialDE^6^, as these had the largest number of cells and highest confluency. Expression data were normalized using the same procedure as described in the cellranger package (https://support.10xgenomics.com/single-cell-gene-expression/software/pipelines/latest/what-is-cell-ranger).

### Comparison to existing spatial gene identification algorithms

To systematically evaluate the performance of scGCO against two published algorithms (spatialDE and trendSceek), we ran spatialDE, trendSceek and scGCO on all the samples in mouse olfactory bulb data (12 replicates), the breast cancer data (4 samples), mouse hippocampus seqFISH data (33 samples), and MERFISH dataset (1 sample). For spatialDE, we downloaded the scripts provided by the authors from their GitHub website and executed the scripts without modification. For trendSceek, we implemented R scripts according to the methods descripted in trendSceek’s original paper. The trendSceek’s scripts and the scripts to run scGCO are provided in the tutorial files in scGCO’s GitHub repository.

To estimate the scalabilities of algorithms, we evaluated memory requirement and running time using simulated data as described by Edsgard et al.^7^. For running time, we executed all algorithms on a desktop computer with Intel^®^ Core™ i7-6700 CPU (8 cores at 3.40GHz), 40 GiB memory, and running the Ubuntu 18.04.1 operating system. For memory profiling, we executed all algorithms on a work station with 2 TB of memory. For spatialDE and trendSceek, both algorithms exceed the capacity of available hardware when the cell numbers are large. Because these algorithms scale quadratically or cubically with the number of cells^6,7^, we estimated their memory requirement and running time by fitting available data to polynomial functions.

### Gene ontology and network analyses

The gene set enrichment analyses were carried out with GSEA^19^ desktop version 3.0 with number of permutations set to 1000, max size (exclude large size) set to 500 and min size (exclude smaller size) set to 15. Gene Ontology analyses were carried out with R package clusterProfiler^20^ using default parameters. The GO enrichment graph was generated with Cytoscape^21^ (version 3.6.1) plugins ClueGO^22^ version 2.5.2 and CluePedia^23^ version 1.5.2 using a kappa score cutoff of 0.6.

## Supporting information

## Code availability

An open source implementation of scGCO is available at GitHub (https://github.com/WangPeng-Lab/scGCO).

## Acknowledgements

This work was supported in part by the National Key R&D Program of China grant 2017YFC0907505, 2017YFC1201200, 2016YFC0901904, and National Natural Science Foundation of China (NSFC) grant 31671380.

## Author Contributions

K.Z. and W.F. implemented the software and performed experiments. P.W. designed the algorithm, supervised the study and implemented the software. P.W. wrote the manuscript with inputs from all authors. All authors approved the final manuscript.

