## Supplementary material for "Identification of spatially variable genes with graph cuts"

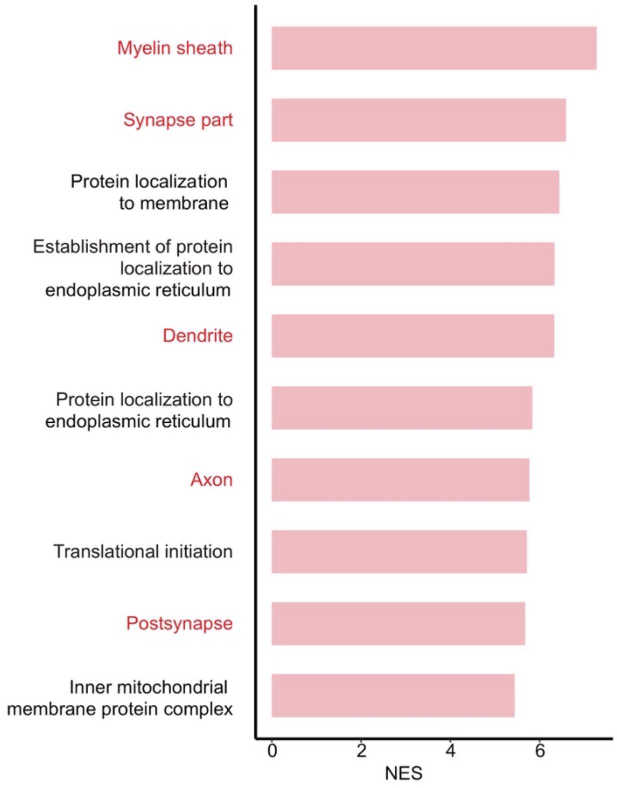


**Supplementary Figure 1.** Genes identified by scGCO from mouse olfactory bulb replicate 11 are enriched with neuron-related functions.

Barcharts showing the top ten significant gene sets identified by GSEA analysis. Functional categories associated with neuron are highlighted in red.

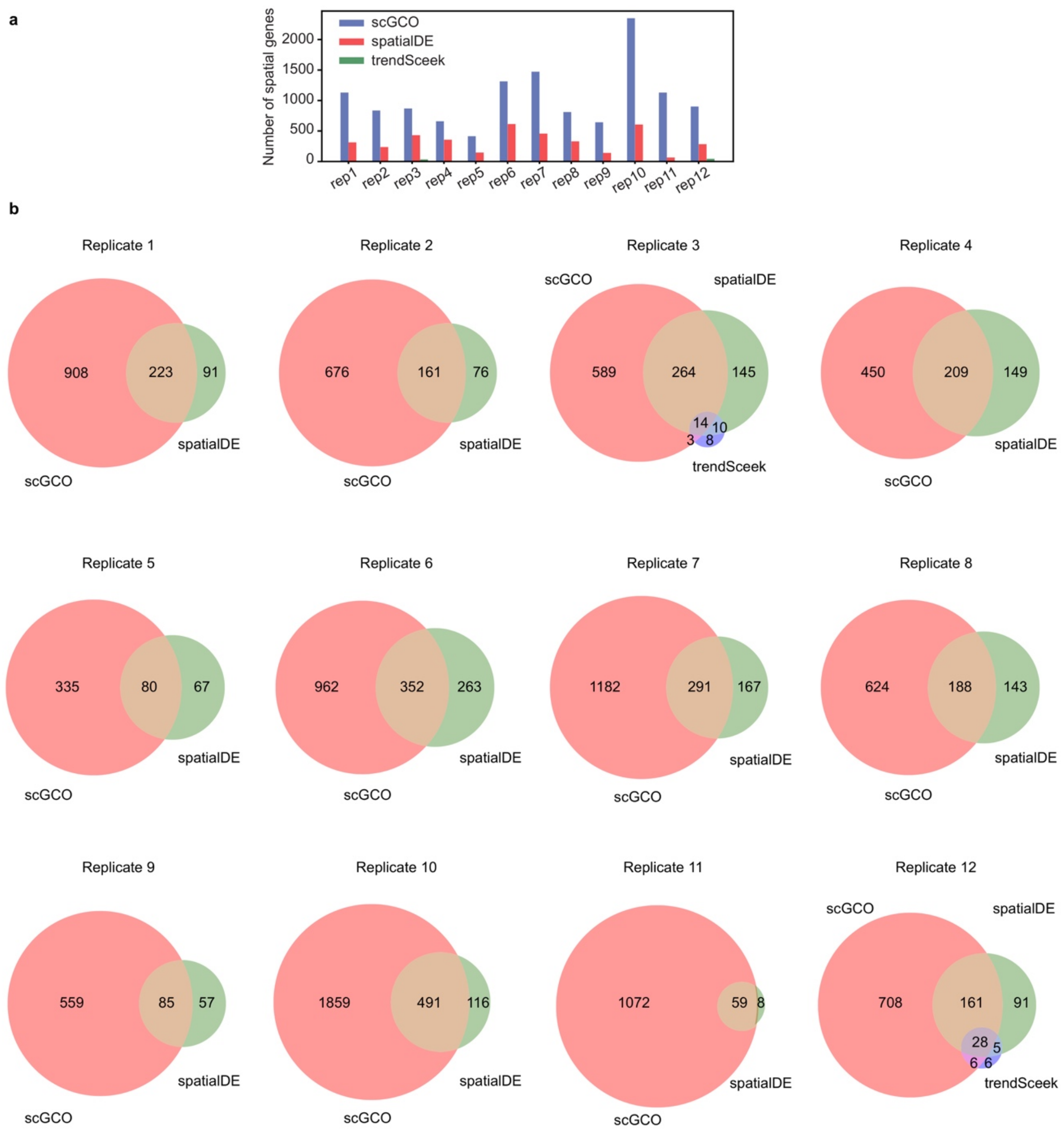


**Supplementary Figure 2.** ScGCO outperformed spatialDE and trendSceek in mouse olfactory bulb data.

(**a**) Barcharts showing the number of spatially variable genes identified by scGCO, spatialDE and trendSceek for all 12 replicates of mouse olfactory bulb data. (**b**) Venn diagrams showing the set relationship among spatial genes identified by scGCO, spatialDE and trendSceek for mouse olfactory bulb data.

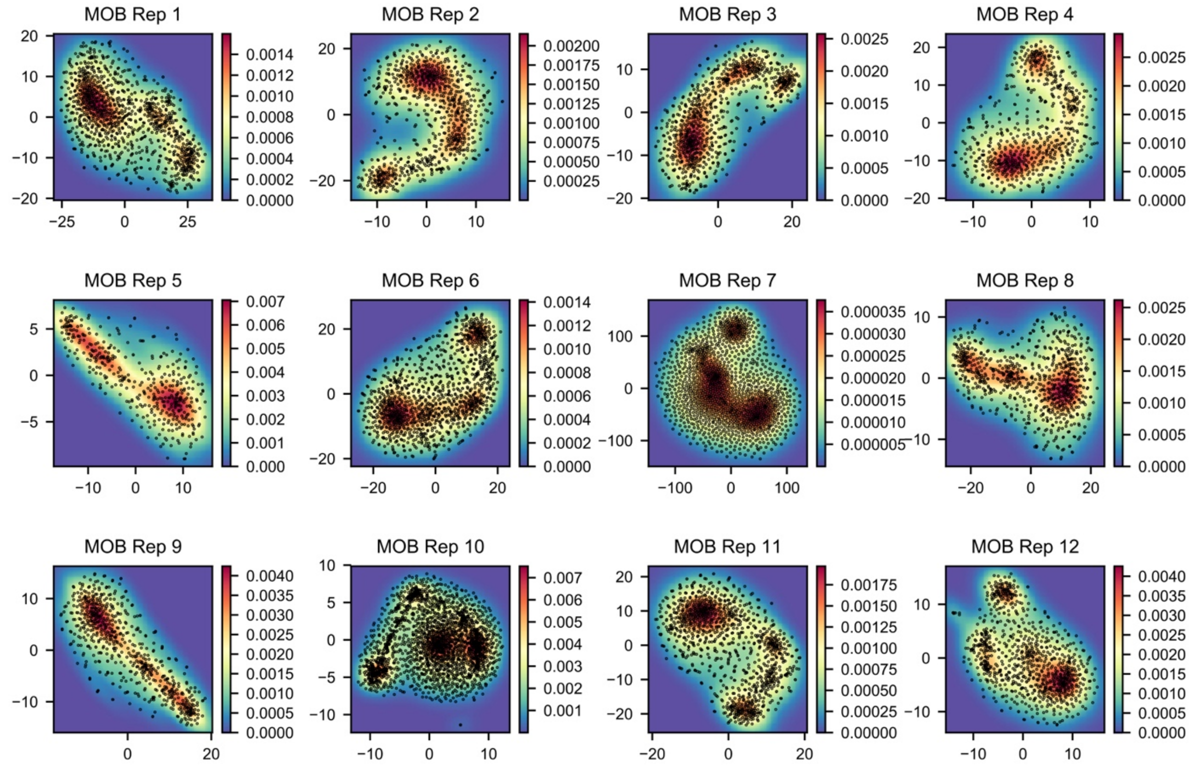


**Supplementary Figure 3.** Cluster analyses of spatially variable genes identified by scGCO for mouse olfactory bulb data.

T-SNE analyses of spatially variable genes identified by scGCO for all 12 replicates of mouse olfactory bulb data. Each point is a gene. Background color indicates density of points determined with kernel density estimation.

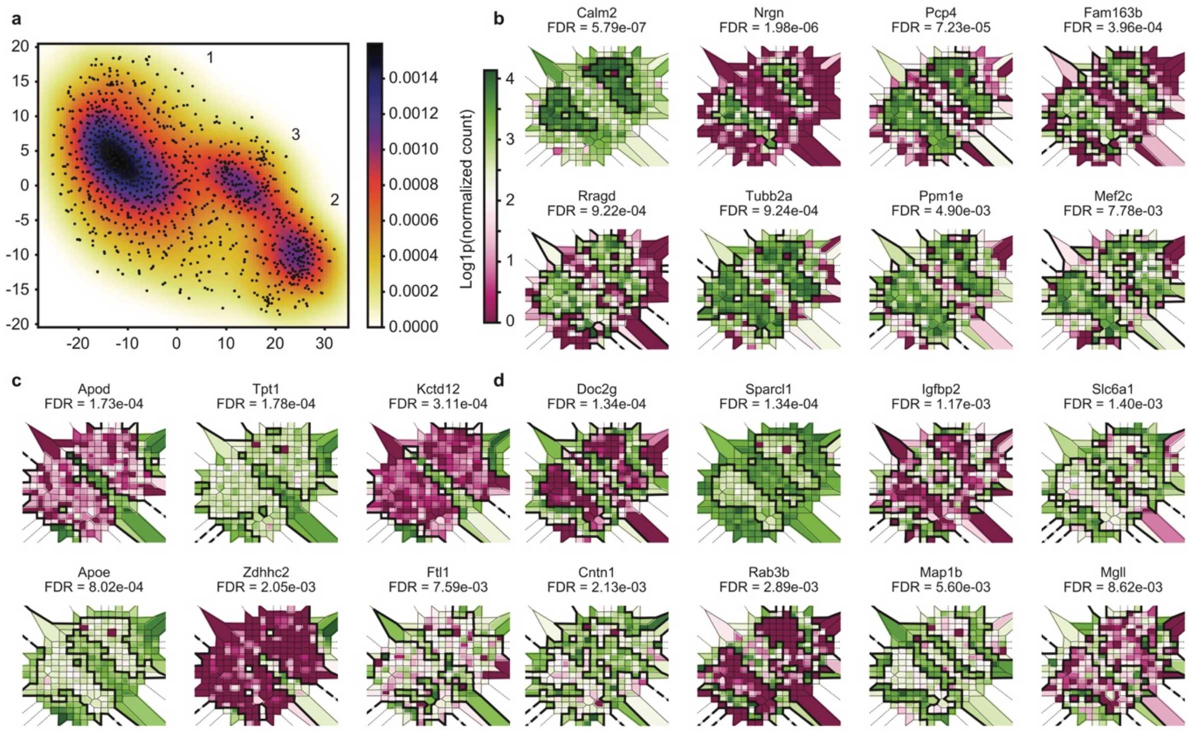


**Supplementary Figure 4.** ScGCO identified spatially variable genes in replicate 1 of mouse olfactory bulb data.

(**a**) T-SNE analyses of spatially variable genes identified by scGCO for replicate 1 of mouse olfactory bulb data. Each point is a gene. Background color indicates density of points determined by kernel density estimation. Numbers in the graph indicate index of clusters. (**b**) Voronoi diagrams showing representative spatial gene expression patterns in cluster 1. Polygon representing each cell is colored according to the associated gene expression level. The boundaries for spatial expression patterns are depicted by thicker black lines. (**c**) same as b for cluster 2. (**d**) same as b for cluster 3.

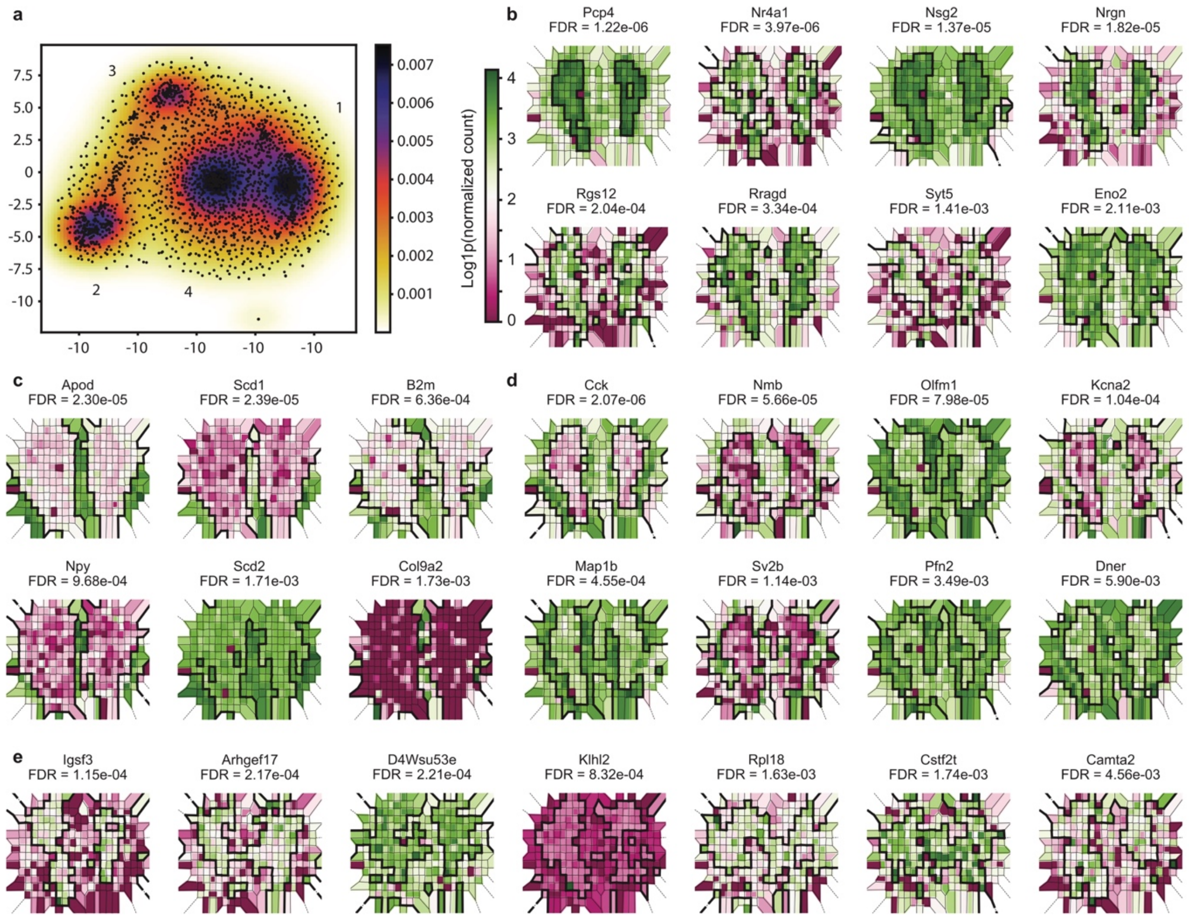


**Supplementary Figure 5.** ScGCO identified spatially variable genes in replicate 10 of mouse olfactory bulb data.

(**a**) T-SNE analyses of spatially variable genes identified by scGCO for replicate 10 of mouse olfactory bulb data. Each point is a gene. Background color indicates density of points determined by kernel density estimation. Numbers in the graph indicate index of clusters. (**b**) Voronoi diagrams showing representative spatial gene expression patterns in cluster 1. Polygon representing each cell is colored according to the associated gene expression level. The boundaries for spatial expression patterns are depicted by thicker black lines. (**c**) same as b for cluster 2. (**d**) same as b for cluster 3. (**e**) same as b for cluster 4.

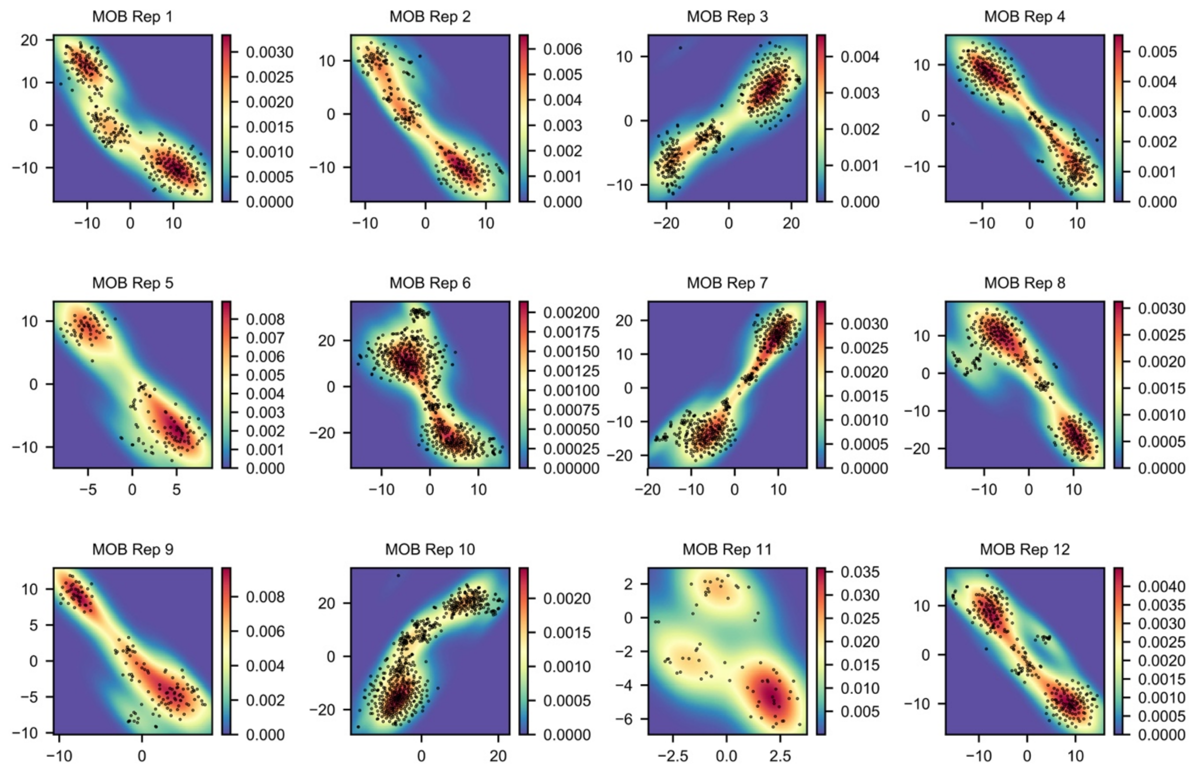


**Supplementary Figure 6.** Cluster analyses of spatially variable genes identified by spatialDE for mouse olfactory bulb data.

T-SNE analyses of spatially variable genes identified by spatialDE for all 12 replicates of mouse olfactory bulb data. Each point is a gene. Background color indicates density of points determined by kernel density estimation.

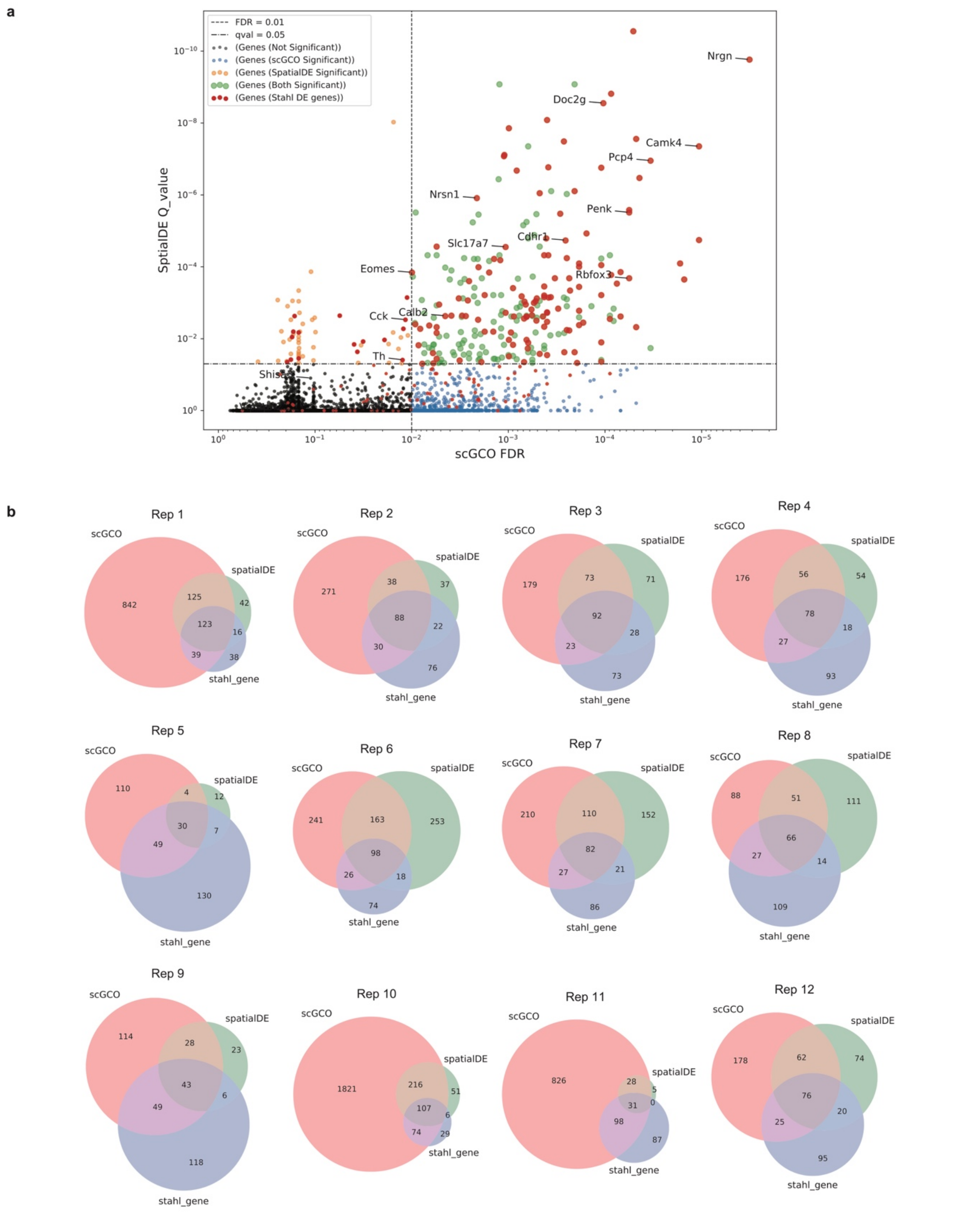


**Supplementary Figure 7.** ScGCO outperformed spatialDE by recovering more spatial genes identified by direct comparison of GCL v.s. GL region of mouse olfactory bulb.

(a) A representative scatter plot showing fraction of spatial genes identified by direct comparing GCL v.s. GL recovered by scGCO versus spatialDE (replicate 1). The dashed horizontal line indicates the significance cutoff for spatialDE (FDR < 0.05). The dashed vertical line indicates the significance cutoff for scGCO (FDR < 0.01). All genes in the mouse olfactory bulb data are shown. Gene names for example known marker genes reported by Ståhl et al. are shown. (**b**) Venn diagrams showing the set relationship among spatial genes identified by scGCO, spatialDE and by direct comparison between GCL and GL layer for mouse olfactory bulb data as described by Ståhl et al..

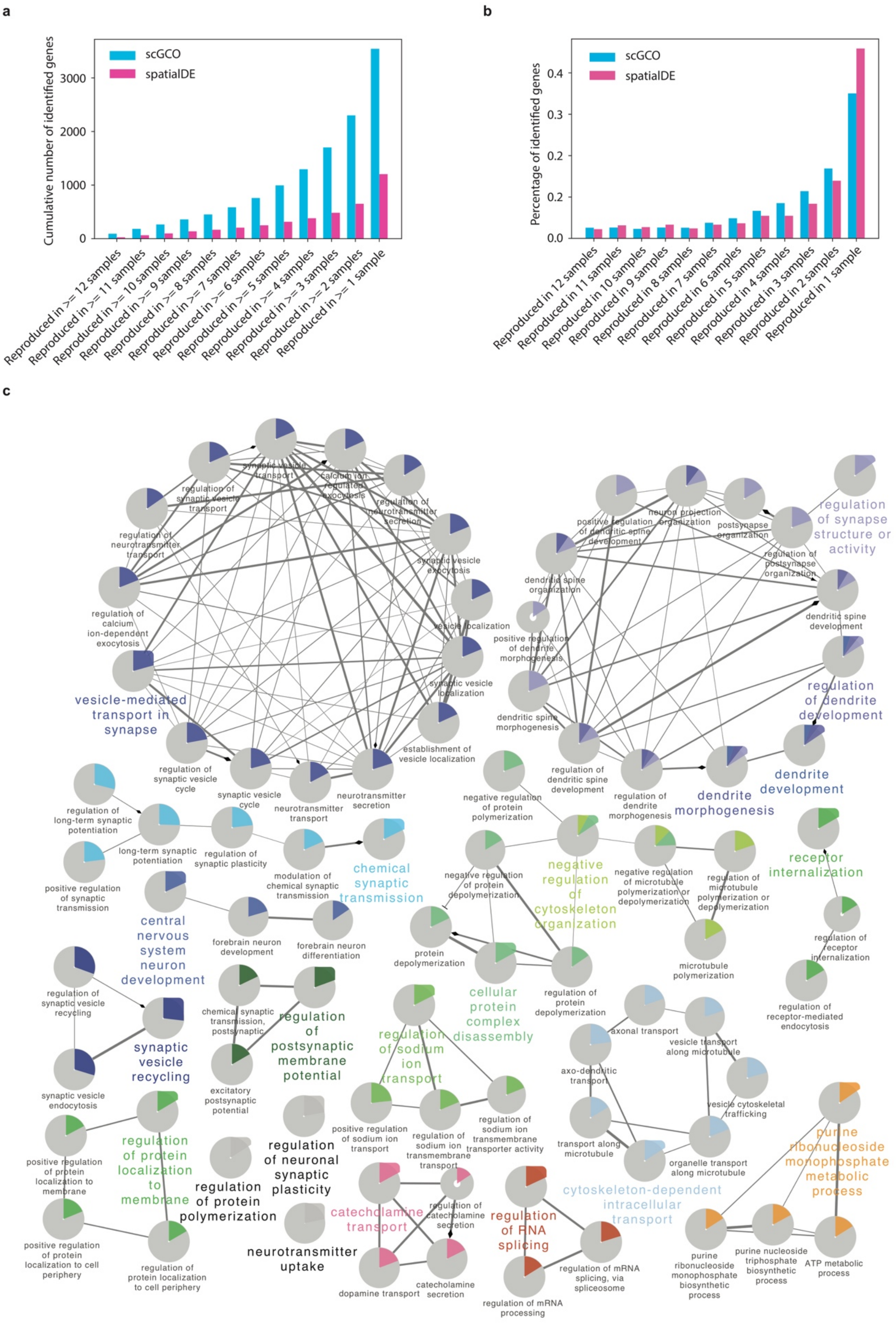
a
b
c
Cumulative number of identified genes
scGCO
spatialDE
Reproduced in >= 12 samples
Reproduced in >= 11 samples
Reproduced in >= 10 samples
Reproduced in >= 9 samples
Reproduced in >= 8 samples
Reproduced in >= 7 samples
Reproduced in >= 6 samples
Reproduced in >= 5 samples
Reproduced in >= 4 samples
Reproduced in >= 3 samples
Reproduced in >= 2 samples
Reproduced in >= 1 sample
Percentage of identified genes
Reproduced in 12 samples
Reproduced in 11 samples
Reproduced in 10 samples
Reproduced in 9 samples
Reproduced in 8 samples
Reproduced in 7 samples
Reproduced in 6 samples
Reproduced in 5 samples
Reproduced in 4 samples
Reproduced in 3 samples
Reproduced in 2 samples
Reproduced in 1 sample
vesicle-mediated transport in synapse
regulation of synapse structure or activity
regulation of dendrite development
dendrite development
dendrite morphogenesis
chemical synaptic transmission
central nervous system neuron development
negative regulation of cytoskeleton organization
receptor internalization
synaptic vesicle recycling
regulation of postsynaptic membrane potential
cellular protein complex disassembly
regulation of sodium ion transport
regulation of protein localization to membrane
regulation of neuronal synaptic plasticity
regulation of protein polymerization
neurotransmitter uptake
catecholamine transport
regulation of RNA splicing
cytoskeleton-dependent intracellular transport
purine ribonucleoside monophosphate metabolic process

**Supplementary Figure 8.** Reproducibility of spatial gene identification algorithms in mouse olfactory bulb data.

(**a**) Barchars showing number of genes reproducibly identified in number of biological replicates for scGCO and spatialDE. (**b**) Barchars showing percentile of genes reproducibly identified in number of biological replicates. (**c**) Network of enriched GO terms in genes reproducibly identified by scGCO (reproducibly identified in >= 6 samples). Related GO terms are connected and labels summarizing their common function are shown in larger font. Colored parts in the pie chart indicate fraction of genes in the GO term that are scGCO identified genes.

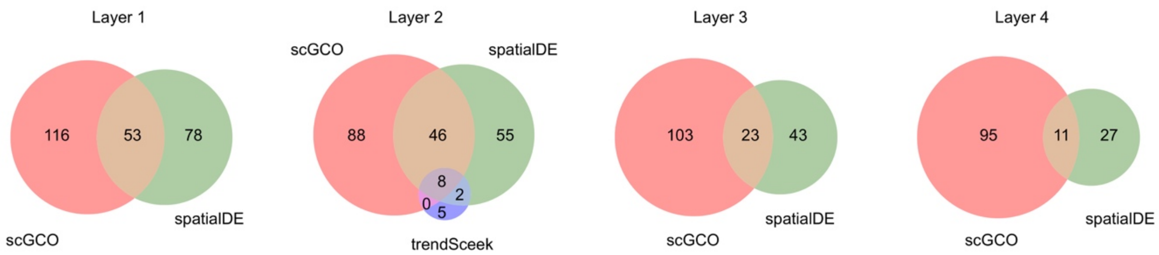


**Supplementary Figure 9.** Comparing scGCO with spatialDE and trendSceek using breast cancer biopsies data.

Venn diagrams showing the set relationship among spatial genes identified by scGCO, spatialDE and trendSceek for breast cancer biopsies (FDR < 0.05).

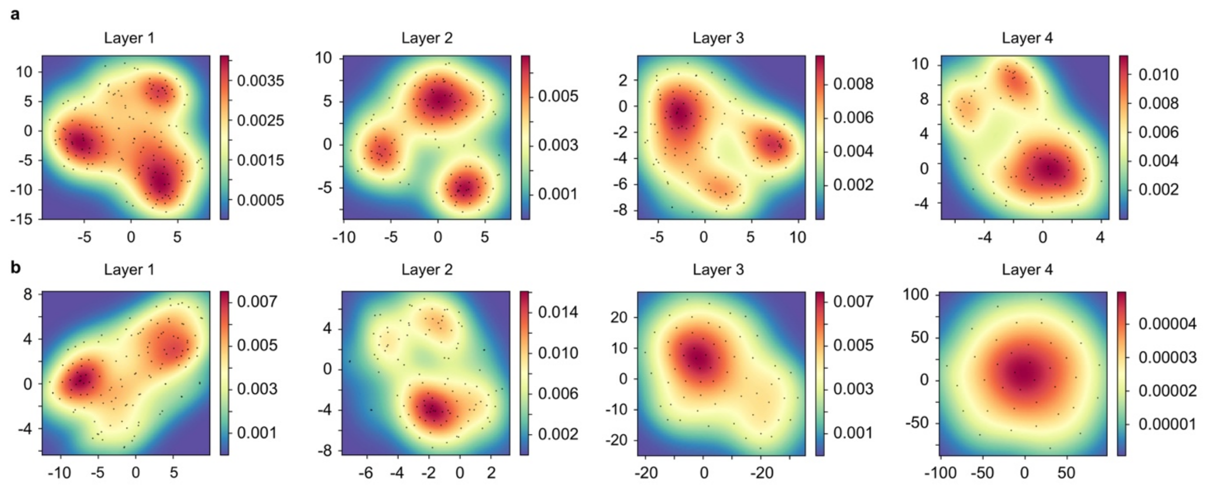


**Supplementary Figure 10.** Spatial genes identified by scGCO demonstrates better robustness than spatialDE in breast cancer biopsies data.

(**a**) t-SNE analysis of spatially variable genes identified by scGCO. Each point is a gene. Background color indicates density of points determined by kernel density estimation. (**b**) same as **a** for spatially variable genes identified by spatialDE.

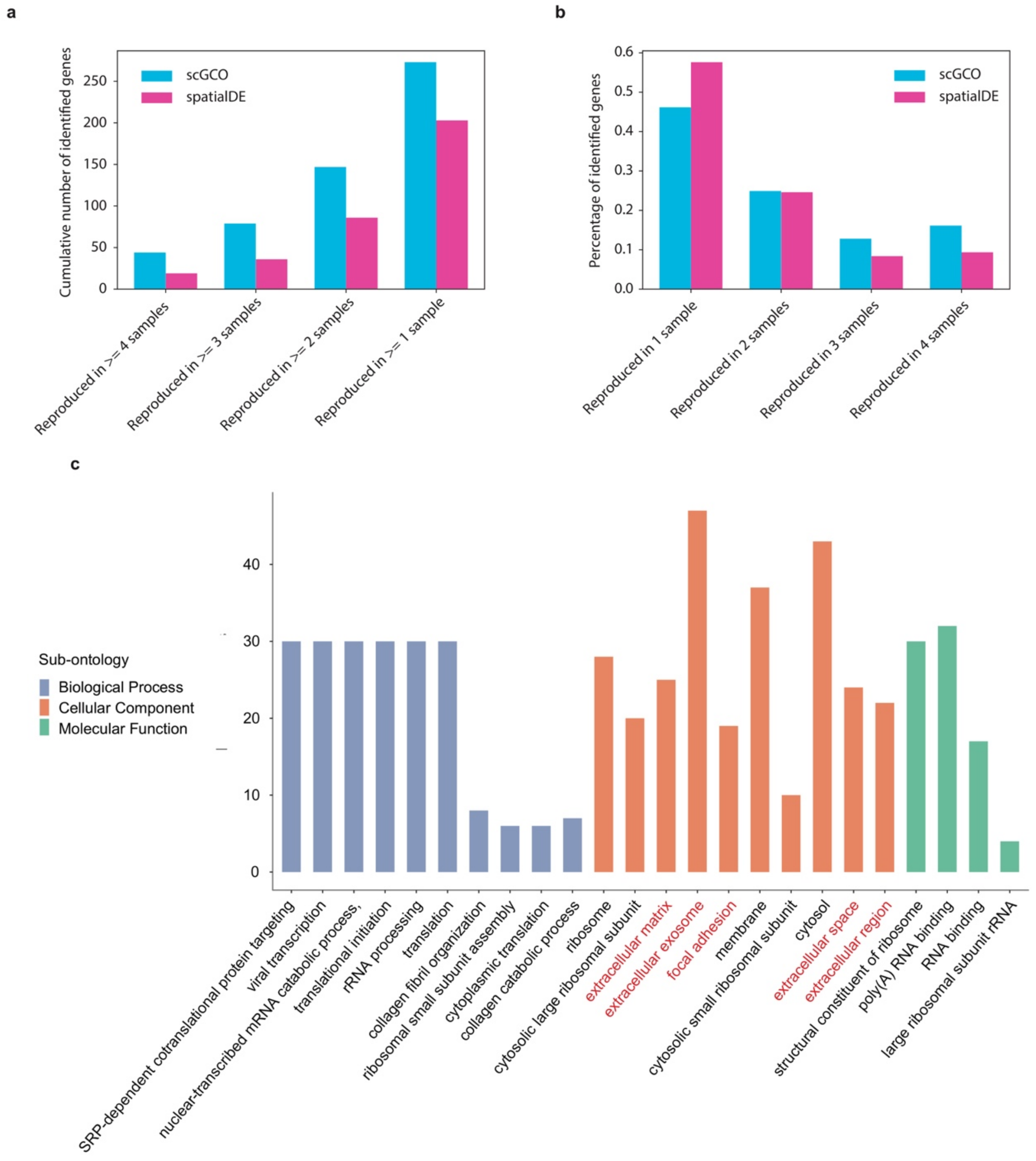


**Supplementary Figure 11.** Reproducibility of spatial gene identification algorithms in breast cancer biopsies data.

(**a**) Barchars showing number of genes reproducibly identified in number of biological replicates for scGCO and spatialDE. (**b**) Barchars showing percentile of genes reproducibly identified in number of biological replicates. (**c**) Barchars showing significant GO terms enriched in genes reproducibly identified by scGCO (reproducibly identified in >= 3 samples). Selected categories potentially related to metastasis (extracellular matrix or focal adhesion) are highlighted in red.

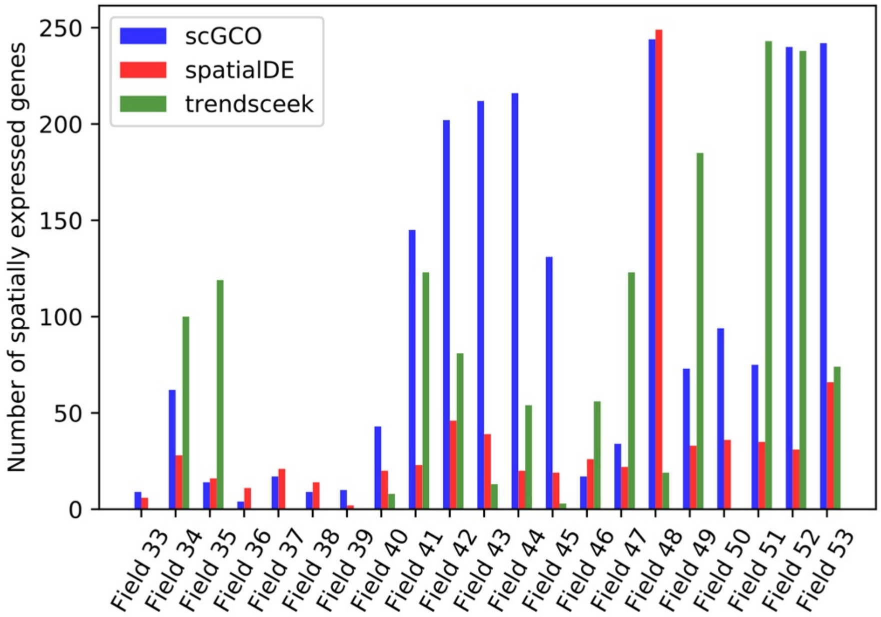


**Supplementary Figure 12.** ScGCO outperformed spatialDE and trendSceek in mouse hippocampus seqFISH data.

Barcharts showing the number of identified spatial genes for scGCO (blue), spatialDE (red) and trendSceek (green) for all 21 fields of mouse hippocampus seqFISH data (FDR < 0.05).

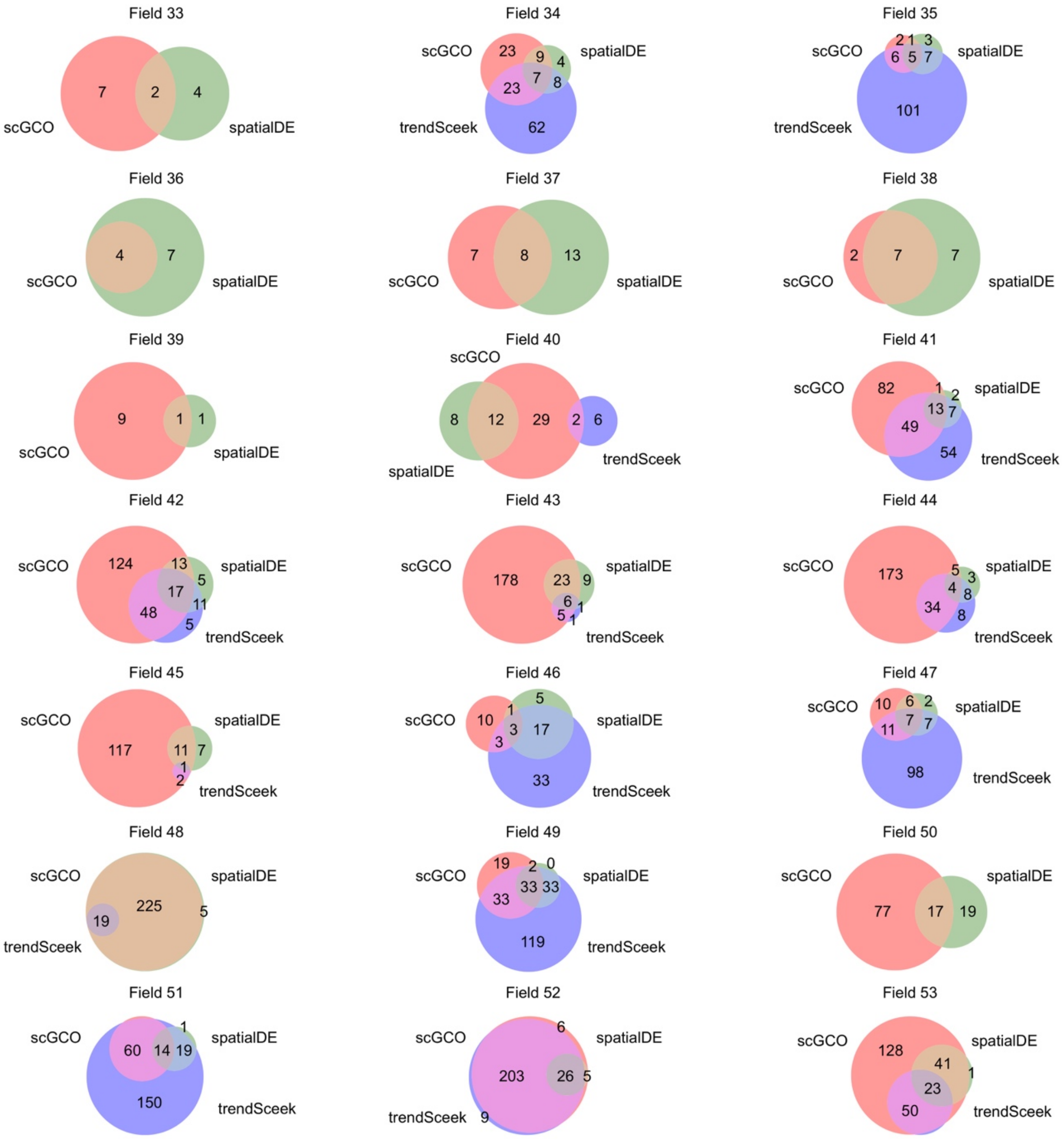


**Supplementary Figure 13.** Comparing scGCO with spatialDE and trendSceek using mouse hippocampus seqFISH data.

Venn diagrams showing the set relationship among spatial genes identified by scGCO, spatialDE and trendSceek for mouse hippocampus seqFISH data (FDR < 0.05).

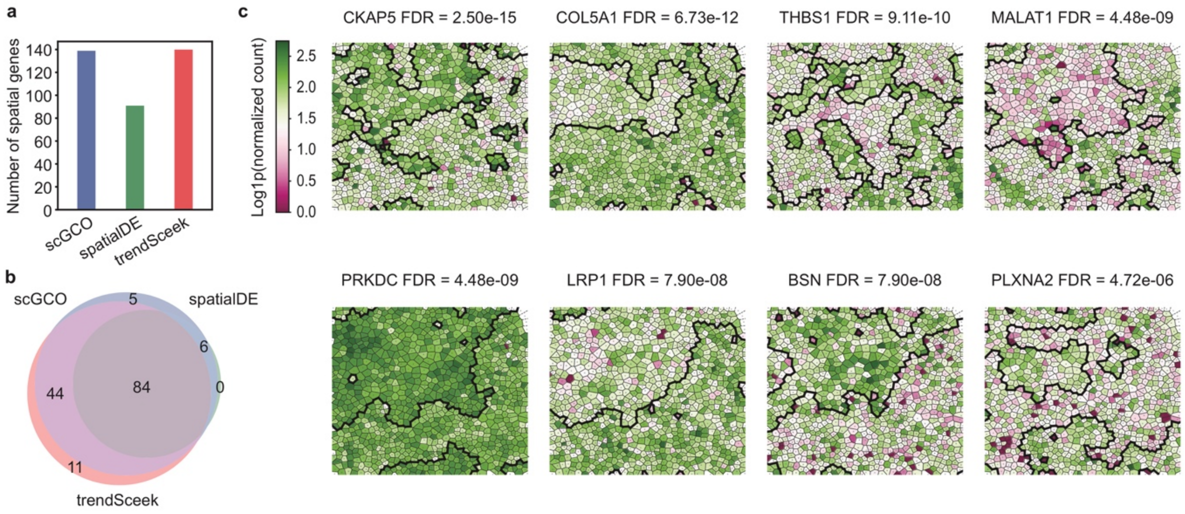


**Supplementary Figure 14**. Identification of spatially variable genes in MERFISH data.

(**a**) Barcharts showing number of spatially variable genes identified by scGCO, spatialDE and trendSceek (FDR < 0.05). (**b**) A Venn diagram showing the set relationship among spatial genes identified by scGCO, spatialDE and trendSceek. (**c**) Representative Voronoi diagrams showing spatial gene expression patterns identified by scGCO in MERFISH data. Polygon representing each cell is colored according to the associated gene expression level. The boundaries for spatial expression patterns are depicted by thicker black lines.
