## Supplementary figures and images for "Identification of spatially variable genes with graph cuts"

### Supplementary file 2

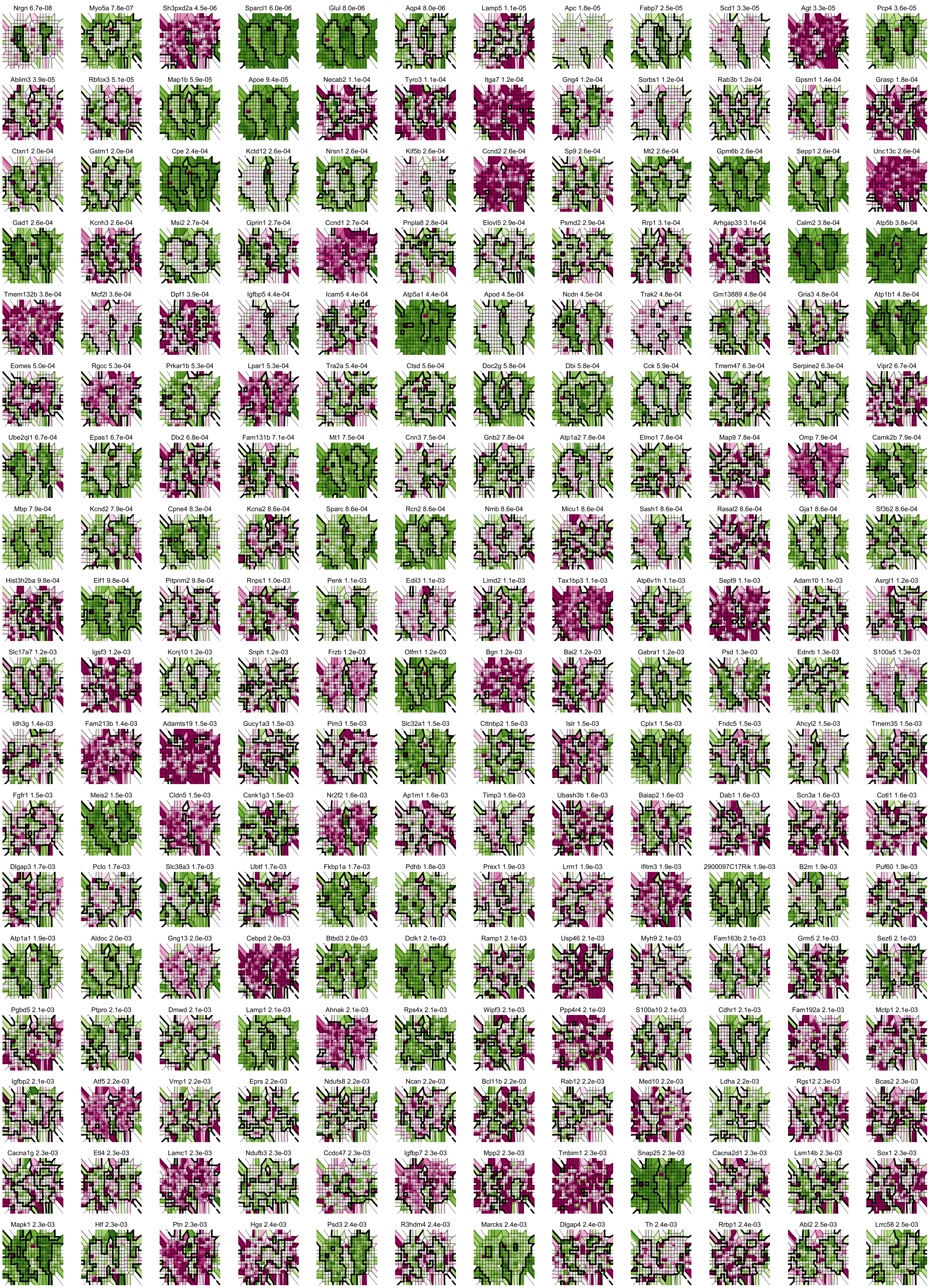

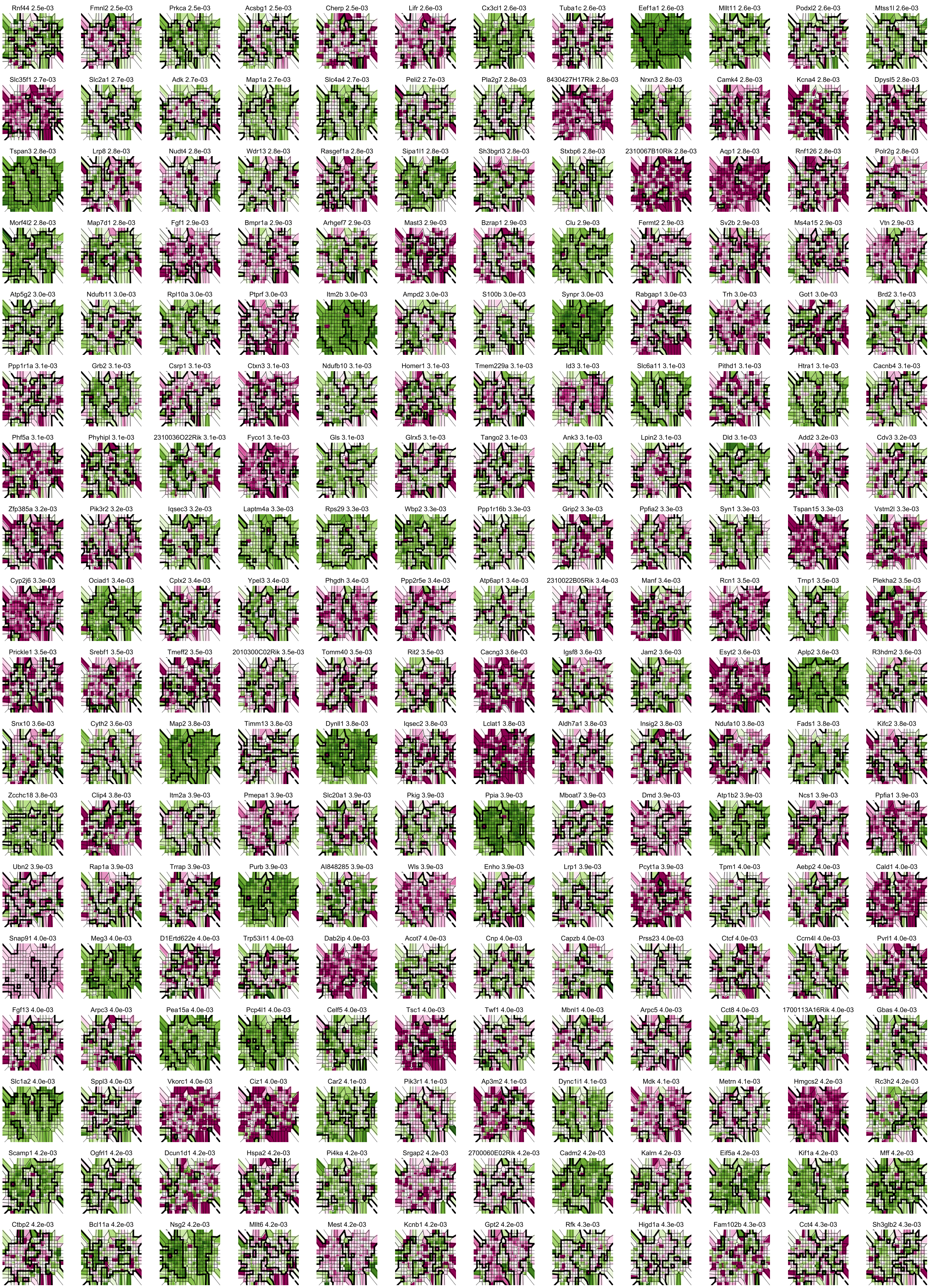

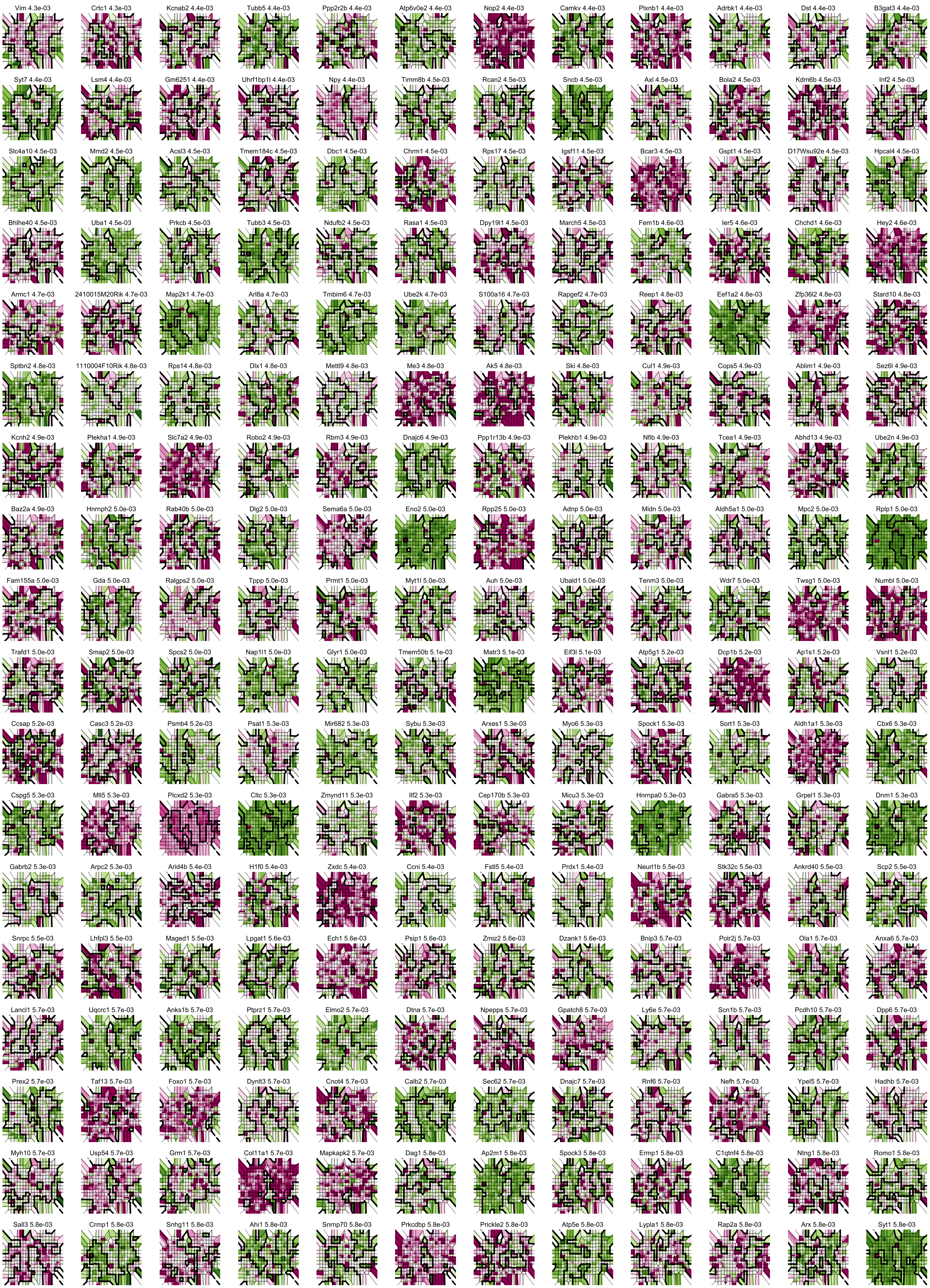

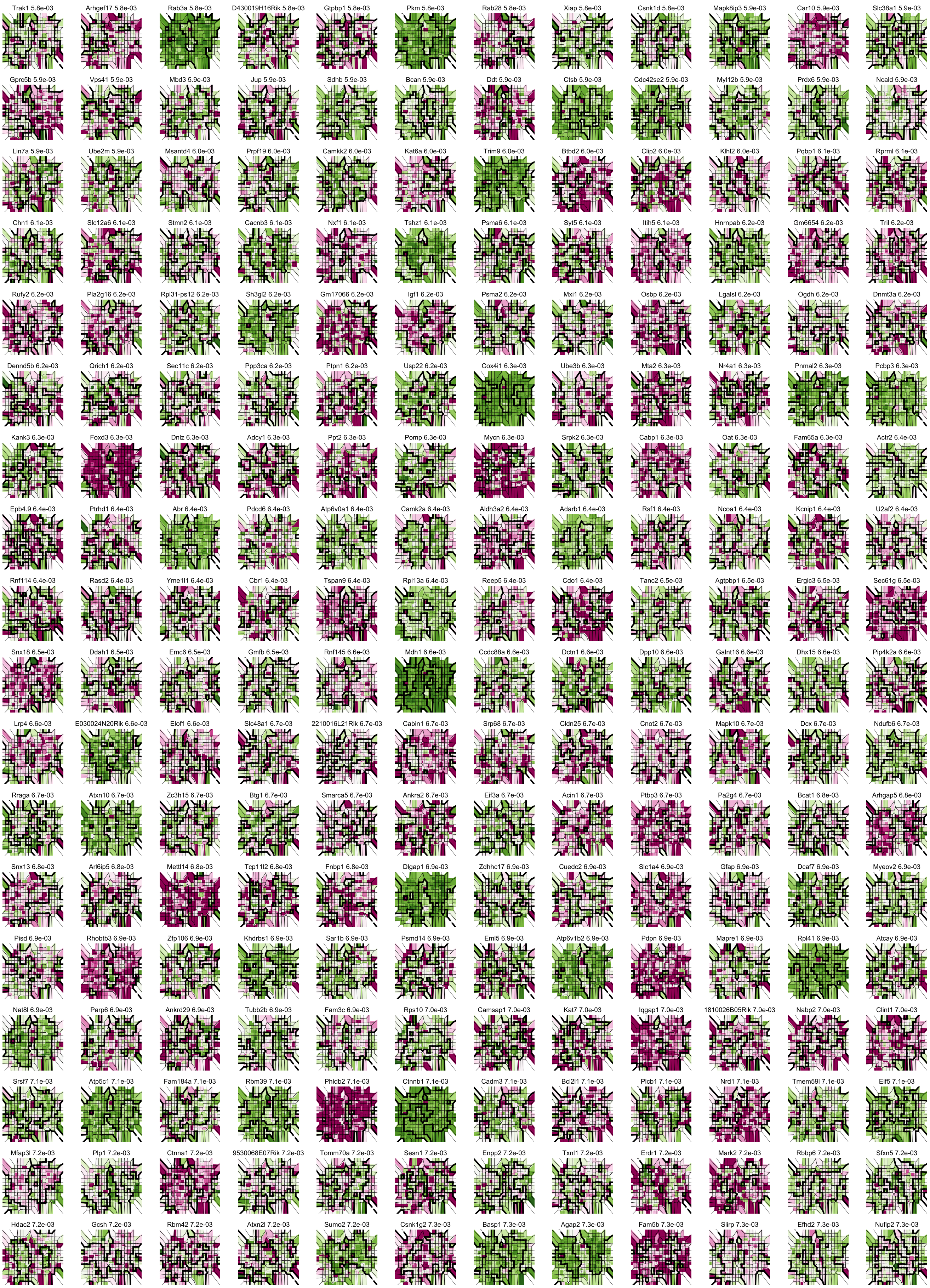

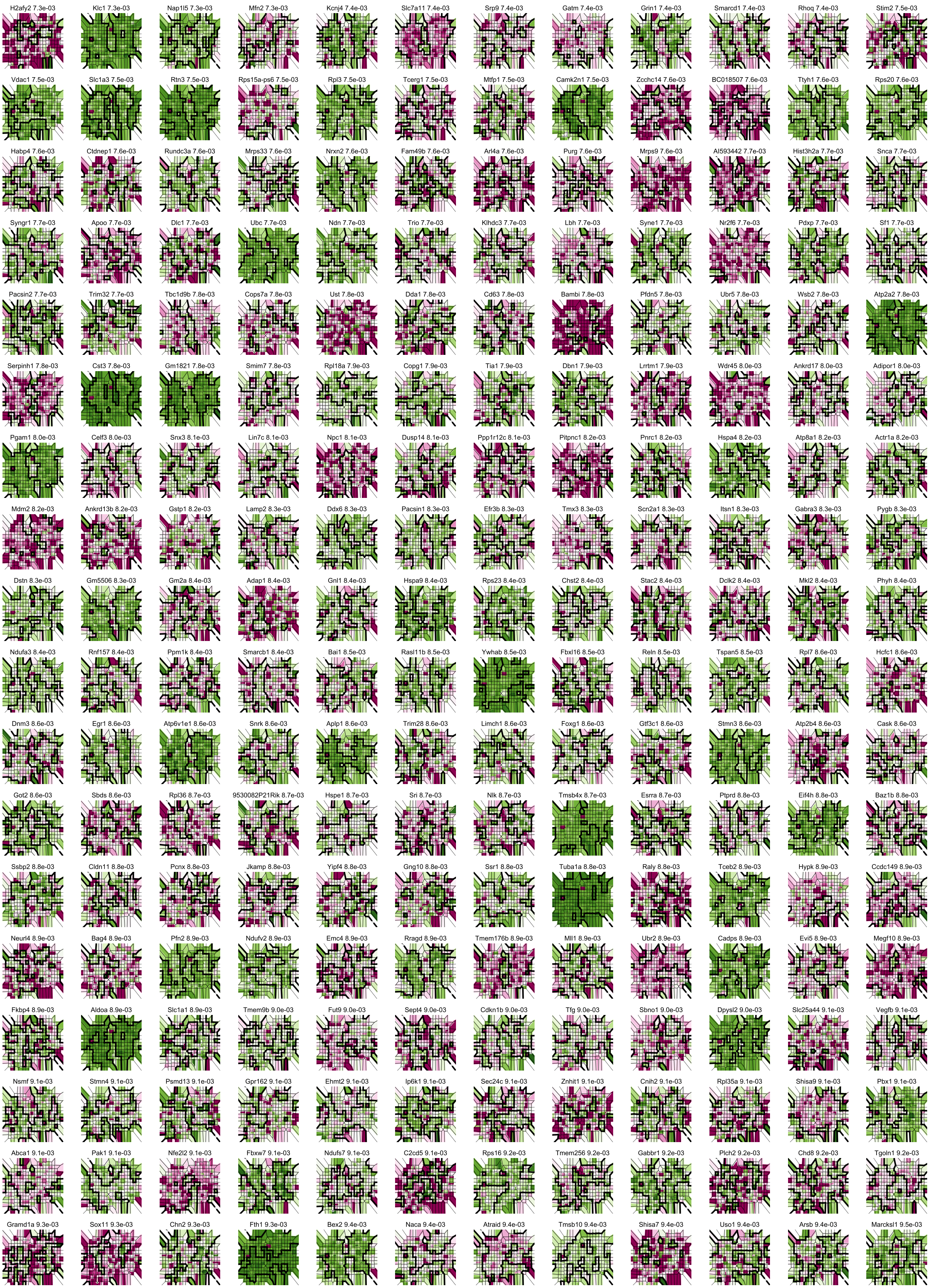

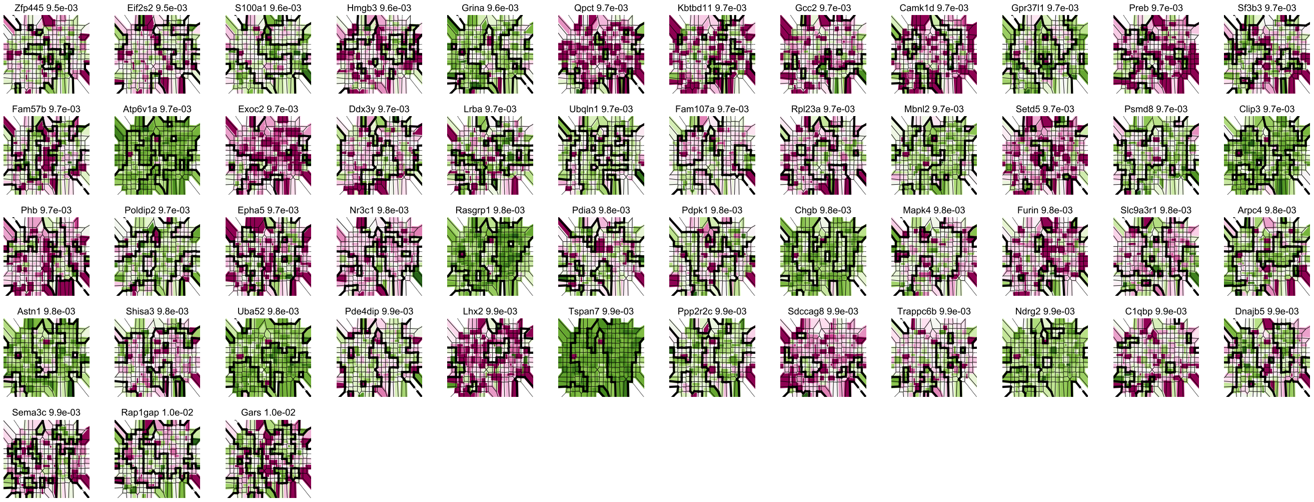

### Supplementary file 3

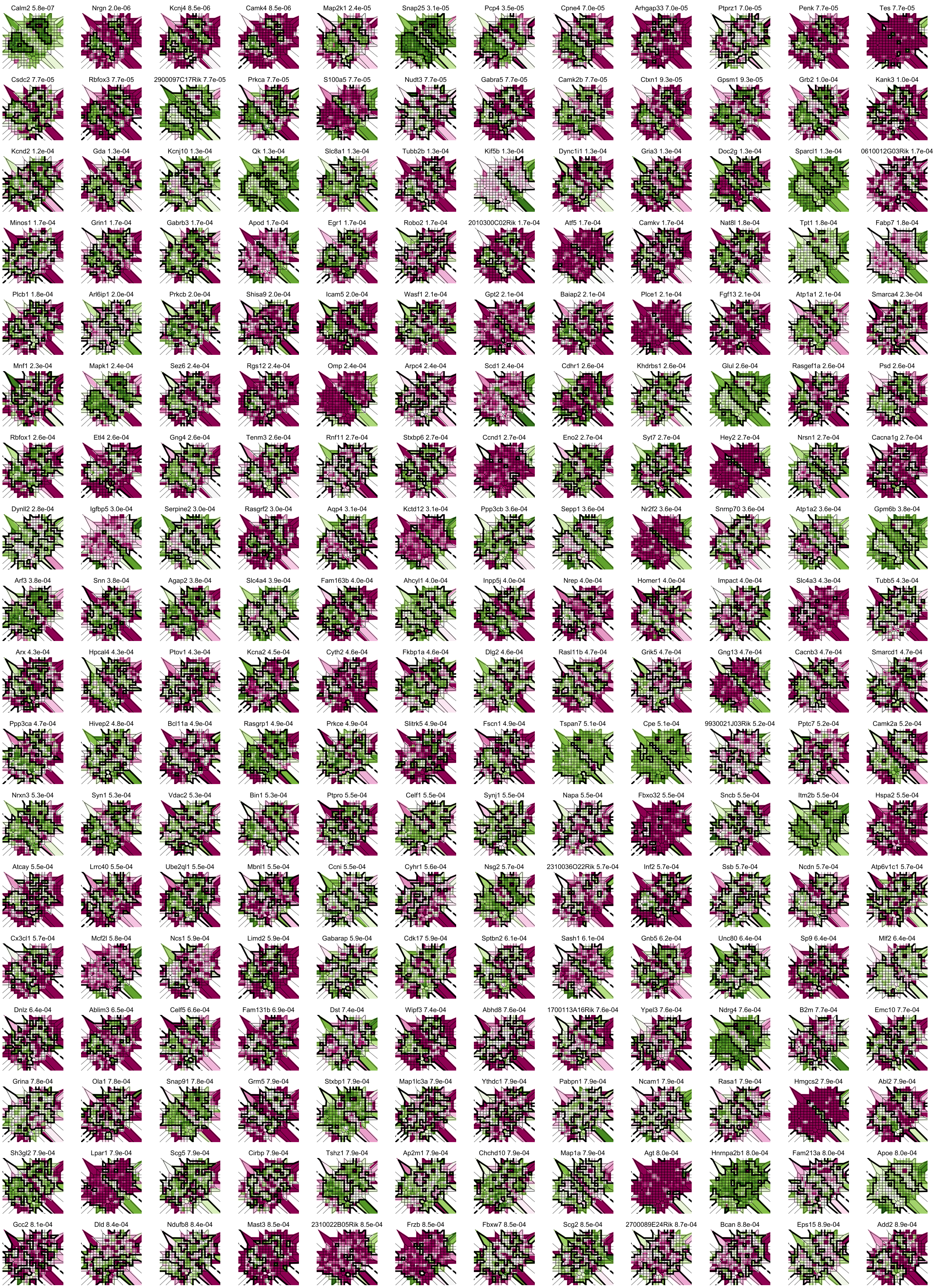

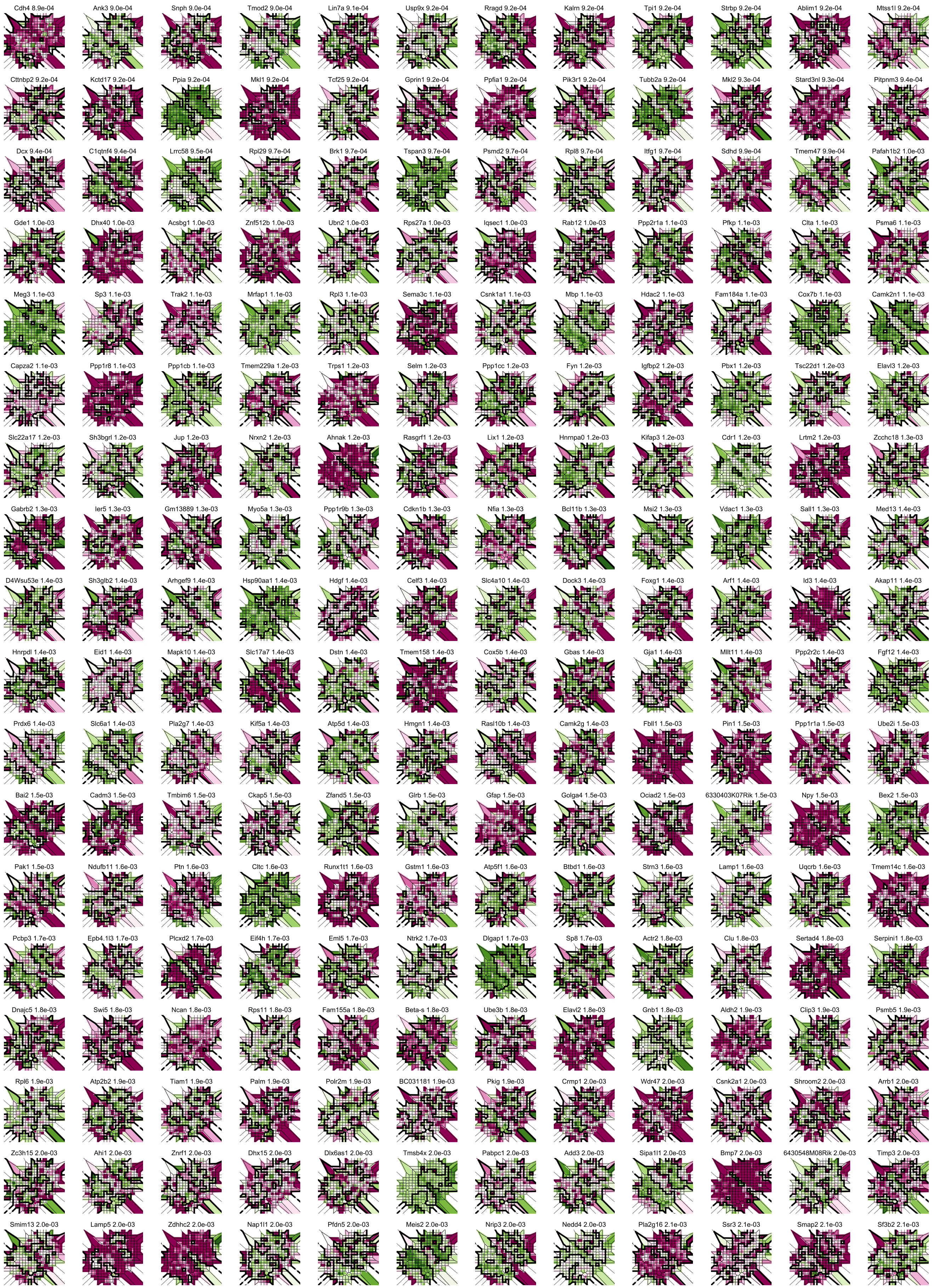

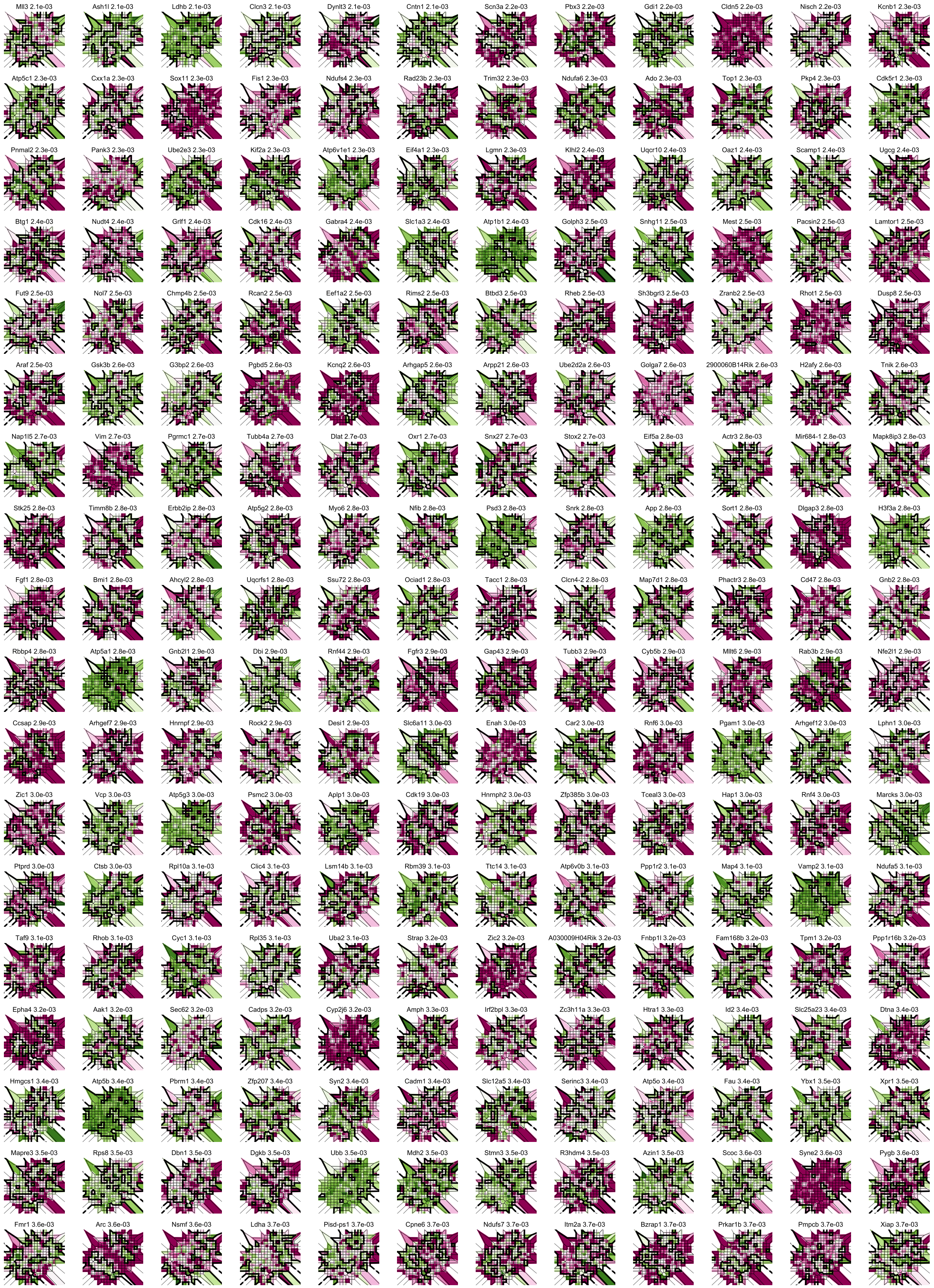

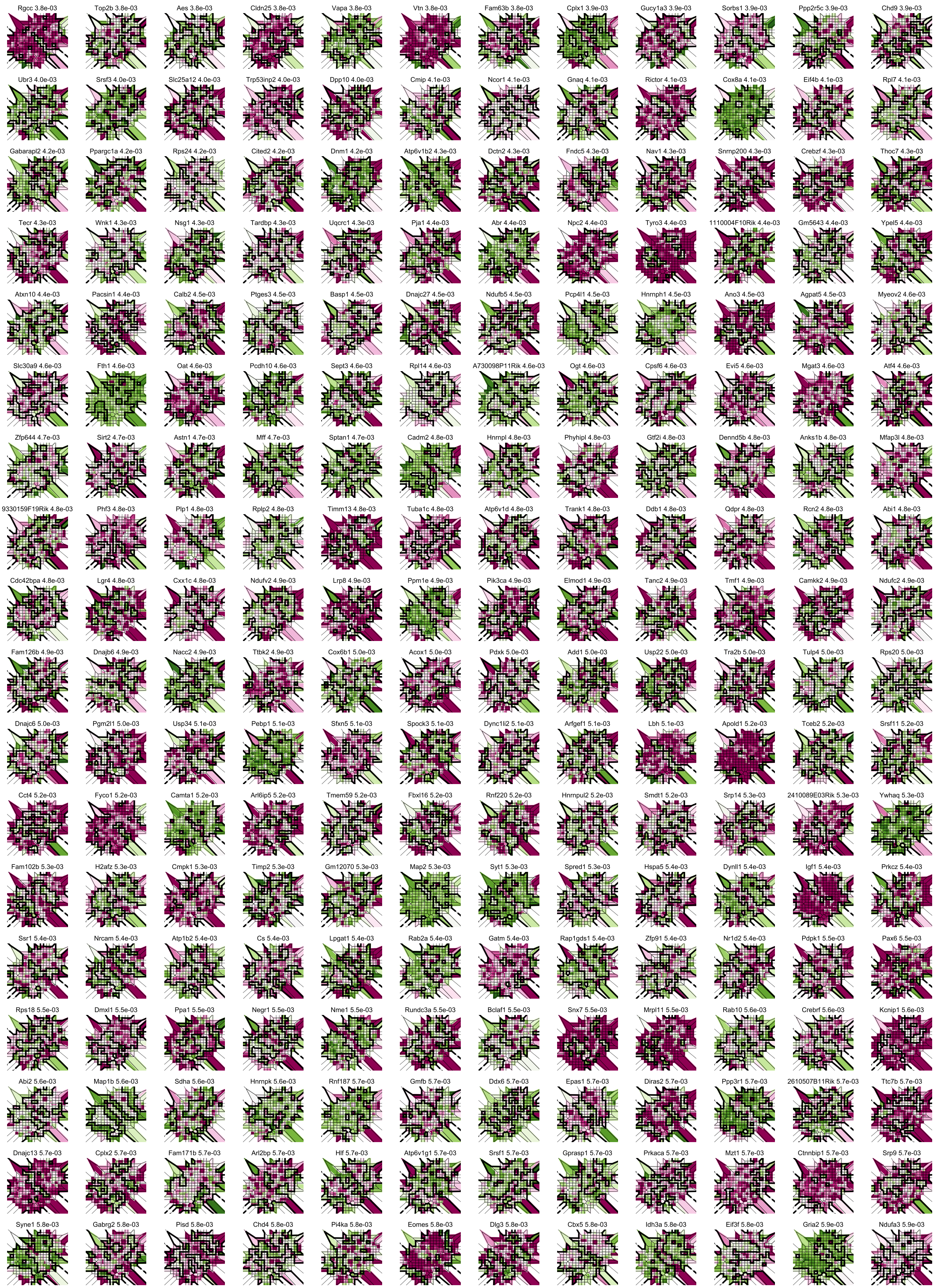

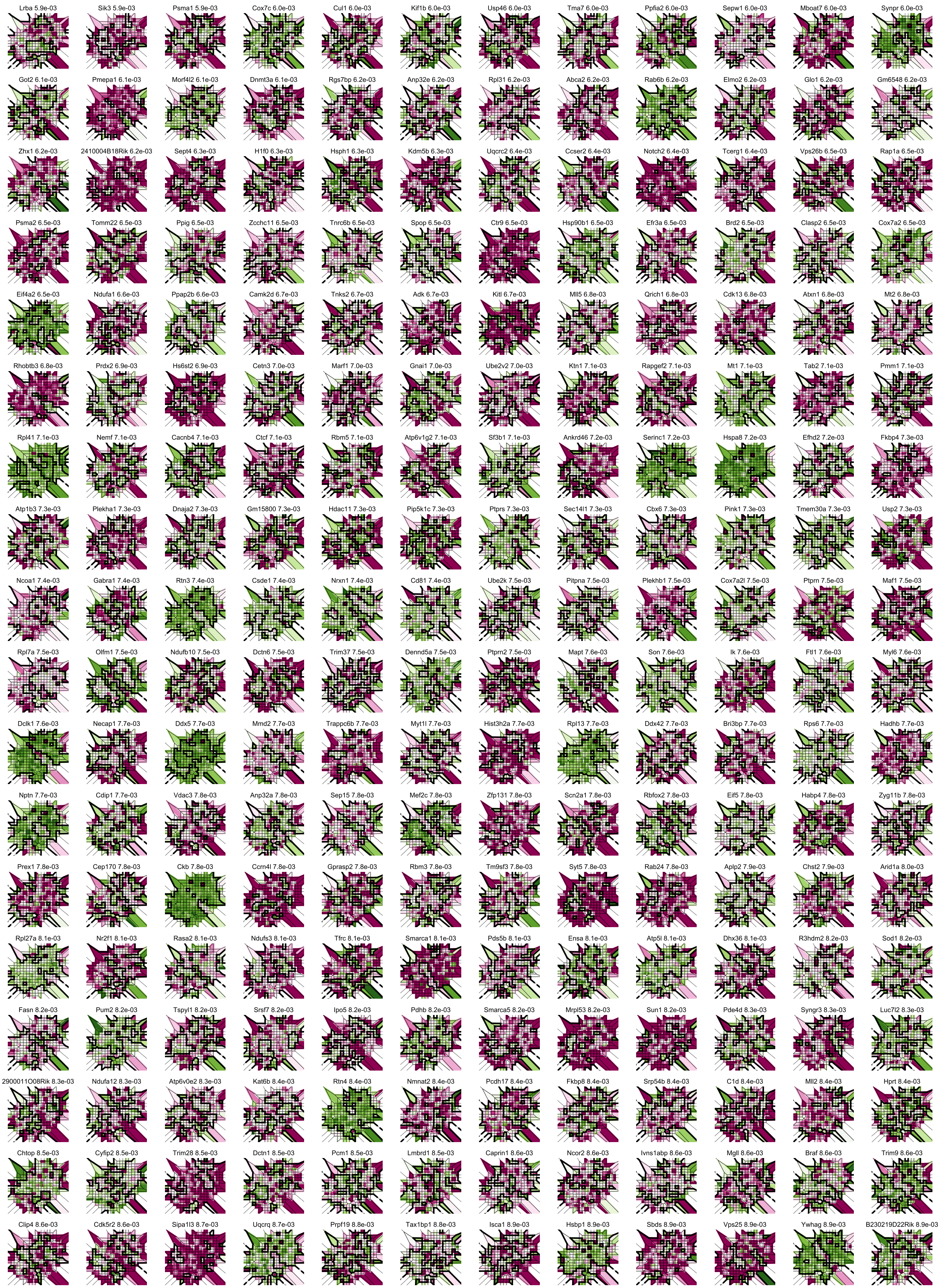

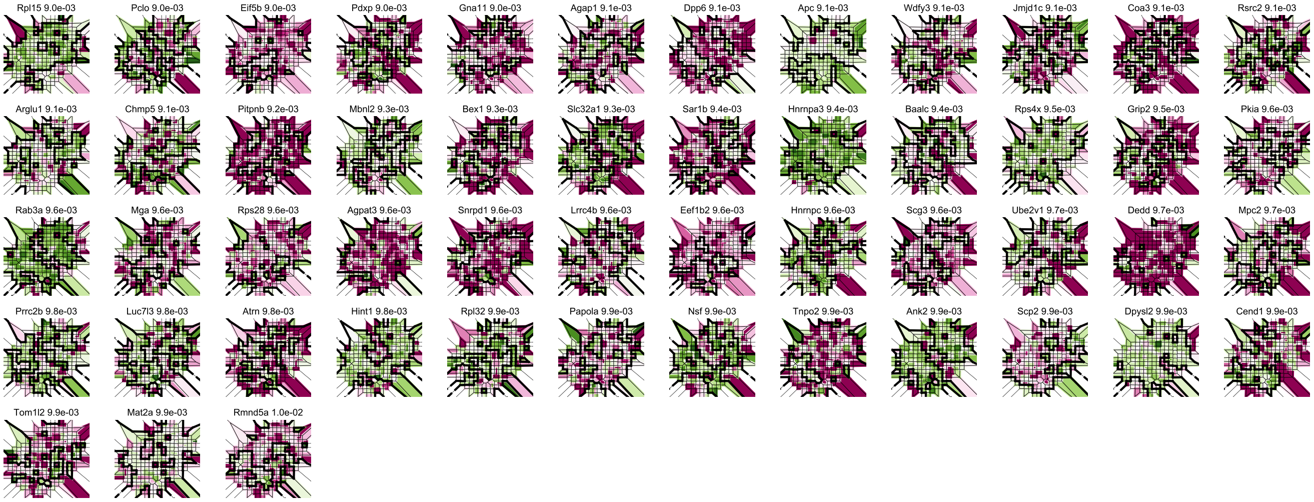
