## Supplementary material for "Identification of spatially variable genes with graph cuts"

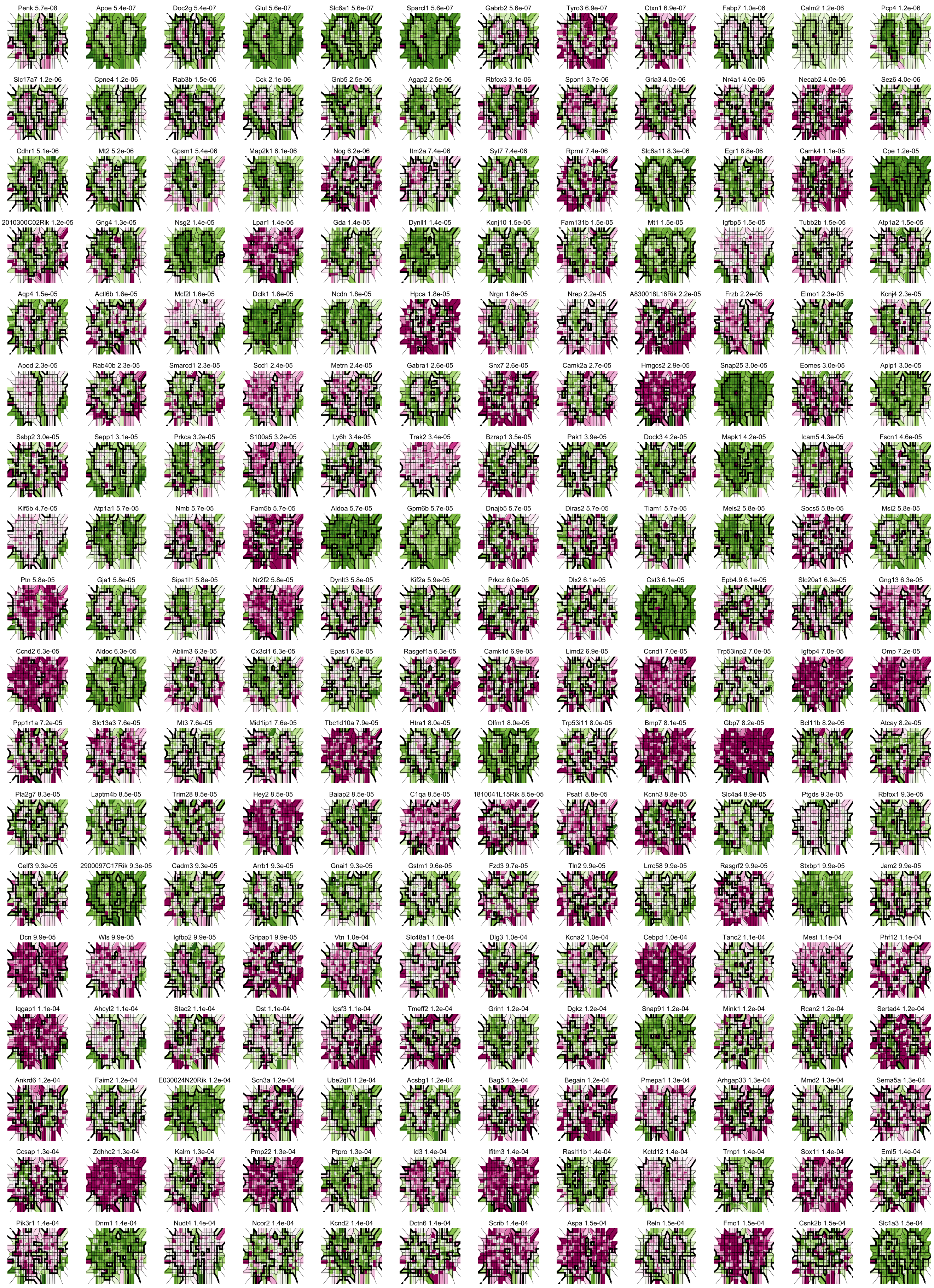
Penk 5.7e-08
Apoe 5.4e-07
Doc2g 5.4e-07
Glul 5.6e-07
Slc6a1 5.6e-07
Sparcl1 5.6e-07
Gabrb2 5.6e-07
Tyro3 6.9e-07
Ctxn1 6.9e-07
Fabp7 1.0e-06
Calm2 1.2e-06
Pcp4 1.2e-06
Slc17a7 1.2e-06
Cpne4 1.2e-06
Rab3b 1.5e-06
Cck 2.1e-06
Gnb5 2.5e-06
Agap2 2.5e-06
Rbfox3 3.1e-06
Spon1 3.7e-06
Gria3 4.0e-06
Nr4a1 4.0e-06
Necab2 4.0e-06
Sez6 4.0e-06
Cdhr1 5.1e-06
Mt2 5.2e-06
Gpsm1 5.4e-06
Map2k1 6.1e-06
Nog 6.2e-06
Itm2a 7.4e-06
Syt7 7.4e-06
Rprml 7.4e-06
Slc6a11 8.3e-06
Egr1 8.8e-06
Camk4 1.1e-05
Cpe 1.2e-05
2010300C02Rik 1.2e-05
Gng4 1.3e-05
Nsg2 1.4e-05
Lpar1 1.4e-05
Gda 1.4e-05
Dynll1 1.4e-05
Kcnj10 1.5e-05
Fam131b 1.5e-05
Mt1 1.5e-05
Igfbp5 1.5e-05
Tubb2b 1.5e-05
Atp1a2 1.5e-05
Aqp4 1.5e-05
Actl6b 1.6e-05
Mcf2l 1.6e-05
Dclk1 1.6e-05
Ncdn 1.8e-05
Hpca 1.8e-05
Nrgn 1.8e-05
Nrep 2.2e-05
A830018L16Rik 2.2e-05
Frzb 2.2e-05
Elmo1 2.3e-05
Kcnj4 2.3e-05
Apod 2.3e-05
Rab40b 2.3e-05
Smarcd1 2.3e-05
Scd1 2.4e-05
Metrn 2.4e-05
Gabra1 2.6e-05
Snx7 2.6e-05
Camk2a 2.7e-05
Hmgcs2 2.9e-05
Snap25 3.0e-05
Eomes 3.0e-05
Aplp1 3.0e-05
Ssbp2 3.0e-05
Sepp1 3.1e-05
Prkca 3.2e-05
S100a5 3.2e-05
Ly6h 3.4e-05
Trak2 3.4e-05
Bzrap1 3.5e-05
Pak1 3.9e-05
Dock3 4.2e-05
Mapk1 4.2e-05
Icam5 4.3e-05
Fscn1 4.6e-05
Kif5b 4.7e-05
Atp1a1 5.7e-05
Nmb 5.7e-05
Fam5b 5.7e-05
Aldoa 5.7e-05
Gpm6b 5.7e-05
Dnajb5 5.7e-05
Diras2 5.7e-05
Tiam1 5.7e-05
Meis2 5.8e-05
Socs5 5.8e-05
Msi2 5.8e-05
Ptn 5.8e-05
Gja1 5.8e-05
Sipa1l1 5.8e-05
Nr2f2 5.8e-05
Dynlt3 5.8e-05
Kif2a 5.9e-05
Prkcz 6.0e-05
Dlx2 6.1e-05
Cst3 6.1e-05
Epb4.9 6.1e-05
Slc20a1 6.3e-05
Gng13 6.3e-05
Ccnd2 6.3e-05
Aldoc 6.3e-05
Ablim3 6.3e-05
Cx3cl1 6.3e-05
Epas1 6.3e-05
Rasgef1a 6.3e-05
Camk1d 6.9e-05
Limd2 6.9e-05
Ccnd1 7.0e-05
Trp53inp2 7.0e-05
Igfbp4 7.0e-05
Omp 7.2e-05
Ppp1r1a 7.2e-05
Slc13a3 7.6e-05
Mt3 7.6e-05
Mid1ip1 7.6e-05
Tbc1d10a 7.9e-05
Htra1 8.0e-05
Olfm1 8.0e-05
Trp53i11 8.0e-05
Bmp7 8.1e-05
Gbp7 8.2e-05
Bcl11b 8.2e-05
Atcay 8.2e-05
Pla2g7 8.3e-05
Laptm4b 8.5e-05
Trim28 8.5e-05
Hey2 8.5e-05
Baiap2 8.5e-05
C1qa 8.5e-05
1810041L15Rik 8.5e-05
Psat1 8.8e-05
Kcnh3 8.8e-05
Slc4a4 8.9e-05
Ptgds 9.3e-05
Rbfox1 9.3e-05
Celf3 9.3e-05
2900097C17Rik 9.3e-05
Cadm3 9.3e-05
Arrb1 9.3e-05
Gnai1 9.3e-05
Gstm1 9.6e-05
Fzd3 9.7e-05
Tln2 9.9e-05
Lrrc58 9.9e-05
Rasgrf2 9.9e-05
Stxbp1 9.9e-05
Jam2 9.9e-05
Dcn 9.9e-05
Wls 9.9e-05
Igfbp2 9.9e-05
Gripap1 9.9e-05
Vtn 1.0e-04
Slc48a1 1.0e-04
Dlg3 1.0e-04
Kcna2 1.0e-04
Cebpd 1.0e-04
Tanc2 1.1e-04
Mest 1.1e-04
Phf12 1.1e-04
Iqgap1 1.1e-04
Ahcyl2 1.1e-04
Stac2 1.1e-04
Dst 1.1e-04
Igsf3 1.1e-04
Tmeff2 1.2e-04
Grin1 1.2e-04
Dgkz 1.2e-04
Snap91 1.2e-04
Mink1 1.2e-04
Rcan2 1.2e-04
Sertad4 1.2e-04
Ankrd6 1.2e-04
Faim2 1.2e-04
E030024N20Rik 1.2e-04
Scn3a 1.2e-04
Ube2ql1 1.2e-04
Acsbg1 1.2e-04
Bag5 1.2e-04
Begain 1.2e-04
Pmepa1 1.3e-04
Arhgap33 1.3e-04
Mmd2 1.3e-04
Sema5a 1.3e-04
Ccsap 1.3e-04
Zdhhc2 1.3e-04
Kalrn 1.3e-04
Pmp22 1.3e-04
Ptpro 1.3e-04
Id3 1.4e-04
Ifitm3 1.4e-04
Rasl11b 1.4e-04
Kctd12 1.4e-04
Trnp1 1.4e-04
Sox11 1.4e-04
Eml5 1.4e-04
Pik3r1 1.4e-04
Dnm1 1.4e-04
Nudt4 1.4e-04
Ncor2 1.4e-04
Kcnd2 1.4e-04
Dctn6 1.4e-04
Scrib 1.4e-04
Aspa 1.5e-04
Reln 1.5e-04
Fmo1 1.5e-04
Csnk2b 1.5e-04
Slc1a3 1.5e-04

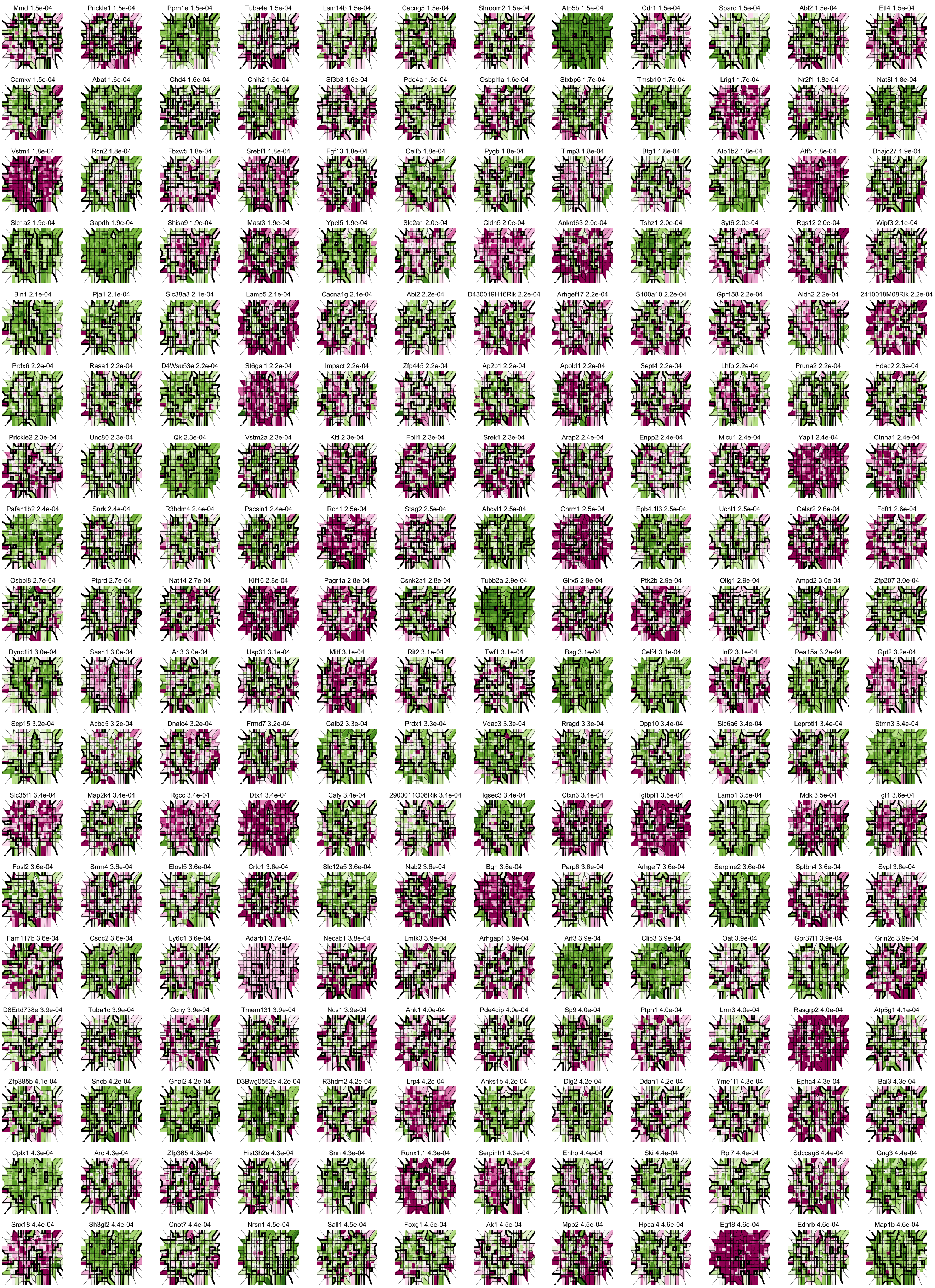
Mmd 1.5e-04
Prickle1 1.5e-04
Ppm1e 1.5e-04
Tuba4a 1.5e-04
Lsm14b 1.5e-04
Cacng5 1.5e-04
Shroom2 1.5e-04
Atp5b 1.5e-04
Cdr1 1.5e-04
Sparc 1.5e-04
Abl2 1.5e-04
Etl4 1.5e-04
Camkv 1.5e-04
Abat 1.6e-04
Chd4 1.6e-04
Cnih2 1.6e-04
Sf3b3 1.6e-04
Pde4a 1.6e-04
Osbpl1a 1.6e-04
Stxbp6 1.7e-04
Tmsb10 1.7e-04
Lrig1 1.7e-04
Nr2f1 1.8e-04
Nat8l 1.8e-04
Vstm4 1.8e-04
Rcn2 1.8e-04
Fbxw5 1.8e-04
Srebf1 1.8e-04
Fgf13 1.8e-04
Celf5 1.8e-04
Pygb 1.8e-04
Timp3 1.8e-04
Btg1 1.8e-04
Atp1b2 1.8e-04
Atf5 1.8e-04
Dnajc27 1.9e-04
Slc1a2 1.9e-04
Gapdh 1.9e-04
Shisa9 1.9e-04
Mast3 1.9e-04
Ypel5 1.9e-04
Slc2a1 2.0e-04
Cldn5 2.0e-04
Ankrd63 2.0e-04
Tshz1 2.0e-04
Syt6 2.0e-04
Rgs12 2.0e-04
Wipf3 2.1e-04
Bin1 2.1e-04
Pja1 2.1e-04
Slc38a3 2.1e-04
Lamp5 2.1e-04
Cacna1g 2.1e-04
Abi2 2.2e-04
D430019H16Rik 2.2e-04
Arhgef17 2.2e-04
S100a10 2.2e-04
Gpr158 2.2e-04
Aldh2 2.2e-04
2410018M08Rik 2.2e-04
Prdx6 2.2e-04
Rasa1 2.2e-04
D4Wsu53e 2.2e-04
St6gal1 2.2e-04
Impact 2.2e-04
Zfp445 2.2e-04
Ap2b1 2.2e-04
Apold1 2.2e-04
Sept4 2.2e-04
Lhfp 2.2e-04
Prune2 2.2e-04
Hdac2 2.3e-04
Prickle2 2.3e-04
Unc80 2.3e-04
Qk 2.3e-04
Vstm2a 2.3e-04
Kitl 2.3e-04
Fbll1 2.3e-04
Srek1 2.3e-04
Arap2 2.4e-04
Enpp2 2.4e-04
Micu1 2.4e-04
Yap1 2.4e-04
Ctnna1 2.4e-04
Pafah1b2 2.4e-04
Snrk 2.4e-04
R3hdm4 2.4e-04
Pacsin1 2.4e-04
Rcn1 2.5e-04
Stag2 2.5e-04
Ahcyl1 2.5e-04
Chrm1 2.5e-04
Epb4.1l3 2.5e-04
Uchl1 2.5e-04
Celsr2 2.6e-04
Fdft1 2.6e-04
Osbpl8 2.7e-04
Ptprd 2.7e-04
Nat14 2.7e-04
Klf16 2.8e-04
Pagr1a 2.8e-04
Csnk2a1 2.8e-04
Tubb2a 2.9e-04
Glrx5 2.9e-04
Ptk2b 2.9e-04
Olig1 2.9e-04
Ampd2 3.0e-04
Zfp207 3.0e-04
Dync1i1 3.0e-04
Sash1 3.0e-04
Arl3 3.0e-04
Usp31 3.1e-04
Mitf 3.1e-04
Rit2 3.1e-04
Twf1 3.1e-04
Bsg 3.1e-04
Celf4 3.1e-04
Inf2 3.1e-04
Pea15a 3.2e-04
Gpt2 3.2e-04
Sep15 3.2e-04
Acbd5 3.2e-04
Dnalc4 3.2e-04
Frmd7 3.2e-04
Calb2 3.3e-04
Prdx1 3.3e-04
Vdac3 3.3e-04
Rragd 3.3e-04
Dpp10 3.4e-04
Slc6a6 3.4e-04
Leprotl1 3.4e-04
Stmn3 3.4e-04
Slc35f1 3.4e-04
Map2k4 3.4e-04
Rgcc 3.4e-04
Dtx4 3.4e-04
Caly 3.4e-04
2900011O08Rik 3.4e-04
Iqsec3 3.4e-04
Ctxn3 3.4e-04
Igfbpl1 3.5e-04
Lamp1 3.5e-04
Mdk 3.5e-04
Igf1 3.6e-04
Fosl2 3.6e-04
Srrm4 3.6e-04
Elovl5 3.6e-04
Crtc1 3.6e-04
Slc12a5 3.6e-04
Nab2 3.6e-04
Bgn 3.6e-04
Parp6 3.6e-04
Arhgef7 3.6e-04
Serpine2 3.6e-04
Sptbn4 3.6e-04
Sypl 3.6e-04
Fam117b 3.6e-04
Csdc2 3.6e-04
Ly6c1 3.6e-04
Adarb1 3.7e-04
Necab1 3.8e-04
Lmtk3 3.9e-04
Arhgap1 3.9e-04
Arf3 3.9e-04
Clip3 3.9e-04
Oat 3.9e-04
Gpr37l1 3.9e-04
Grin2c 3.9e-04
D8Ertd738e 3.9e-04
Tuba1c 3.9e-04
Ccny 3.9e-04
Tmem131 3.9e-04
Ncs1 3.9e-04
Ank1 4.0e-04
Pde4dip 4.0e-04
Sp9 4.0e-04
Ptpn1 4.0e-04
Lrrn3 4.0e-04
Rasgrp2 4.0e-04
Atp5g1 4.1e-04
Zfp385b 4.1e-04
Sncb 4.2e-04
Gnai2 4.2e-04
D3Bwg0562e 4.2e-04
R3hdm2 4.2e-04
Lrp4 4.2e-04
Anks1b 4.2e-04
Dlg2 4.2e-04
Ddah1 4.2e-04
Yme1l1 4.3e-04
Epha4 4.3e-04
Bai3 4.3e-04
Cplx1 4.3e-04
Arc 4.3e-04
Zfp365 4.3e-04
Hist3h2a 4.3e-04
Snn 4.3e-04
Runx1t1 4.3e-04
Serpinh1 4.3e-04
Enho 4.4e-04
Ski 4.4e-04
Rpl7 4.4e-04
Sdccag8 4.4e-04
Gng3 4.4e-04
Snx18 4.4e-04
Sh3gl2 4.4e-04
Cnot7 4.4e-04
Nrsn1 4.5e-04
Sall1 4.5e-04
Foxg1 4.5e-04
Ak1 4.5e-04
Mpp2 4.5e-04
Hpcal4 4.6e-04
Egfl8 4.6e-04
Ednrb 4.6e-04
Map1b 4.6e-04

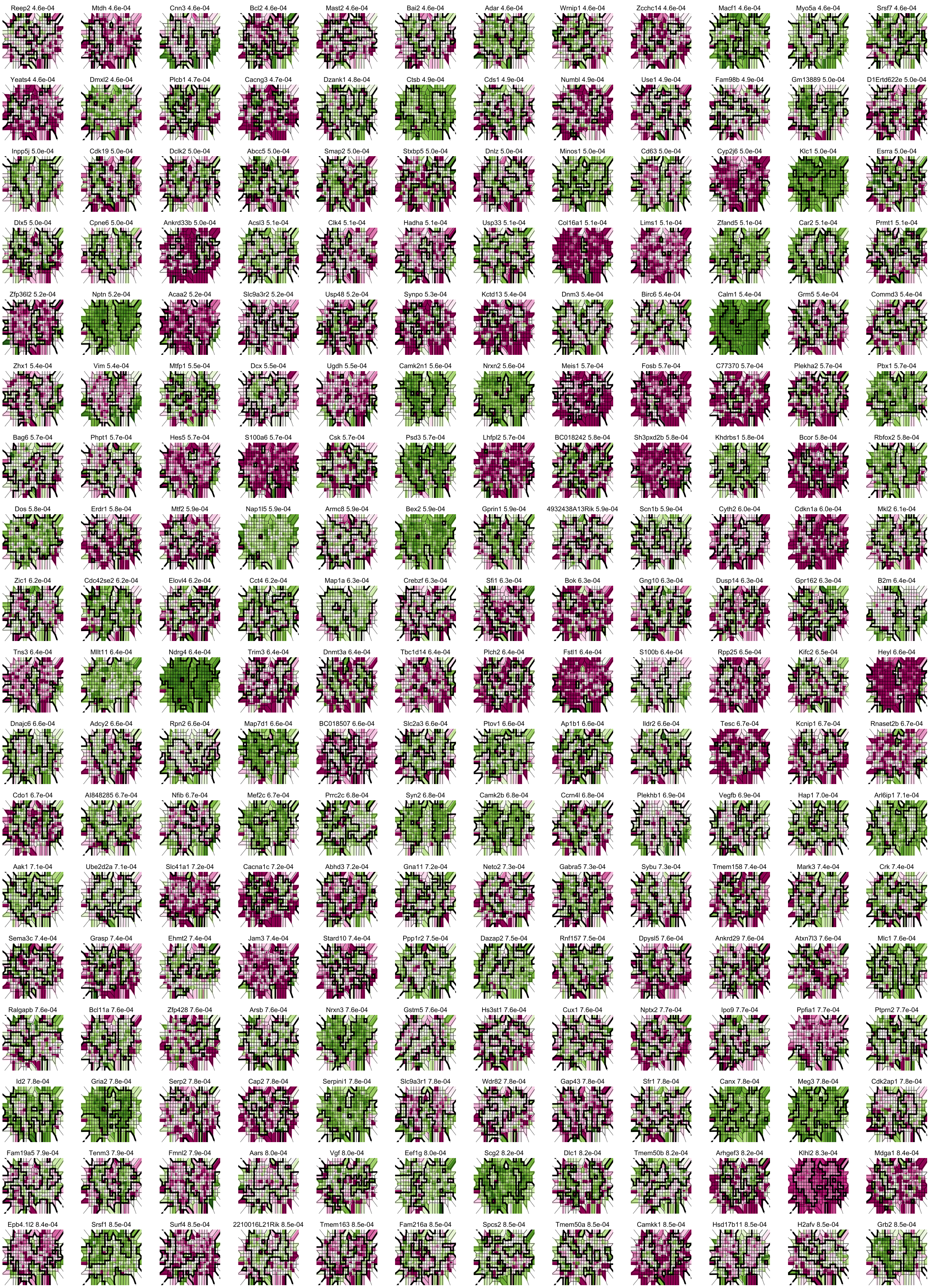

Reep2 4.6e-04
Mtdh 4.6e-04
Cnn3 4.6e-04
Bcl2 4.6e-04
Mast2 4.6e-04
Bai2 4.6e-04
Adar 4.6e-04
Wrnip1 4.6e-04
Zcchc14 4.6e-04
Macf1 4.6e-04
Myo5a 4.6e-04
Srsf7 4.6e-04
Yeats4 4.6e-04
Dmxl2 4.6e-04
Plcb1 4.7e-04
Cacng3 4.7e-04
Dzank1 4.8e-04
Ctsb 4.9e-04
Cds1 4.9e-04
Numbl 4.9e-04
Use1 4.9e-04
Fam98b 4.9e-04
Gm13889 5.0e-04
D1Ertd622e 5.0e-04
Inpp5j 5.0e-04
Cdk19 5.0e-04
Dclk2 5.0e-04
Abcc5 5.0e-04
Smap2 5.0e-04
Stxbp5 5.0e-04
Dnlz 5.0e-04
Minos1 5.0e-04
Cd63 5.0e-04
Cyp2j6 5.0e-04
Klc1 5.0e-04
Esrra 5.0e-04
Dlx5 5.0e-04
Cpne6 5.0e-04
Ankrd33b 5.0e-04
Acsl3 5.1e-04
Clk4 5.1e-04
Hadha 5.1e-04
Usp33 5.1e-04
Col16a1 5.1e-04
Lims1 5.1e-04
Zfand5 5.1e-04
Car2 5.1e-04
Prmt1 5.1e-04
Zfp36l2 5.2e-04
Nptn 5.2e-04
Acaa2 5.2e-04
Slc9a3r2 5.2e-04
Usp48 5.2e-04
Synpo 5.3e-04
Kctd13 5.4e-04
Dnm3 5.4e-04
Birc6 5.4e-04
Calm1 5.4e-04
Grm5 5.4e-04
Commd3 5.4e-04
Zhx1 5.4e-04
Vim 5.4e-04
Mtfp1 5.5e-04
Dcx 5.5e-04
Ugdh 5.5e-04
Camk2n1 5.6e-04
Nrxn2 5.6e-04
Meis1 5.7e-04
Fosb 5.7e-04
C77370 5.7e-04
Plekha2 5.7e-04
Pbx1 5.7e-04
Bag6 5.7e-04
Phpt1 5.7e-04
Hes5 5.7e-04
S100a6 5.7e-04
Csk 5.7e-04
Psd3 5.7e-04
Lhfpl2 5.7e-04
BC018242 5.8e-04
Sh3pxd2b 5.8e-04
Khdrbs1 5.8e-04
Bcor 5.8e-04
Rbfox2 5.8e-04
Dos 5.8e-04
Erdr1 5.8e-04
Mtf2 5.9e-04
Nap1l5 5.9e-04
Armc8 5.9e-04
Bex2 5.9e-04
Gprin1 5.9e-04
4932438A13Rik 5.9e-04
Scn1b 5.9e-04
Cyth2 6.0e-04
Cdkn1a 6.0e-04
Mkl2 6.1e-04
Zic1 6.2e-04
Cdc42se2 6.2e-04
Elovl4 6.2e-04
Cct4 6.2e-04
Map1a 6.3e-04
Crebzf 6.3e-04
Sfi1 6.3e-04
Bok 6.3e-04
Gng10 6.3e-04
Dusp14 6.3e-04
Gpr162 6.3e-04
B2m 6.4e-04
Tns3 6.4e-04
Mllt11 6.4e-04
Ndrg4 6.4e-04
Trim3 6.4e-04
Dnmt3a 6.4e-04
Tbc1d14 6.4e-04
Plch2 6.4e-04
Fstl1 6.4e-04
S100b 6.4e-04
Rpp25 6.5e-04
Kifc2 6.5e-04
Heyl 6.6e-04
Dnajc6 6.6e-04
Adcy2 6.6e-04
Rpn2 6.6e-04
Map7d1 6.6e-04
BC018507 6.6e-04
Slc2a3 6.6e-04
Ptov1 6.6e-04
Ap1b1 6.6e-04
Ildr2 6.6e-04
Tesc 6.7e-04
Kcnip1 6.7e-04
Rnaset2b 6.7e-04
Cdo1 6.7e-04
AI848285 6.7e-04
Nfib 6.7e-04
Mef2c 6.7e-04
Prrc2c 6.8e-04
Syn2 6.8e-04
Camk2b 6.8e-04
Ccrn4l 6.8e-04
Plekhb1 6.9e-04
Vegfb 6.9e-04
Hap1 7.0e-04
Arl6ip1 7.1e-04
Aak1 7.1e-04
Ube2d2a 7.1e-04
Slc41a1 7.2e-04
Cacna1c 7.2e-04
Abhd3 7.2e-04
Gna11 7.2e-04
Neto2 7.3e-04
Gabra5 7.3e-04
Sybu 7.3e-04
Tmem158 7.4e-04
Mark3 7.4e-04
Crk 7.4e-04
Sema3c 7.4e-04
Grasp 7.4e-04
Ehmt2 7.4e-04
Jam3 7.4e-04
Stard10 7.4e-04
Ppp1r2 7.5e-04
Dazap2 7.5e-04
Rnf157 7.5e-04
Dpysl5 7.6e-04
Ankrd29 7.6e-04
Atxn7l3 7.6e-04
Mlc1 7.6e-04
Ralgapb 7.6e-04
Bcl11a 7.6e-04
Zfp428 7.6e-04
Arsb 7.6e-04
Nrxn3 7.6e-04
Gstm5 7.6e-04
Hs3st1 7.6e-04
Cux1 7.6e-04
Nptx2 7.7e-04
Ipo9 7.7e-04
Ppfia1 7.7e-04
Ptprn2 7.7e-04
Id2 7.8e-04
Gria2 7.8e-04
Serp2 7.8e-04
Cap2 7.8e-04
Serpini1 7.8e-04
Slc9a3r1 7.8e-04
Wdr82 7.8e-04
Gap43 7.8e-04
Sfr1 7.8e-04
Canx 7.8e-04
Meg3 7.8e-04
Cdk2ap1 7.8e-04
Fam19a5 7.9e-04
Tenm3 7.9e-04
Fmnl2 7.9e-04
Aars 8.0e-04
Vgf 8.0e-04
Eef1g 8.0e-04
Scg2 8.2e-04
Dlc1 8.2e-04
Tmem50b 8.2e-04
Arhgef3 8.2e-04
Klhl2 8.3e-04
Mdga1 8.4e-04
Epb4.1l2 8.4e-04
Srsf1 8.5e-04
Surf4 8.5e-04
2210016L21Rik 8.5e-04
Tmem163 8.5e-04
Fam216a 8.5e-04
Spcs2 8.2e-04
Tmem50a 8.5e-04
Camkk1 8.5e-04
Hsd17b11 8.5e-04
H2afv 8.5e-04
Grb2 8.5e-04

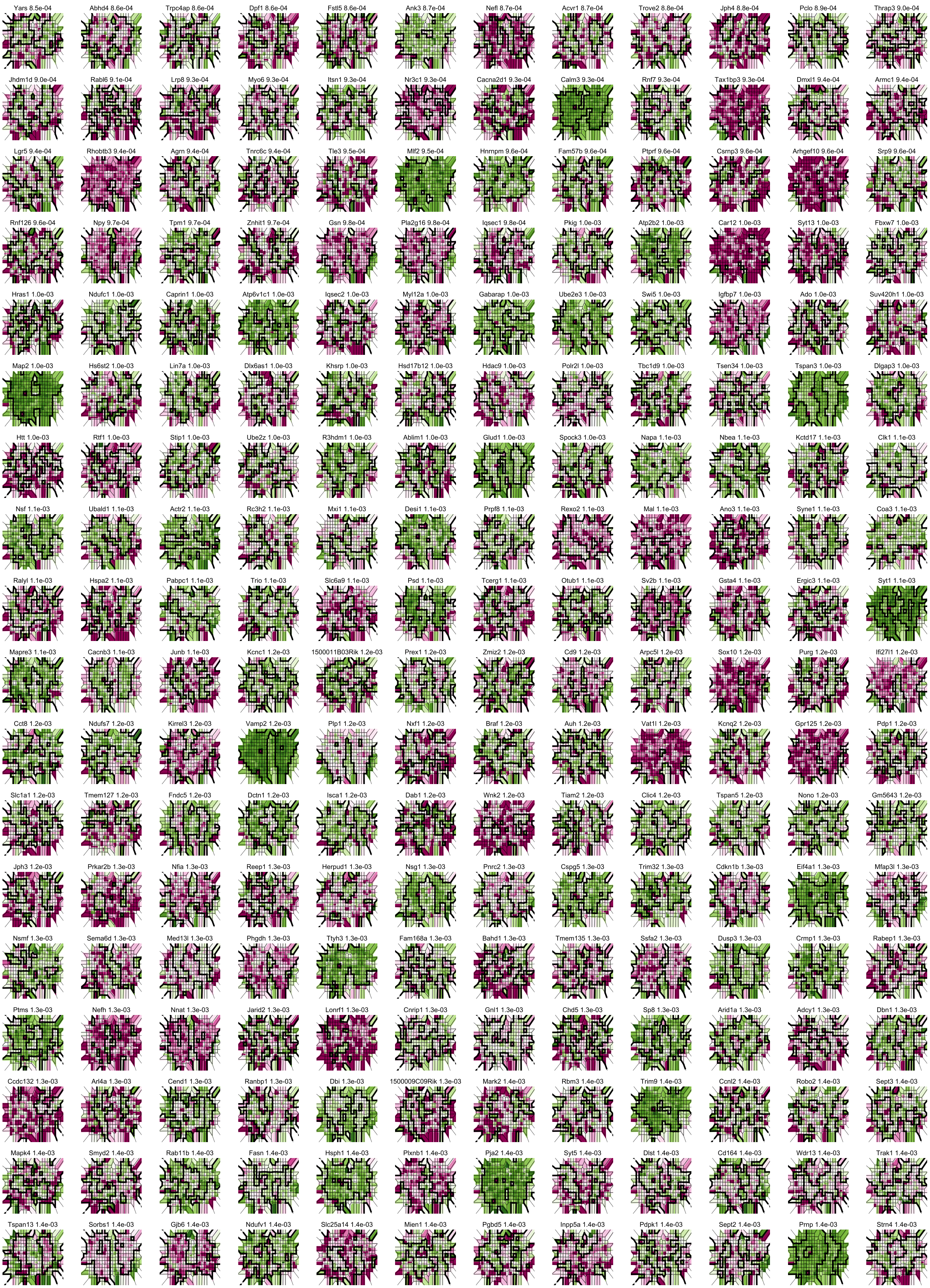
Yars 8.5e-04
Abhd4 8.6e-04
Trpc4ap 8.6e-04
Dpf1 8.6e-04
Fstl5 8.6e-04
Ank3 8.7e-04
Nefl 8.7e-04
Acvr1 8.7e-04
Trove2 8.8e-04
Jph4 8.8e-04
Pclo 8.9e-04
Thrap3 9.0e-04
Jhdm1d 9.0e-04
Rabl6 9.1e-04
Lrp8 9.3e-04
Myo6 9.3e-04
Itsn1 9.3e-04
Nr3c1 9.3e-04
Cacna2d1 9.3e-04
Calm3 9.3e-04
Rnf7 9.3e-04
Tax1bp3 9.3e-04
Dmxl1 9.4e-04
Armc1 9.4e-04
Lgr5 9.4e-04
Rhobtb3 9.4e-04
Agrn 9.4e-04
Tnrc6c 9.4e-04
Tle3 9.5e-04
Mlf2 9.5e-04
Hnrnpm 9.6e-04
Fam57b 9.6e-04
Ptprf 9.6e-04
Csrnp3 9.6e-04
Arhgef10 9.6e-04
Srp9 9.6e-04
Rnf126 9.6e-04
Npy 9.7e-04
Tpm1 9.7e-04
Znhit1 9.7e-04
Gsn 9.8e-04
Pla2g16 9.8e-04
Iqsec1 9.8e-04
Pkig 1.0e-03
Atp2b2 1.0e-03
Car12 1.0e-03
Syt13 1.0e-03
Fbxw7 1.0e-03
Hras1 1.0e-03
Ndufc1 1.0e-03
Caprin1 1.0e-03
Atp6v1c1 1.0e-03
Iqsec2 1.0e-03
Myl12a 1.0e-03
Gabarap 1.0e-03
Ube2e3 1.0e-03
Swi5 1.0e-03
Igfbp7 1.0e-03
Ado 1.0e-03
Suv420h1 1.0e-03
Map2 1.0e-03
Hs6st2 1.0e-03
Lin7a 1.0e-03
Dlx6as1 1.0e-03
Khsrp 1.0e-03
Hsd17b12 1.0e-03
Hdac9 1.0e-03
Polr2l 1.0e-03
Tbc1d9 1.0e-03
Tsen34 1.0e-03
Tspan3 1.0e-03
Dlgap3 1.0e-03
Htt 1.0e-03
Rtf1 1.0e-03
Stip1 1.0e-03
Ube2z 1.0e-03
R3hdm1 1.0e-03
Ablim1 1.0e-03
Glud1 1.0e-03
Spock3 1.0e-03
Napa 1.1e-03
Nbea 1.1e-03
Kctd17 1.1e-03
Clk1 1.1e-03
Nsf 1.1e-03
Ubald1 1.1e-03
Actr2 1.1e-03
Rc3h2 1.1e-03
Mxi1 1.1e-03
Desi1 1.1e-03
Prpf8 1.1e-03
Rexo2 1.1e-03
Mal 1.1e-03
Ano3 1.1e-03
Syne1 1.1e-03
Coa3 1.1e-03
Ralyl 1.1e-03
Hspa2 1.1e-03
Pabpc1 1.1e-03
Trio 1.1e-03
Slc6a9 1.1e-03
Psd 1.1e-03
Tcerg1 1.1e-03
Otub1 1.1e-03
Sv2b 1.1e-03
Gsta4 1.1e-03
Ergic3 1.1e-03
Syt1 1.1e-03
Mapre3 1.1e-03
Cacnb3 1.1e-03
Junb 1.1e-03
Kcnc1 1.2e-03
1500011B03Rik 1.2e-03
Prex1 1.2e-03
Zmiz2 1.2e-03
Cd9 1.2e-03
Arpc5l 1.2e-03
Sox10 1.2e-03
Purg 1.2e-03
Ifi27l1 1.2e-03
Cct8 1.2e-03
Ndufs7 1.2e-03
Kirrel3 1.2e-03
Vamp2 1.2e-03
Plp1 1.2e-03
Nxf1 1.2e-03
Braf 1.2e-03
Auh 1.2e-03
Vat1l 1.2e-03
Kcnq2 1.2e-03
Gpr125 1.2e-03
Pdp1 1.2e-03
Slc1a1 1.2e-03
Tmem127 1.2e-03
Fndc5 1.2e-03
Dctn1 1.2e-03
Isca1 1.2e-03
Dab1 1.2e-03
Wnk2 1.2e-03
Tiam2 1.2e-03
Clic4 1.2e-03
Tspan5 1.2e-03
Nono 1.2e-03
Gm5643 1.2e-03
Jph3 1.2e-03
Prkar2b 1.3e-03
Nfia 1.3e-03
Reep1 1.3e-03
Herpud1 1.3e-03
Nsg1 1.3e-03
Pnrc2 1.3e-03
Cspg5 1.3e-03
Trim32 1.3e-03
Cdkn1b 1.3e-03
Eif4a1 1.3e-03
Mfap3l 1.3e-03
Nsmf 1.3e-03
Sema6d 1.3e-03
Med13l 1.3e-03
Phgdh 1.3e-03
Ttyh3 1.3e-03
Fam168a 1.3e-03
Bahd1 1.3e-03
Tmem135 1.3e-03
Ssfa2 1.3e-03
Dusp3 1.3e-03
Crmp1 1.3e-03
Rabep1 1.3e-03
Ptms 1.3e-03
Nefh 1.3e-03
Nnat 1.3e-03
Jarid2 1.3e-03
Lonrf1 1.3e-03
Cnrip1 1.3e-03
Gnl1 1.3e-03
Chd5 1.3e-03
Sp8 1.3e-03
Arid1a 1.3e-03
Adcy1 1.3e-03
Dbn1 1.3e-03
Ccdc132 1.3e-03
Arl4a 1.3e-03
Ccnd1 1.3e-03
Ranbp1 1.3e-03
Dbi 1.3e-03
1500009C09Rik 1.3e-03
Mark2 1.4e-03
Rbm3 1.4e-03
Trim9 1.4e-03
Ccnl2 1.4e-03
Robo2 1.4e-03
Sept3 1.4e-03
Mapk4 1.4e-03
Smyd2 1.4e-03
Rab11b 1.4e-03
Fasn 1.4e-03
Hsph1 1.4e-03
Plxnb1 1.4e-03
Pja2 1.4e-03
Syt5 1.4e-03
Dlst 1.4e-03
Cd164 1.4e-03
Wdr13 1.4e-03
Trak1 1.4e-03
Tspan13 1.4e-03
Sorbs1 1.4e-03
Gjb6 1.4e-03
Ndufv1 1.4e-03
Slc25a14 1.4e-03
Mien1 1.4e-03
Pgbd5 1.4e-03
Inpp5a 1.4e-03
Pdpk1 1.4e-03
Sept2 1.4e-03
Prnp 1.4e-03
Strn4 1.4e-03

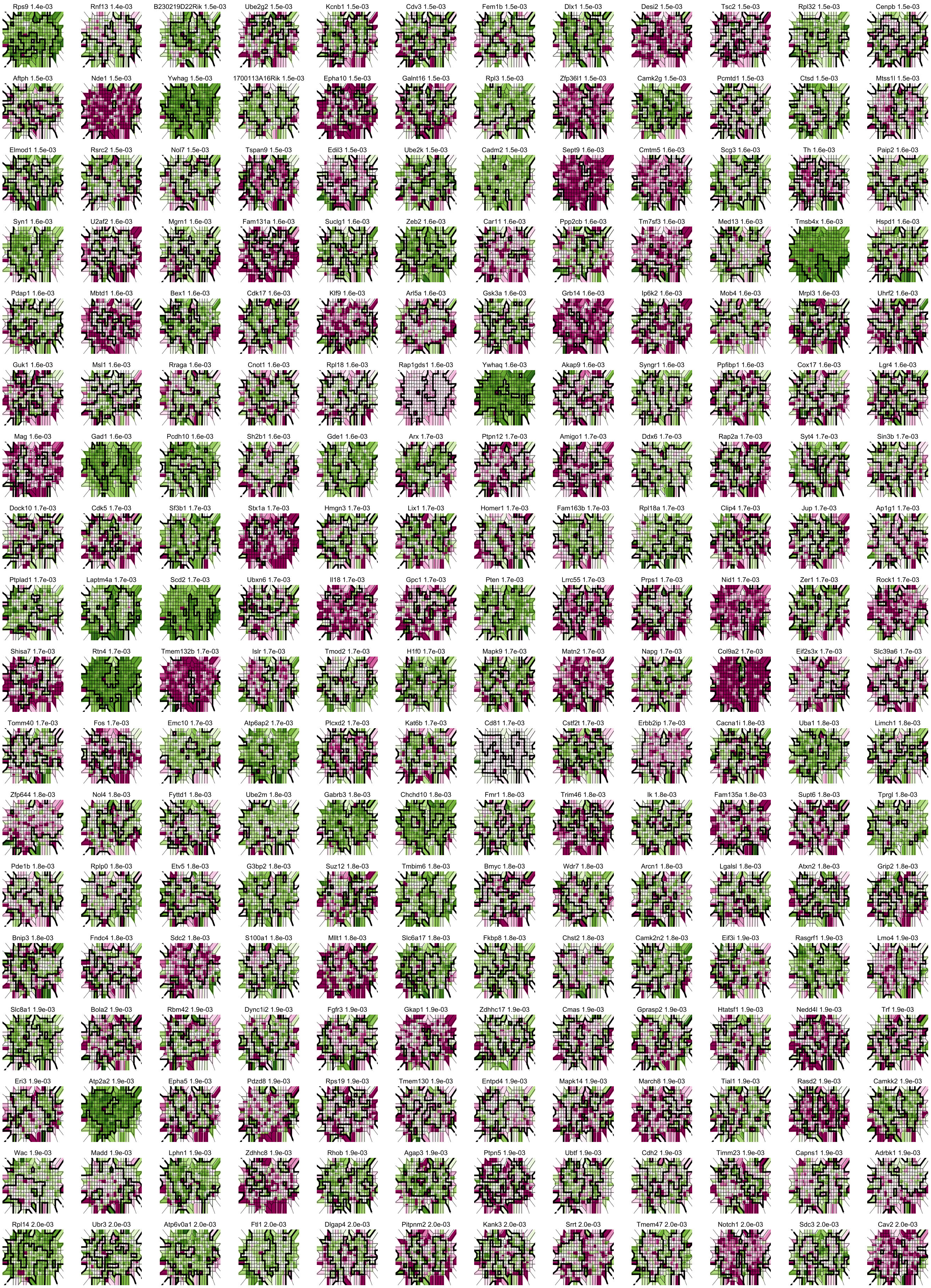
Rps9 1.4e-03
Rnf13 1.4e-03
B230219D22Rik 1.5e-03
Ube2g2 1.5e-03
Kcnb1 1.5e-03
Cdv3 1.5e-03
Fem1b 1.5e-03
Dlx1 1.5e-03
Desi2 1.5e-03
Tsc2 1.5e-03
Rpl32 1.5e-03
Cenpb 1.5e-03
Aftph 1.5e-03
Nde1 1.5e-03
Ywhag 1.5e-03
1700113A16Rik 1.5e-03
Epha10 1.5e-03
Galnt16 1.5e-03
Rpl3 1.5e-03
Zfp36l1 1.5e-03
Camk2g 1.5e-03
Pcmtd1 1.5e-03
Ctsd 1.5e-03
Mtss1l 1.5e-03
Elmod1 1.5e-03
Rsrc2 1.5e-03
Nol7 1.5e-03
Tspan9 1.5e-03
Edil3 1.5e-03
Ube2k 1.5e-03
Cadm2 1.5e-03
Sept9 1.6e-03
Cmtm5 1.6e-03
Scg3 1.6e-03
Th 1.6e-03
Paip2 1.6e-03
Syn1 1.6e-03
U2af2 1.6e-03
Mgrn1 1.6e-03
Fam131a 1.6e-03
Suclg1 1.6e-03
Zeb2 1.6e-03
Car11 1.6e-03
Ppp2cb 1.6e-03
Tm7sf3 1.6e-03
Med13 1.6e-03
Tmsb4x 1.6e-03
Hspd1 1.6e-03
Pdap1 1.6e-03
Mbtd1 1.6e-03
Bex1 1.6e-03
Cdk17 1.6e-03
Klf9 1.6e-03
Arl5a 1.6e-03
Gsk3a 1.6e-03
Grb14 1.6e-03
Ip6k2 1.6e-03
Mob4 1.6e-03
Mrpl3 1.6e-03
Uhrf2 1.6e-03
Guk1 1.6e-03
Msl1 1.6e-03
Rraga 1.6e-03
Cnot1 1.6e-03
Rpl18 1.6e-03
Rap1gds1 1.6e-03
Ywhaq 1.6e-03
Akap9 1.6e-03
Syngr1 1.6e-03
Ppfibp1 1.6e-03
Cox17 1.6e-03
Lgr4 1.6e-03
Mag 1.6e-03
Gad1 1.6e-03
Pcdh10 1.6e-03
Sh2b1 1.6e-03
Gde1 1.6e-03
Arx 1.7e-03
Ptpn12 1.7e-03
Amigo1 1.7e-03
Ddx6 1.7e-03
Rap2a 1.7e-03
Syt4 1.7e-03
Sin3b 1.7e-03
Dock10 1.7e-03
Cdk5 1.7e-03
Sf3b1 1.7e-03
Stx1a 1.7e-03
Hmgn3 1.7e-03
Lix1 1.7e-03
Homer1 1.7e-03
Fam163b 1.7e-03
Rpl18a 1.7e-03
Clip4 1.7e-03
Jup 1.7e-03
Ap1g1 1.7e-03
Ptplad1 1.7e-03
Laptm4a 1.7e-03
Scd2 1.7e-03
Ubxn6 1.7e-03
Il18 1.7e-03
Gpc1 1.7e-03
Pten 1.7e-03
Lrrc55 1.7e-03
Prps1 1.7e-03
Nid1 1.7e-03
Zer1 1.7e-03
Rock1 1.7e-03
Shisa7 1.7e-03
Rtn4 1.7e-03
Tmem132b 1.7e-03
Islr 1.7e-03
Tmod2 1.7e-03
H1f0 1.7e-03
Mapk9 1.7e-03
Matn2 1.7e-03
Napg 1.7e-03
Col9a2 1.7e-03
Eif2s3x 1.7e-03
Slc39a6 1.7e-03
Tomm40 1.7e-03
Fos 1.7e-03
Emc10 1.7e-03
Atp6ap2 1.7e-03
Plcxd2 1.7e-03
Kat6b 1.7e-03
Cd81 1.7e-03
Cstf2t 1.7e-03
Erbb2ip 1.7e-03
Cacna1i 1.8e-03
Uba1 1.8e-03
Limch1 1.8e-03
Zfp644 1.8e-03
Nol4 1.8e-03
Fyttd1 1.8e-03
Ube2m 1.8e-03
Gabrb3 1.8e-03
Chchd10 1.8e-03
Fmr1 1.8e-03
Trim46 1.8e-03
Ik 1.8e-03
Fam135a 1.8e-03
Supt6 1.8e-03
Tprgl 1.8e-03
Pde1b 1.8e-03
Rplp0 1.8e-03
Etv5 1.8e-03
G3bp2 1.8e-03
Suz12 1.8e-03
Tmbim6 1.8e-03
Bmyc 1.8e-03
Wdr7 1.8e-03
Arcn1 1.8e-03
Lgalsl 1.8e-03
Atxn2 1.8e-03
Grip2 1.8e-03
Bnip3 1.8e-03
Fndc4 1.8e-03
Sdc2 1.8e-03
S100a1 1.8e-03
Mllt1 1.8e-03
Slc6a17 1.8e-03
Fkbp8 1.8e-03
Chst2 1.8e-03
Camk2n2 1.8e-03
Eif3i 1.9e-03
Rasgrf1 1.9e-03
Lmo4 1.9e-03
Slc8a1 1.9e-03
Bola2 1.9e-03
Rbm42 1.9e-03
Dync1i2 1.9e-03
Fgfr3 1.9e-03
Gkap1 1.9e-03
Zdhhc17 1.9e-03
Cmas 1.9e-03
Gprasp2 1.9e-03
Htatsf1 1.9e-03
Nedd4l 1.9e-03
Trf 1.9e-03
Eri3 1.9e-03
Atp2a2 1.9e-03
Epha5 1.9e-03
Pdzd8 1.9e-03
Rps19 1.9e-03
Tmem130 1.9e-03
Entpd4 1.9e-03
Mapk14 1.9e-03
March8 1.9e-03
Tial1 1.9e-03
Rasd2 1.9e-03
Camkk2 1.9e-03
Wac 1.9e-03
Madd 1.9e-03
Lphn1 1.9e-03
Zdhhc8 1.9e-03
Rhob 1.9e-03
Agap3 1.9e-03
Ptpn5 1.9e-03
Ubtf 1.9e-03
Cdh2 1.9e-03
Timm23 1.9e-03
Capns1 1.9e-03
Adrbk1 1.9e-03
Rpl14 2.0e-03
Ubr3 2.0e-03
Atp6v0a1 2.0e-03
Ftl1 2.0e-03
Dlgap4 2.0e-03
Pitpnm2 2.0e-03
Kank3 2.0e-03
Srrt 2.0e-03
Tmem47 2.0e-03
Notch1 2.0e-03
Sdc3 2.0e-03
Cav2 2.0e-03

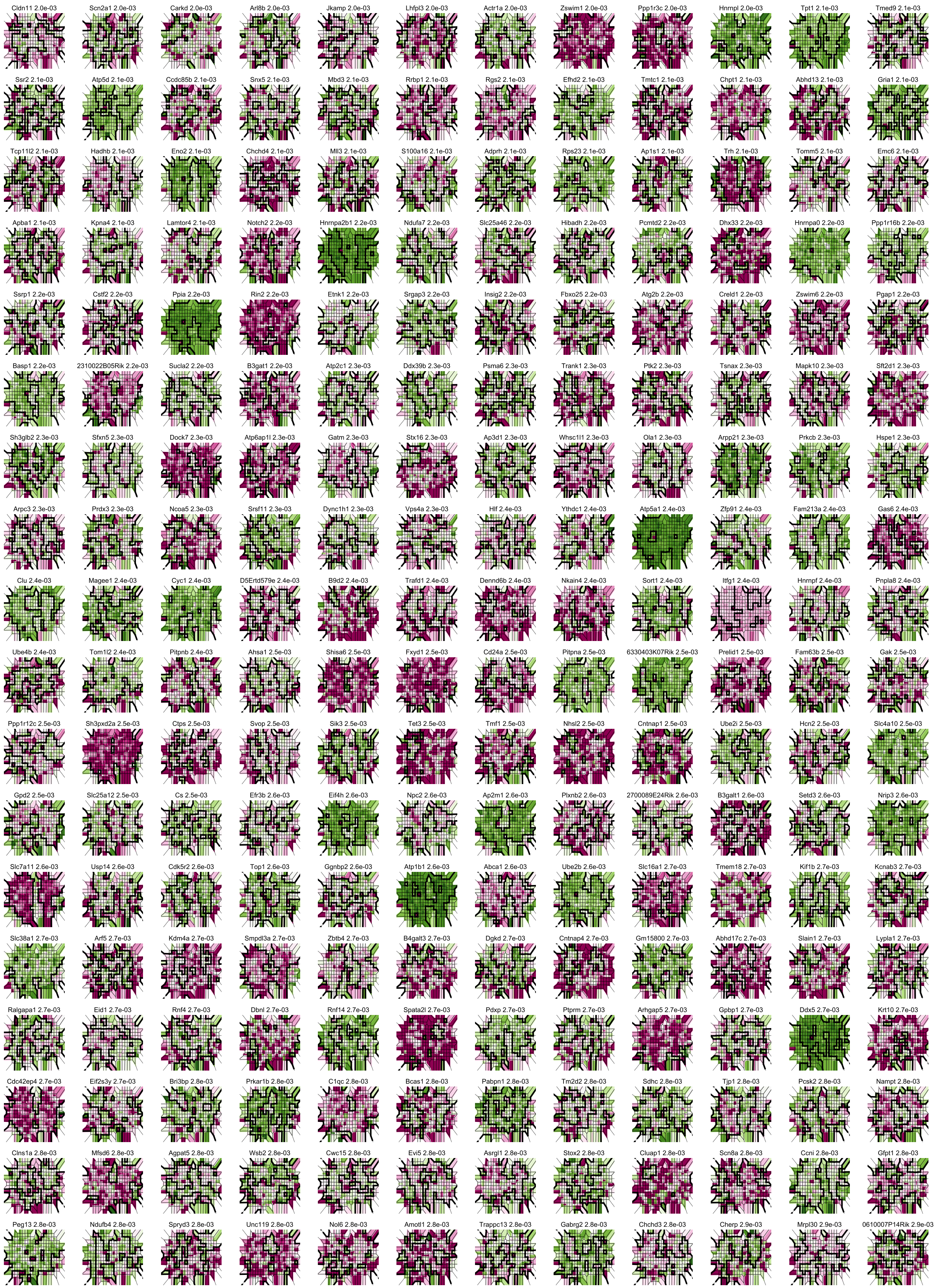
Cldn11 2.0e-03
Scn2a1 2.0e-03
Carkd 2.0e-03
Arl8b 2.0e-03
Jkamp 2.0e-03
Lhfpl3 2.0e-03
Actr1a 2.0e-03
Zswim1 2.0e-03
Ppp1r3c 2.0e-03
Hnrnpl 2.0e-03
Tpt1 2.1e-03
Tmed9 2.1e-03
Ssr2 2.1e-03
Atp5d 2.1e-03
Ccdc85b 2.1e-03
Snx5 2.1e-03
Mbd3 2.1e-03
Rrbp1 2.1e-03
Rgs2 2.1e-03
Efhd2 2.1e-03
Tmtc1 2.1e-03
Chpt1 2.1e-03
Abhd13 2.1e-03
Gria1 2.1e-03
Tcp11l2 2.1e-03
Hadhb 2.1e-03
Eno2 2.1e-03
Chchd4 2.1e-03
Mll3 2.1e-03
S100a16 2.1e-03
Adprh 2.1e-03
Rps23 2.1e-03
Ap1s1 2.1e-03
Trh 2.1e-03
Tomm5 2.1e-03
Emc6 2.1e-03
Apba1 2.1e-03
Kpna4 2.1e-03
Lamtor4 2.1e-03
Notch2 2.2e-03
Hnrnpa2b1 2.2e-03
Ndufa7 2.2e-03
Slc25a46 2.2e-03
Hibadh 2.2e-03
Pcmtd2 2.2e-03
Dhx33 2.2e-03
Hnrnpa0 2.2e-03
Ppp1r16b 2.2e-03
Ssrp1 2.2e-03
Cstf2 2.2e-03
Ppia 2.2e-03
Rin2 2.2e-03
Etnk1 2.2e-03
Srgap3 2.2e-03
Insig2 2.2e-03
Fbxo25 2.2e-03
Atg2b 2.2e-03
Creld1 2.2e-03
Zswim6 2.2e-03
Pgap1 2.2e-03
Basp1 2.2e-03
2310022B05Rik 2.2e-03
Sucla2 2.2e-03
B3gat1 2.2e-03
Atp2c1 2.3e-03
Ddx39b 2.3e-03
Psma6 2.3e-03
Trank1 2.3e-03
Ptk2 2.3e-03
Tsnax 2.3e-03
Mapk10 2.3e-03
Sft2d1 2.3e-03
Sh3glb2 2.3e-03
Sfxn5 2.3e-03
Dock7 2.3e-03
Atp6ap1l 2.3e-03
Gatm 2.3e-03
Stx16 2.3e-03
Ap3d1 2.3e-03
Whsc1l1 2.3e-03
Ola1 2.3e-03
Arpp21 2.3e-03
Prkcb 2.3e-03
Hspe1 2.3e-03
Arpc3 2.3e-03
Prdx3 2.3e-03
Ncoa5 2.3e-03
Srsf11 2.3e-03
Dync1h1 2.3e-03
Vps4a 2.3e-03
Hlf 2.4e-03
Ythdc1 2.4e-03
Atp5a1 2.4e-03
Zfp91 2.4e-03
Fam213a 2.4e-03
Gas6 2.4e-03
Clu 2.4e-03
Magee1 2.4e-03
Cyc1 2.4e-03
D5Ertd579e 2.4e-03
B9d2 2.4e-03
Trafd1 2.4e-03
Dennd6b 2.4e-03
Nkain4 2.4e-03
Sort1 2.4e-03
Itfg1 2.4e-03
Hnrnpf 2.4e-03
Pnpla8 2.4e-03
Ube4b 2.4e-03
Tom1l2 2.4e-03
Pitpnb 2.4e-03
Ahsa1 2.5e-03
Shisa6 2.5e-03
Fxyd1 2.5e-03
Cd24a 2.5e-03
Pitpna 2.5e-03
6330403K07Rik 2.5e-03
Prelid1 2.5e-03
Fam63b 2.5e-03
Gak 2.5e-03
Ppp1r12c 2.5e-03
Sh3pxd2a 2.5e-03
Ctps 2.5e-03
Svop 2.5e-03
Sik3 2.5e-03
Tet3 2.5e-03
Tmf1 2.5e-03
Nhsl2 2.5e-03
Cntnap1 2.5e-03
Ube2i 2.5e-03
Hcn2 2.5e-03
Slc4a10 2.5e-03
Gpd2 2.5e-03
Slc25a12 2.5e-03
Cs 2.5e-03
Efr3b 2.6e-03
Eif4h 2.6e-03
Npc2 2.6e-03
Ap2m1 2.6e-03
Plxnb2 2.6e-03
2700089E24Rik 2.6e-03
B3galt1 2.6e-03
Setd3 2.6e-03
Nrip3 2.6e-03
Slc7a11 2.6e-03
Usp14 2.6e-03
Cdk5r2 2.6e-03
Top1 2.6e-03
Gnpbp2 2.6e-03
Atp1b1 2.6e-03
Abca1 2.6e-03
Ube2b 2.6e-03
Slc16a1 2.7e-03
Tmem18 2.7e-03
Kif1b 2.7e-03
Kcnab3 2.7e-03
Slc38a1 2.7e-03
Arf5 2.7e-03
Kdm4a 2.7e-03
Smpdl3a 2.7e-03
Zbtb4 2.7e-03
B4galt3 2.7e-03
Dgkd 2.7e-03
Cntnap4 2.7e-03
Gm15800 2.7e-03
Abhd17c 2.7e-03
Slain1 2.7e-03
Lypla1 2.7e-03
Ralgapa1 2.7e-03
Eid1 2.7e-03
Rnf4 2.7e-03
Dbnl 2.7e-03
Rnf14 2.7e-03
Spata2l 2.7e-03
Pdxp 2.7e-03
Ptprm 2.7e-03
Arhgap5 2.7e-03
Gpbp1 2.7e-03
Ddx5 2.7e-03
Krt10 2.7e-03
Cdc42ep4 2.7e-03
Eif2s3y 2.7e-03
Bri3bp 2.8e-03
Prkar1b 2.8e-03
C1qc 2.8e-03
Bcas1 2.8e-03
Pabpn1 2.8e-03
Tm2d2 2.8e-03
Sdhc 2.8e-03
Tjp1 2.8e-03
Pcsk2 2.8e-03
Nampt 2.8e-03
Clns1a 2.8e-03
Mfsd6 2.8e-03
Agpat5 2.8e-03
Wsb2 2.8e-03
Cwc15 2.8e-03
Evi5 2.8e-03
Asrgl1 2.8e-03
Stox2 2.8e-03
Cluap1 2.8e-03
Scn8a 2.8e-03
Ccni 2.8e-03
Gfpt1 2.8e-03
Peg13 2.8e-03
Ndufb4 2.8e-03
Spryd3 2.8e-03
Unc119 2.8e-03
Nol6 2.8e-03
Amotl1 2.8e-03
Trappc13 2.8e-03
Gabrg2 2.8e-03
Chchd3 2.8e-03
Cherp 2.9e-03
Mrpl30 2.9e-03
0610007P14Rik 2.9e-03

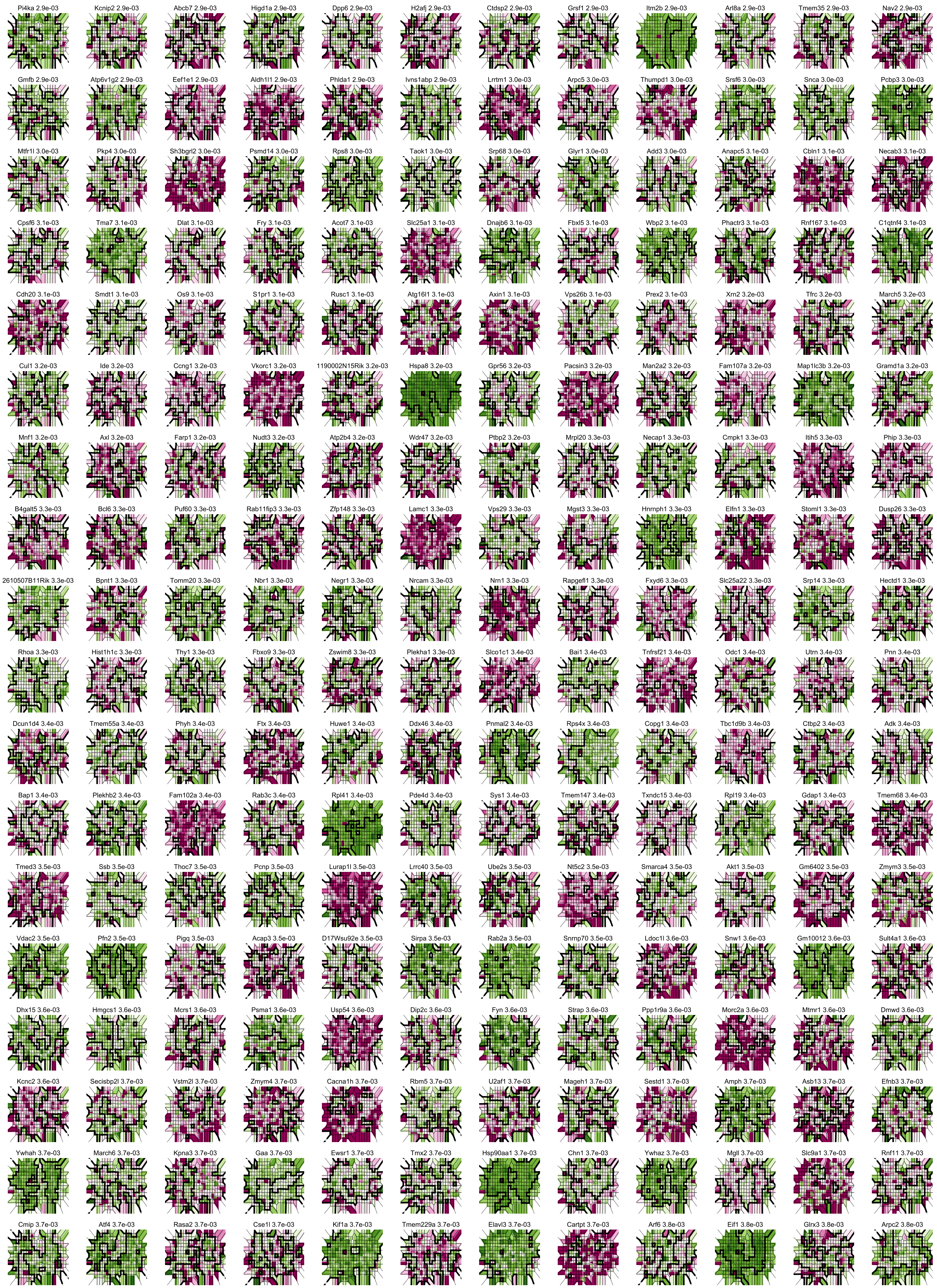
Pi4ka 2.9e-03
Kcnip2 2.9e-03
Abcb7 2.9e-03
Higd1a 2.9e-03
Dpp6 2.9e-03
H2afj 2.9e-03
Ctdsp2 2.9e-03
Grsf1 2.9e-03
Itm2b 2.9e-03
Arl8a 2.9e-03
Tmem35 2.9e-03
Nav2 2.9e-03
Gmfb 2.9e-03
Atp6v1g2 2.9e-03
Eef1e1 2.9e-03
Aldh1l1 2.9e-03
Phlda1 2.9e-03
Ivns1abp 2.9e-03
Lrrtm1 3.0e-03
Arpc5 3.0e-03
Thumpd1 3.0e-03
Srsf6 3.0e-03
Snca 3.0e-03
Pcbp3 3.0e-03
Mtfr1l 3.0e-03
Pkp4 3.0e-03
Sh3bgrl2 3.0e-03
Psmd14 3.0e-03
Rps8 3.0e-03
Taok1 3.0e-03
Srp68 3.0e-03
Glyr1 3.0e-03
Add3 3.0e-03
Anapc5 3.1e-03
Cbln1 3.1e-03
Necab3 3.1e-03
Cpsf6 3.1e-03
Tma7 3.1e-03
Dlat 3.1e-03
Fry 3.1e-03
Acot7 3.1e-03
Slc25a1 3.1e-03
Dnajb6 3.1e-03
Fbxl5 3.1e-03
Wbp2 3.1e-03
Phactr3 3.1e-03
Rnf167 3.1e-03
C1qtnf4 3.1e-03
Cdh20 3.1e-03
Smdt1 3.1e-03
Os9 3.1e-03
S1pr1 3.1e-03
Rusc1 3.1e-03
Atg16l1 3.1e-03
Axin1 3.1e-03
Vps26b 3.1e-03
Prex2 3.1e-03
Xrn2 3.2e-03
Tfrc 3.2e-03
March5 3.2e-03
Cul1 3.2e-03
Ide 3.2e-03
Ccng1 3.2e-03
Vkorc1 3.2e-03
1190002N15Rik 3.2e-03
Hspa8 3.2e-03
Gpr56 3.2e-03
Pacsin3 3.2e-03
Man2a2 3.2e-03
Fam107a 3.2e-03
Map1lc3b 3.2e-03
Gramd1a 3.2e-03
Mnf1 3.2e-03
Axl 3.2e-03
Farp1 3.2e-03
Nudt3 3.2e-03
Atp2b4 3.2e-03
Wdr47 3.2e-03
Ptbp2 3.2e-03
Mrpl20 3.3e-03
Necap1 3.3e-03
Cmpk1 3.3e-03
Itih5 3.3e-03
Phip 3.3e-03
B4galt5 3.3e-03
Bcl6 3.3e-03
Puf60 3.3e-03
Rab11fip3 3.3e-03
Zfp148 3.3e-03
Lamc1 3.3e-03
Vps29 3.3e-03
Mgst3 3.3e-03
Hnrnph1 3.3e-03
Elfn1 3.3e-03
Stoml1 3.3e-03
Dusp26 3.3e-03
2610507B11Rik 3.3e-03
Bpnt1 3.3e-03
Tomm20 3.3e-03
Nbr1 3.3e-03
Negr1 3.3e-03
Nrcam 3.3e-03
Nrn1 3.3e-03
Rapgefl1 3.3e-03
Fxyd6 3.3e-03
Slc25a22 3.3e-03
Srp14 3.3e-03
Hectd1 3.3e-03
Rhoa 3.3e-03
Hist1h1c 3.3e-03
Thy1 3.3e-03
Fbxo9 3.3e-03
Zswim8 3.3e-03
Plekha1 3.3e-03
Slco1c1 3.4e-03
Bai1 3.4e-03
Tnfrsf21 3.4e-03
Odc1 3.4e-03
Utrn 3.4e-03
Pnn 3.4e-03
Dcun1d4 3.4e-03
Tmem55a 3.4e-03
Phyh 3.4e-03
Ftx 3.4e-03
Huwe1 3.4e-03
Ddx46 3.4e-03
Pnmal2 3.4e-03
Rps4x 3.4e-03
Copg1 3.4e-03
Tbc1d9b 3.4e-03
Ctbp2 3.4e-03
Adk 3.4e-03
Bap1 3.4e-03
Plekhb2 3.4e-03
Fam102a 3.4e-03
Rab3c 3.4e-03
Rpl41 3.4e-03
Pde4d 3.4e-03
Sys1 3.4e-03
Tmem147 3.4e-03
Txndc15 3.4e-03
Rpl19 3.4e-03
Gdap1 3.4e-03
Tmem68 3.4e-03
Tmed3 3.5e-03
Ssb 3.5e-03
Thoc7 3.5e-03
Pcnp 3.5e-03
Lurap1l 3.5e-03
Lrrc40 3.5e-03
Ube2s 3.5e-03
Nt5c2 3.5e-03
Smarca4 3.5e-03
Akt1 3.5e-03
Gm6402 3.5e-03
Zmym3 3.5e-03
Vdac2 3.5e-03
Pfn2 3.5e-03
Pigq 3.5e-03
Acap3 3.5e-03
D17Wsu92e 3.5e-03
Sirpa 3.5e-03
Rab2a 3.5e-03
Snrnp70 3.5e-03
Ldoc1l 3.6e-03
Snw1 3.6e-03
Gm10012 3.6e-03
Sult4a1 3.6e-03
Dhx15 3.6e-03
Hmgcs1 3.6e-03
Mcrs1 3.6e-03
Psma1 3.6e-03
Usp54 3.6e-03
Dip2c 3.6e-03
Fyn 3.6e-03
Strap 3.6e-03
Ppp1r9a 3.6e-03
Morc2a 3.6e-03
Mtmr1 3.6e-03
Dmwd 3.6e-03
Kcnc2 3.6e-03
Secisbp2l 3.7e-03
Vstm2l 3.7e-03
Zmym4 3.7e-03
Cacna1h 3.7e-03
Rbm5 3.7e-03
U2af1 3.7e-03
Mageh1 3.7e-03
Sestd1 3.7e-03
Amph 3.7e-03
Asb13 3.7e-03
Efnb3 3.7e-03
Ywhah 3.7e-03
March6 3.7e-03
Kpna3 3.7e-03
Gaa 3.7e-03
Ewsr1 3.7e-03
Tmx2 3.7e-03
Hsp90aa1 3.7e-03
Chn1 3.7e-03
Ywhaz 3.7e-03
Mgll 3.7e-03
Slc9a1 3.7e-03
Rnf11 3.7e-03
Cmip 3.7e-03
Atf4 3.7e-03
Rasa2 3.7e-03
Cse1l 3.7e-03
Kif1a 3.7e-03
Tmem229a 3.7e-03
Elavl3 3.7e-03
Cartpt 3.7e-03
Arf6 3.8e-03
Eif1 3.8e-03
Glrx3 3.8e-03
Arpc2 3.8e-03

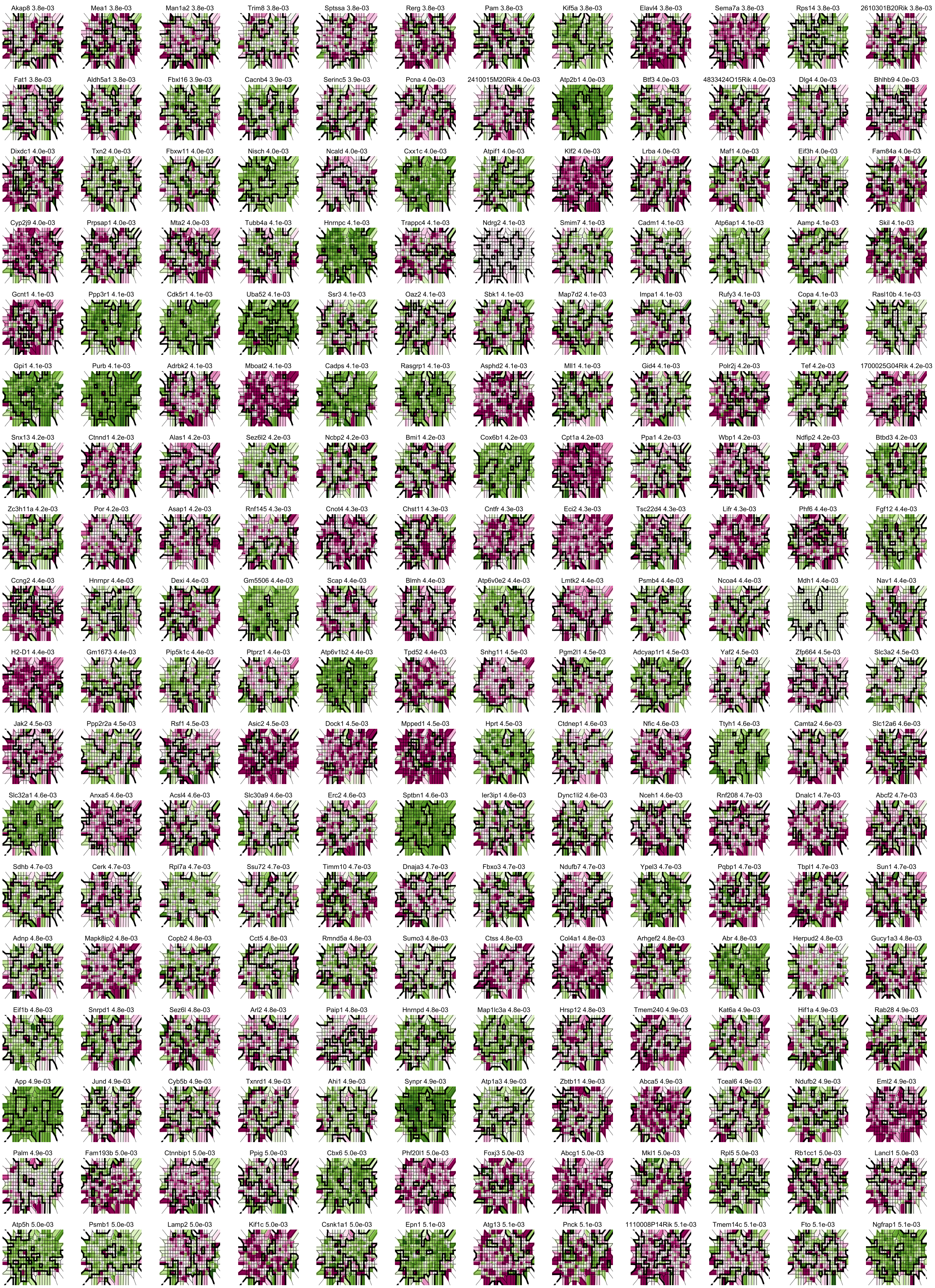

Akap8 3.8e-03
Mea1 3.8e-03
Man1a2 3.8e-03
Trim8 3.8e-03
Sptssa 3.8e-03
Rerg 3.8e-03
Pam 3.8e-03
Kif5a 3.8e-03
Elavl4 3.8e-03
Sema7a 3.8e-03
Rps14 3.8e-03
2610301B20Rik 3.8e-03
Fat1 3.8e-03
Aldh5a1 3.8e-03
Fbxl16 3.9e-03
Cacnb4 3.9e-03
Serinc5 3.9e-03
Pcna 4.0e-03
2410015M20Rik 4.0e-03
Atp2b1 4.0e-03
Btf3 4.0e-03
4833424O15Rik 4.0e-03
Dlg4 4.0e-03
Bhlhb9 4.0e-03
Dixdc1 4.0e-03
Txn2 4.0e-03
Fbxw11 4.0e-03
Nisch 4.0e-03
Ncald 4.0e-03
Cxx1c 4.0e-03
Atpif1 4.0e-03
Klf2 4.0e-03
Lrba 4.0e-03
Maf1 4.0e-03
Eif3h 4.0e-03
Fam84a 4.0e-03
Cyp2j9 4.0e-03
Prpsap1 4.0e-03
Mta2 4.0e-03
Tubb4a 4.1e-03
Hnrnpc 4.1e-03
Trappc4 4.1e-03
Ndrg2 4.1e-03
Smim7 4.1e-03
Cadm1 4.1e-03
Atp6ap1 4.1e-03
Aamp 4.1e-03
Skil 4.1e-03
Gcnt1 4.1e-03
Ppp3r1 4.1e-03
Cdk5r1 4.1e-03
Uba52 4.1e-03
Ssr3 4.1e-03
Oaz2 4.1e-03
Sbk1 4.1e-03
Map7d2 4.1e-03
Impa1 4.1e-03
Rufy3 4.1e-03
Copa 4.1e-03
Rasl10b 4.1e-03
Gpi1 4.1e-03
Purb 4.1e-03
Adrbk2 4.1e-03
Mboat2 4.1e-03
Cadps 4.1e-03
Rasgrp1 4.1e-03
Asphd2 4.1e-03
Mll1 4.1e-03
Gid4 4.1e-03
Polr2j 4.2e-03
Tef 4.2e-03
1700025G04Rik 4.2e-03
Snx13 4.2e-03
Ctnnd1 4.2e-03
Alas1 4.2e-03
Sez6l2 4.2e-03
Ncbp2 4.2e-03
Bmi1 4.2e-03
Cox6b1 4.2e-03
Cpt1a 4.2e-03
Ppa1 4.2e-03
Wbp1 4.2e-03
Ndfip2 4.2e-03
Btbd3 4.2e-03
Zc3h11a 4.2e-03
Por 4.2e-03
Asap1 4.2e-03
Rnf145 4.3e-03
Cnot4 4.3e-03
Chst11 4.3e-03
Cntfr 4.3e-03
Eci2 4.3e-03
Tsc22d4 4.3e-03
Lifr 4.3e-03
Phf6 4.4e-03
Fgf12 4.4e-03
Ccng2 4.4e-03
Hnrnpr 4.4e-03
Dexi 4.4e-03
Gm5506 4.4e-03
Scap 4.4e-03
Blmh 4.4e-03
Atp6v0e2 4.4e-03
Lmtk2 4.4e-03
Psmb4 4.4e-03
Ncoa4 4.4e-03
Mdh1 4.4e-03
Nav1 4.4e-03
H2-D1 4.4e-03
Gm1673 4.4e-03
Pip5k1c 4.4e-03
Ptprz1 4.4e-03
Atp6v1b2 4.4e-03
Tpd52 4.4e-03
Snhg11 4.5e-03
Pgm2l1 4.5e-03
Adcyap1r1 4.5e-03
Yaf2 4.5e-03
Zfp664 4.5e-03
Slc3a2 4.5e-03
Jak2 4.5e-03
Ppp2r2a 4.5e-03
Rsf1 4.5e-03
Asic2 4.5e-03
Dock1 4.5e-03
Mpped1 4.5e-03
Hprt 4.5e-03
Ctdnep1 4.6e-03
Nfic 4.6e-03
Ttyh1 4.6e-03
Camta2 4.6e-03
Slc12a6 4.6e-03
Slc32a1 4.6e-03
Anxa5 4.6e-03
Acsl4 4.6e-03
Slc30a9 4.6e-03
Erc2 4.6e-03
Sptbn1 4.6e-03
Ier3ip1 4.6e-03
Dync1li2 4.6e-03
Nceh1 4.6e-03
Rnf208 4.7e-03
Dnalc1 4.7e-03
Abcf2 4.7e-03
Sdhb 4.7e-03
Cerk 4.7e-03
Rpl7a 4.7e-03
Ssu72 4.7e-03
Timm10 4.7e-03
Dnaja3 4.7e-03
Fbxo3 4.7e-03
Ndufb7 4.7e-03
Ypel3 4.7e-03
Pqbp1 4.7e-03
Tbpl1 4.7e-03
Sun1 4.7e-03
Adnp 4.8e-03
Mapk8ip2 4.8e-03
Copb2 4.8e-03
Cct5 4.8e-03
Rmnd5a 4.8e-03
Sumo3 4.8e-03
Ctss 4.8e-03
Col4a1 4.8e-03
Arhgef2 4.8e-03
Abr 4.8e-03
Herpud2 4.8e-03
Gucy1a3 4.8e-03
Eif1b 4.8e-03
Snrpd1 4.8e-03
Sez6l 4.8e-03
Arl2 4.8e-03
Paip1 4.8e-03
Hnrnpd 4.8e-03
Map1lc3a 4.8e-03
Hrsp12 4.8e-03
Tmem240 4.9e-03
Kat6a 4.9e-03
Hif1a 4.9e-03
Rab28 4.9e-03
App 4.9e-03
Jund 4.9e-03
Cyb5b 4.9e-03
Txnrd1 4.9e-03
Ahi1 4.9e-03
Synpr 4.9e-03
Atp1a3 4.9e-03
Zbtb11 4.9e-03
Abca5 4.9e-03
Tceal6 4.9e-03
Ndufb2 4.9e-03
Eml2 4.9e-03
Palm 4.9e-03
Fam193b 5.0e-03
Ctnnbip1 5.0e-03
Ppig 5.0e-03
Cbx6 5.0e-03
Phf20l1 5.0e-03
Foxj3 5.0e-03
Abcg1 5.0e-03
Mkl1 5.0e-03
Rpl5 5.0e-03
Rb1cc1 5.0e-03
Lancl1 5.0e-03
Atp5h 5.0e-03
Psmb1 5.0e-03
Lamp2 5.0e-03
Kif1c 5.0e-03
Csnk1a1 5.0e-03
Epn1 5.1e-03
Atg13 5.1e-03
Pnck 5.1e-03
1110008P14Rik 5.1e-03
Tmem14c 5.1e-03
Fto 5.1e-03
Ngfrap1 5.1e-03

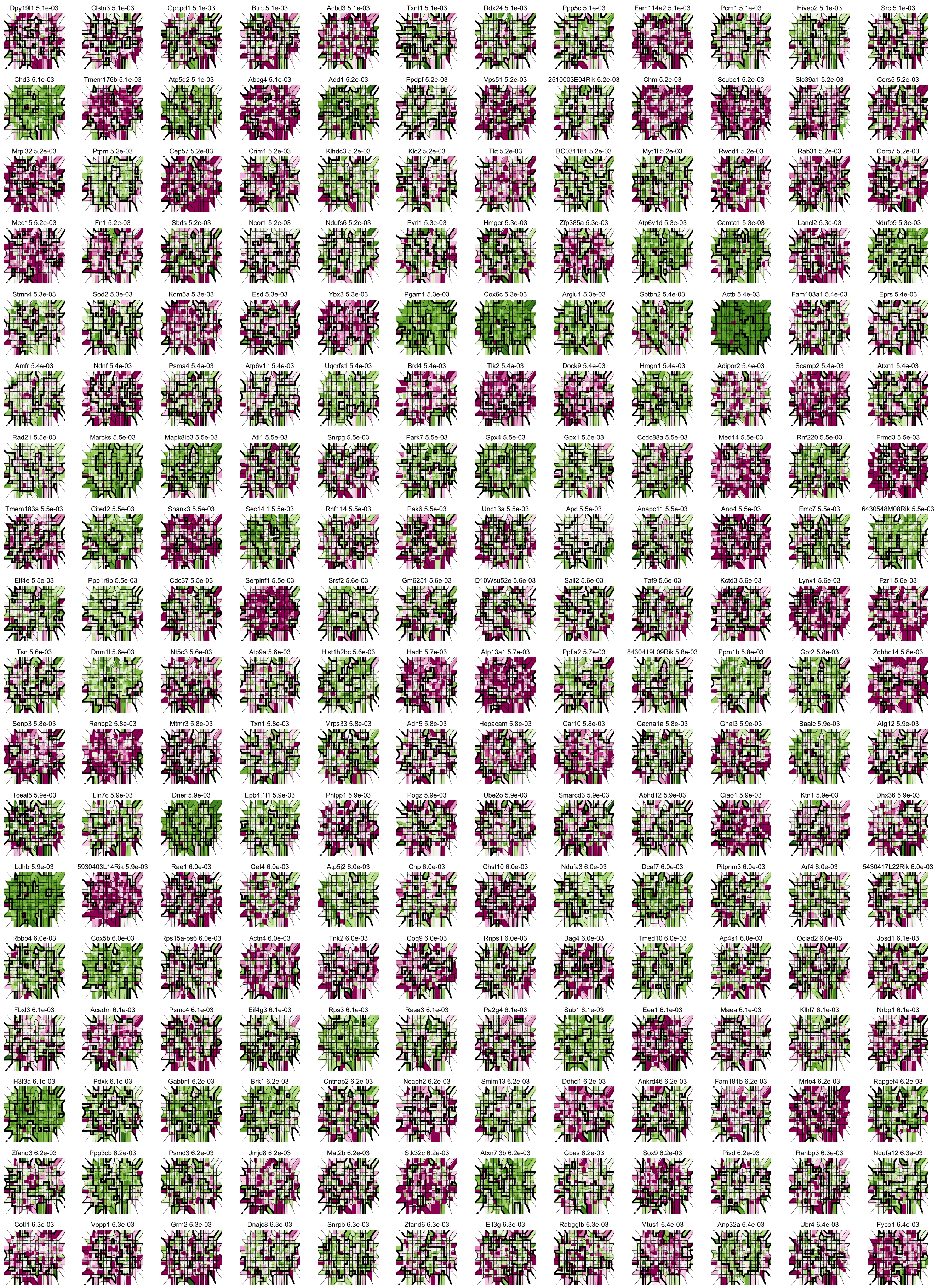

Dpy19l1 5.1e-03
Clstn3 5.1e-03
Gpcpd1 5.1e-03
Btrc 5.1e-03
Acbd3 5.1e-03
Txnl1 5.1e-03
Ddx24 5.1e-03
Ppp5c 5.1e-03
Fam114a2 5.1e-03
Pcm1 5.1e-03
Hivep2 5.1e-03
Src 5.1e-03
Chd3 5.1e-03
Tmem176b 5.1e-03
Atp5g2 5.1e-03
Abcg4 5.1e-03
Add1 5.2e-03
Ppdpf 5.2e-03
Vps51 5.2e-03
2510003E04Rik 5.2e-03
Chm 5.2e-03
Scube1 5.2e-03
Slc39a1 5.2e-03
Cers5 5.2e-03
Mrpl32 5.2e-03
Ptprn 5.2e-03
Cep57 5.2e-03
Crim1 5.2e-03
Klhdc3 5.2e-03
Klc2 5.2e-03
Tkt 5.2e-03
BC031181 5.2e-03
Myt1l 5.2e-03
Rwdd1 5.2e-03
Rab31 5.2e-03
Coro7 5.2e-03
Med15 5.2e-03
Fn1 5.2e-03
Sbds 5.2e-03
Ncor1 5.2e-03
Ndufs6 5.2e-03
Pvrl1 5.3e-03
Hmgcr 5.3e-03
Zfp385a 5.3e-03
Atp6v1d 5.3e-03
Camta1 5.3e-03
Lancl2 5.3e-03
Ndufb9 5.3e-03
Stmn4 5.3e-03
Sod2 5.3e-03
Kdm5a 5.3e-03
Esd 5.3e-03
Ybx3 5.3e-03
Pgam1 5.3e-03
Cox6c 5.3e-03
Arglu1 5.3e-03
Sptbn2 5.4e-03
Actb 5.4e-03
Fam103a1 5.4e-03
Eprs 5.4e-03
Amfr 5.4e-03
Ndnf 5.4e-03
Psma4 5.4e-03
Atp6v1h 5.4e-03
Uqcrfs1 5.4e-03
Brd4 5.4e-03
Tlk2 5.4e-03
Dock9 5.4e-03
Hmgn1 5.4e-03
Adipor2 5.4e-03
Scamp2 5.4e-03
Atxn1 5.4e-03
Rad21 5.5e-03
Marcks 5.5e-03
Mapk8ip3 5.5e-03
Atl1 5.5e-03
Snrpg 5.5e-03
Park7 5.5e-03
Gpx4 5.5e-03
Gpx1 5.5e-03
Ccdc88a 5.5e-03
Med14 5.5e-03
Rnf220 5.5e-03
Frmd3 5.5e-03
Tmem183a 5.5e-03
Cited2 5.5e-03
Shank3 5.5e-03
Sec14l1 5.5e-03
Rnf114 5.5e-03
Pak6 5.5e-03
Unc13a 5.5e-03
Apc 5.5e-03
Anapc11 5.5e-03
Ano4 5.5e-03
Emc7 5.5e-03
6430548M08Rik 5.5e-03
Eif4e 5.5e-03
Ppp1r9b 5.5e-03
Cdc37 5.5e-03
Serpinf1 5.5e-03
Srsf2 5.6e-03
Gm6251 5.6e-03
D10Wsu52e 5.6e-03
Sall2 5.6e-03
Taf9 5.6e-03
Kctd3 5.6e-03
Lynx1 5.6e-03
Fzr1 5.6e-03
Tsn 5.6e-03
Dnm1l 5.6e-03
Nt5c3 5.6e-03
Atp9a 5.6e-03
Hist1h2bc 5.6e-03
Hadh 5.7e-03
Atp13a1 5.7e-03
Ppfia2 5.7e-03
8430419L09Rik 5.8e-03
Ppm1b 5.8e-03
Got2 5.8e-03
Zdhhc14 5.8e-03
Senp3 5.8e-03
Ranbp2 5.8e-03
Mtmr3 5.8e-03
Txn1 5.8e-03
Mrps33 5.8e-03
Adh5 5.8e-03
Hepacam 5.8e-03
Car10 5.8e-03
Cacna1a 5.8e-03
Gnai3 5.9e-03
Baalc 5.9e-03
Atg12 5.9e-03
Tceal5 5.9e-03
Lin7c 5.9e-03
Dner 5.9e-03
Epb4.1l1 5.9e-03
Phlpp1 5.9e-03
Pogz 5.9e-03
Ube2o 5.9e-03
Smarcd3 5.9e-03
Abhd12 5.9e-03
Ciao1 5.9e-03
Ktn1 5.9e-03
Dhx36 5.9e-03
Ldhb 5.9e-03
5930403L14Rik 5.9e-03
Rae1 6.0e-03
Get4 6.0e-03
Atp5j2 6.0e-03
Cnp 6.0e-03
Chst10 6.0e-03
Ndufa3 6.0e-03
Dcaf7 6.0e-03
Pitpnm3 6.0e-03
Arf4 6.0e-03
5430417L22Rik 6.0e-03
Rbbp4 6.0e-03
Cox5b 6.0e-03
Rps15a-ps6 6.0e-03
Actn4 6.0e-03
Tnk2 6.0e-03
Coq9 6.0e-03
Rnps1 6.0e-03
Bag4 6.0e-03
Tmed10 6.0e-03
Ap4s1 6.0e-03
Ociad2 6.0e-03
Josd1 6.1e-03
Fbxl3 6.1e-03
Acadm 6.1e-03
Psmc4 6.1e-03
Eif4g3 6.1e-03
Rps3 6.1e-03
Rasa3 6.1e-03
Pa2g4 6.1e-03
Sub1 6.1e-03
Eea1 6.1e-03
Maea 6.1e-03
Klhl7 6.1e-03
Nrbp1 6.1e-03
H3f3a 6.1e-03
Pdxk 6.1e-03
Gabbr1 6.2e-03
Brk1 6.2e-03
Cntnap2 6.2e-03
Ncaph2 6.2e-03
Smim13 6.2e-03
Ddhd1 6.2e-03
Ankrd46 6.2e-03
Fam181b 6.2e-03
Mrto4 6.2e-03
Rapgef4 6.2e-03
Zfand3 6.2e-03
Ppp3cb 6.2e-03
Psmd3 6.2e-03
Jmjd8 6.2e-03
Mat2b 6.2e-03
Stk32c 6.2e-03
Atxn7l3b 6.2e-03
Gbas 6.2e-03
Sox9 6.2e-03
Pisd 6.2e-03
Ranbp3 6.3e-03
Ndufa12 6.3e-03
Cotl1 6.3e-03
Vopp1 6.3e-03
Grm2 6.3e-03
Dnajc8 6.3e-03
Snrpb 6.3e-03
Zfand6 6.3e-03
Eif3g 6.3e-03
Rabggtb 6.3e-03
Mtus1 6.4e-03
Anp32a 6.4e-03
Ubr4 6.4e-03
Fyco1 6.4e-03

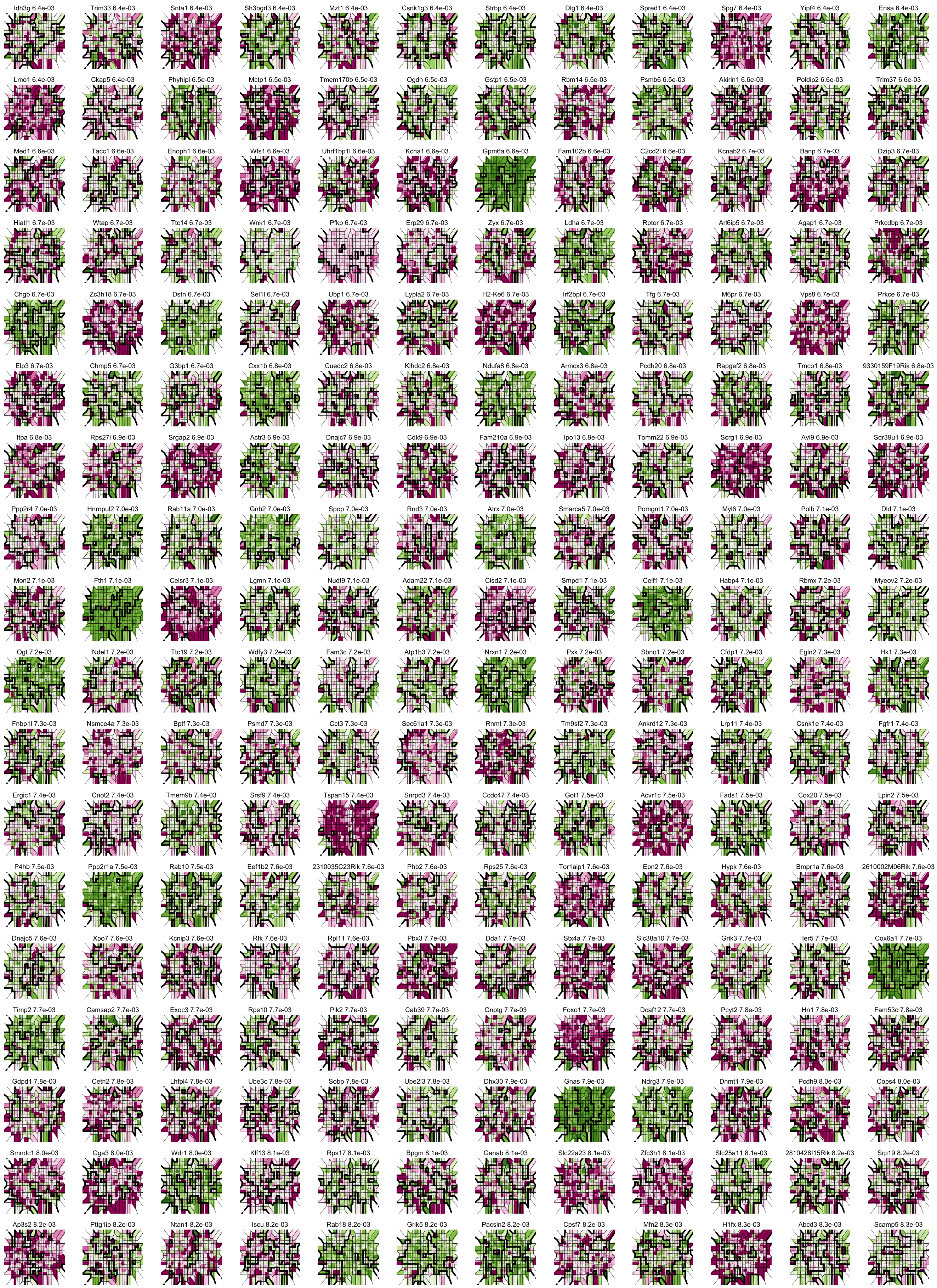

Idh3g 6.4e-03
Trim33 6.4e-03
Snta1 6.4e-03
Sh3bgrl3 6.4e-03
Mzt1 6.4e-03
Csnk1g3 6.4e-03
Strbp 6.4e-03
Dlg1 6.4e-03
Spred1 6.4e-03
Spg7 6.4e-03
Yipf4 6.4e-03
Ensa 6.4e-03
Lmo1 6.4e-03
Ckap5 6.4e-03
Phyhipl 6.5e-03
Mctp1 6.5e-03
Tmem170b 6.5e-03
Ogdh 6.5e-03
Gstp1 6.5e-03
Rbm14 6.5e-03
Psmb6 6.5e-03
Akirin1 6.6e-03
Poldip2 6.6e-03
Trim37 6.6e-03
Med1 6.6e-03
Tacc1 6.6e-03
Enoph1 6.6e-03
Wfs1 6.6e-03
Uhrf1bp1l 6.6e-03
Kcna1 6.6e-03
Gpm6a 6.6e-03
Fam102b 6.6e-03
C2cd2l 6.6e-03
Kcnab2 6.7e-03
Banp 6.7e-03
Dzip3 6.7e-03
Hiatl1 6.7e-03
Wtap 6.7e-03
Ttc14 6.7e-03
Wnk1 6.7e-03
Pfkp 6.7e-03
Erp29 6.7e-03
Zyx 6.7e-03
Ldha 6.7e-03
Rptor 6.7e-03
Arl6ip5 6.7e-03
Agap1 6.7e-03
Prkcdbp 6.7e-03
Chgb 6.7e-03
Zc3h18 6.7e-03
Dstn 6.7e-03
Sel1l 6.7e-03
Ubp1 6.7e-03
Lypla2 6.7e-03
H2-Ke6 6.7e-03
Irf2bpl 6.7e-03
Tfg 6.7e-03
M6pr 6.7e-03
Vps8 6.7e-03
Prkce 6.7e-03
Elp3 6.7e-03
Chmp5 6.7e-03
G3bp1 6.7e-03
Cxx1b 6.8e-03
Cuedc2 6.8e-03
Klhdc2 6.8e-03
Ndufa8 6.8e-03
Armcx3 6.8e-03
Pcdh20 6.8e-03
Rapgef2 6.8e-03
Tmco1 6.8e-03
9330159F19Rik 6.8e-03
Itpa 6.8e-03
Rps27l 6.9e-03
Srgap2 6.9e-03
Actr3 6.9e-03
Dnajc7 6.9e-03
Cdk9 6.9e-03
Fam210a 6.9e-03
Ipo13 6.9e-03
Tomm22 6.9e-03
Scrg1 6.9e-03
Avl9 6.9e-03
Sdr39u1 6.9e-03
Ppp2r4 7.0e-03
Hnrnpul2 7.0e-03
Rab11a 7.0e-03
Gnb2 7.0e-03
Spop 7.0e-03
Rnd3 7.0e-03
Atrx 7.0e-03
Smarca5 7.0e-03
Pomgnt1 7.0e-03
Myl6 7.0e-03
Polb 7.1e-03
Dld 7.1e-03
Mon2 7.1e-03
Fth1 7.1e-03
Celsr3 7.1e-03
Lgmn 7.1e-03
Nudt9 7.1e-03
Adam22 7.1e-03
Cisd2 7.1e-03
Smpd1 7.1e-03
Celf1 7.1e-03
Habp4 7.1e-03
Rbmx 7.2e-03
Myeov2 7.2e-03
Ogt 7.2e-03
Ndel1 7.2e-03
Ttc19 7.2e-03
Wdfy3 7.2e-03
Fam3c 7.2e-03
Atp1b3 7.2e-03
Nrxn1 7.2e-03
Pxk 7.2e-03
Sbno1 7.2e-03
Cfdp1 7.2e-03
Egln2 7.3e-03
Hk1 7.3e-03
Fnbp1l 7.3e-03
Nsmce4a 7.3e-03
Bptf 7.3e-03
Psmd7 7.3e-03
Cct3 7.3e-03
Sec61a1 7.3e-03
Rnmt 7.3e-03
Tm9sf2 7.3e-03
Ankrd12 7.3e-03
Lrp11 7.4e-03
Csnk1e 7.4e-03
Fgfr1 7.4e-03
Ergic1 7.4e-03
Cnot2 7.4e-03
Tmem9b 7.4e-03
Srsf9 7.4e-03
Tspan15 7.4e-03
Snrpd3 7.4e-03
Ccdc47 7.4e-03
Got1 7.5e-03
Acvr1c 7.5e-03
Fads1 7.5e-03
Cox20 7.5e-03
Lpin2 7.5e-03
P4hb 7.5e-03
Ppp2r1a 7.5e-03
Rab10 7.5e-03
Eef1b2 7.6e-03
2310035C23Rik 7.6e-03
Phb2 7.6e-03
Rps25 7.6e-03
Tor1aip1 7.6e-03
Epn2 7.6e-03
Hypk 7.6e-03
Bmpr1a 7.6e-03
2610002M06Rik 7.6e-03
Dnajc5 7.6e-03
Xpo7 7.6e-03
Kcnip3 7.6e-03
Rfk 7.6e-03
Rpl11 7.6e-03
Pbx3 7.7e-03
Dda1 7.7e-03
Stx4a 7.7e-03
Slc38a10 7.7e-03
Grik3 7.7e-03
Ier5 7.7e-03
Cox6a1 7.7e-03
Timp2 7.7e-03
Camsap2 7.7e-03
Exoc3 7.7e-03
Rps10 7.7e-03
Plk2 7.7e-03
Cab39 7.7e-03
Gnptg 7.7e-03
Foxo1 7.7e-03
Dcaf12 7.7e-03
Pcyt2 7.8e-03
Hn1 7.8e-03
Fam53c 7.8e-03
Gdpd1 7.8e-03
Cetn2 7.8e-03
Lhfpl4 7.8e-03
Ube3c 7.8e-03
Sobp 7.8e-03
Ube2l3 7.8e-03
Dhx30 7.9e-03
Gnas 7.9e-03
Ndrg3 7.9e-03
Dnmt1 7.9e-03
Pcdh9 8.0e-03
Cops4 8.0e-03
Smndc1 8.0e-03
Gga3 8.0e-03
Wdr1 8.0e-03
Klf13 8.1e-03
Rps17 8.1e-03
Bpgm 8.1e-03
Ganab 8.1e-03
Slc22a23 8.1e-03
Zfc3h1 8.1e-03
Slc25a11 8.1e-03
2810428I15Rik 8.2e-03
Srp19 8.2e-03
Ap3s2 8.2e-03
Pttg1ip 8.2e-03
Ntan1 8.2e-03
Iscu 8.2e-03
Rab18 8.2e-03
Grik5 8.2e-03
Pacsin2 8.2e-03
Cpsf7 8.2e-03
Mfn2 8.3e-03
H1fx 8.3e-03
Abcd3 8.3e-03
Scamp5 8.3e-03

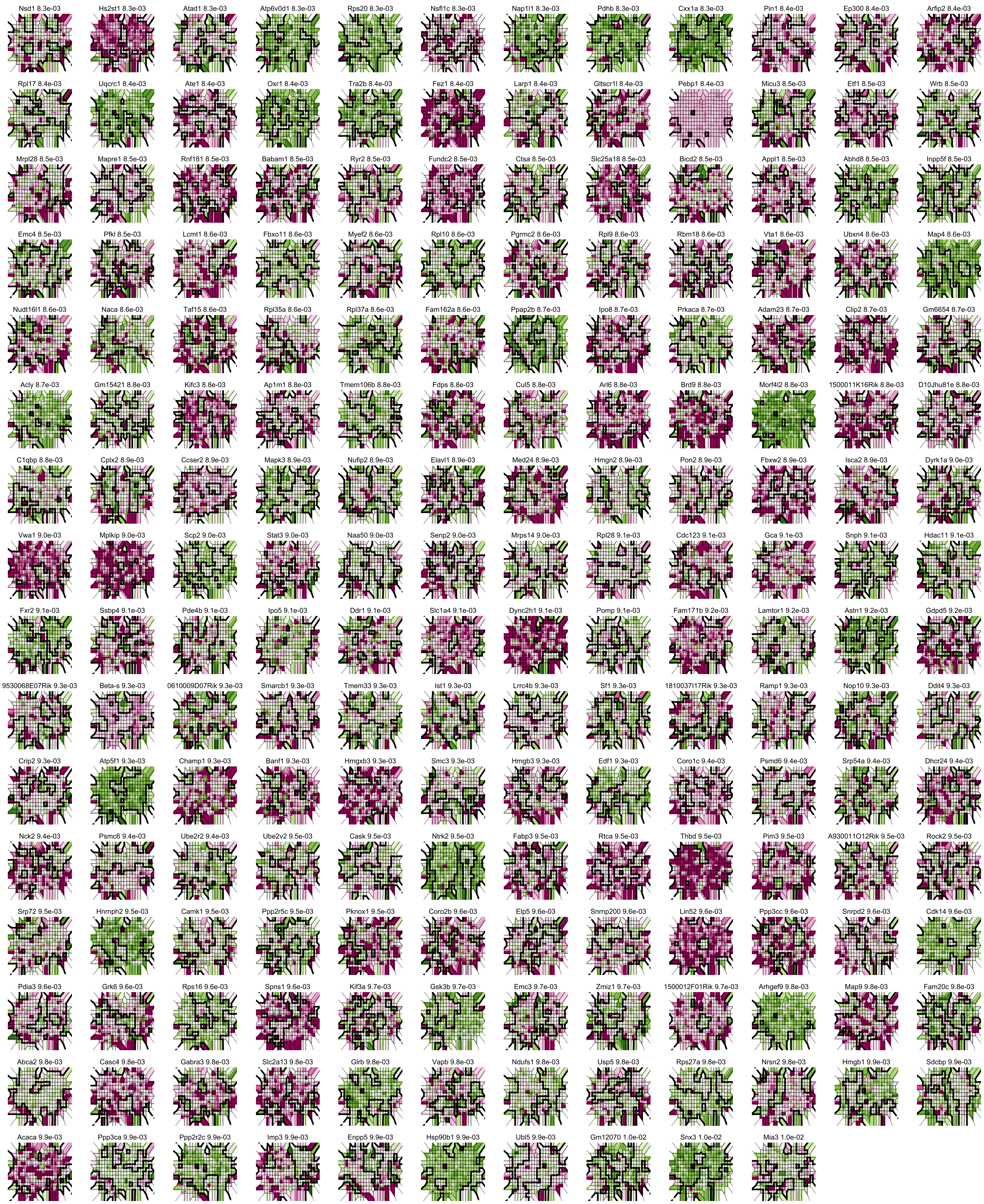
Nsd1 8.3e-03
Hs2st1 8.3e-03
Atad1 8.3e-03
Atp6v0d1 8.3e-03
Rps20 8.3e-03
Nsf1c 8.3e-03
Nap1l1 8.3e-03
Pdhb 8.3e-03
Cxx1a 8.3e-03
Pin1 8.4e-03
Ep300 8.4e-03
Arfip2 8.4e-03
Rpl17 8.4e-03
Uqcrc1 8.4e-03
Ate1 8.4e-03
Oxr1 8.4e-03
Tra2b 8.4e-03
Fez1 8.4e-03
Larp1 8.4e-03
Gltscr1l 8.4e-03
Pebp1 8.4e-03
Micu3 8.5e-03
Etf1 8.5e-03
Wrb 8.5e-03
Mrpl28 8.5e-03
Mapre1 8.5e-03
Rnf181 8.5e-03
Babam1 8.5e-03
Ryr2 8.5e-03
Fundc2 8.5e-03
Ctsa 8.5e-03
Slc25a18 8.5e-03
Bicd2 8.5e-03
Appl1 8.5e-03
Abhd8 8.5e-03
Inpp5f 8.5e-03
Emc4 8.5e-03
Pfkl 8.5e-03
Lcmt1 8.6e-03
Fbxo11 8.6e-03
Myef2 8.6e-03
Rpl10 8.6e-03
Pgrmc2 8.6e-03
Rpl9 8.6e-03
Rbm18 8.6e-03
Vta1 8.6e-03
Ubxn4 8.6e-03
Map4 8.6e-03
Nudt16l1 8.6e-03
Naca 8.6e-03
Taf15 8.6e-03
Rpl35a 8.6e-03
Rpl37a 8.6e-03
Fam162a 8.6e-03
Ppap2b 8.7e-03
Ipo8 8.7e-03
Prkaca 8.7e-03
Adam23 8.7e-03
Clip2 8.7e-03
Gm6654 8.7e-03
Acly 8.7e-03
Gm15421 8.8e-03
Kifc3 8.8e-03
Ap1m1 8.8e-03
Tmem106b 8.8e-03
Fdps 8.8e-03
Cul5 8.8e-03
Arl6 8.8e-03
Brd9 8.8e-03
Morf4l2 8.8e-03
1500011K16Rik 8.8e-03
D10Jhu81e 8.8e-03
C1qbp 8.8e-03
Cplx2 8.9e-03
Ccser2 8.9e-03
Mapk3 8.9e-03
Nufip2 8.9e-03
Elavl1 8.9e-03
Med24 8.9e-03
Hmgn2 8.9e-03
Pon2 8.9e-03
Fbxw2 8.9e-03
Isca2 8.9e-03
Dyrk1a 9.0e-03
Vwa1 9.0e-03
Mplkip 9.0e-03
Scp2 9.0e-03
Stat3 9.0e-03
Naa50 9.0e-03
Senp2 9.0e-03
Mrps14 9.0e-03
Rpl28 9.1e-03
Cdc123 9.1e-03
Gca 9.1e-03
Snph 9.1e-03
Hdac11 9.1e-03
Fxr2 9.1e-03
Ssbp4 9.1e-03
Pde4b 9.1e-03
Ipo5 9.1e-03
Ddr1 9.1e-03
Slc1a4 9.1e-03
Dync2h1 9.1e-03
Pomp 9.1e-03
Fam171b 9.2e-03
Lamtor1 9.2e-03
Astn1 9.2e-03
Gdpd5 9.2e-03
9530068E07Rik 9.3e-03
Beta-s 9.3e-03
0610009D07Rik 9.3e-03
Smarcb1 9.3e-03
Tmem33 9.3e-03
Ist1 9.3e-03
Lrrc4b 9.3e-03
Sf1 9.3e-03
1810037I17Rik 9.3e-03
Ramp1 9.3e-03
Nop10 9.3e-03
Ddit4 9.3e-03
Crip2 9.3e-03
Atp5f1 9.3e-03
Champ1 9.3e-03
Banf1 9.3e-03
Hmgxb3 9.3e-03
Smc3 9.3e-03
Hmgb3 9.3e-03
Edf1 9.3e-03
Coro1c 9.4e-03
Psmd6 9.4e-03
Srp54a 9.4e-03
Dhcr24 9.4e-03
Nck2 9.4e-03
Psmc6 9.4e-03
Ube2r2 9.4e-03
Ube2v2 9.5e-03
Cask 9.5e-03
Ntrk2 9.5e-03
Fabp3 9.5e-03
Rtca 9.5e-03
Thbd 9.5e-03
Pim3 9.5e-03
A930011O12Rik 9.5e-03
Rock2 9.5e-03
Srp72 9.5e-03
Hnrnph2 9.5e-03
Camk1 9.5e-03
Ppp2r5c 9.5e-03
Pknox1 9.5e-03
Coro2b 9.6e-03
Elp5 9.6e-03
Snrnp200 9.6e-03
Lin52 9.6e-03
Ppp3cc 9.6e-03
Snrpd2 9.6e-03
Cdk14 9.6e-03
Pdia3 9.6e-03
Grk6 9.6e-03
Rps16 9.6e-03
Spns1 9.6e-03
Kif3a 9.7e-03
Gsk3b 9.7e-03
Emc3 9.7e-03
Zmiz1 9.7e-03
1500012F01Rik 9.7e-03
Arhgef9 9.8e-03
Map9 9.8e-03
Fam20c 9.8e-03
Abca2 9.8e-03
Casc4 9.8e-03
Gabra3 9.8e-03
Slc2a13 9.8e-03
Glrb 9.8e-03
Vapb 9.8e-03
Ndufs1 9.8e-03
Usp5 9.8e-03
Rps27a 9.8e-03
Nrsn2 9.8e-03
Hmgb1 9.9e-03
Sdcbp 9.9e-03
Acaca 9.9e-03
Ppp3ca 9.9e-03
Ppp2r2c 9.9e-03
Imp3 9.9e-03
Enpp5 9.9e-03
Hsp90b1 9.9e-03
Ubl5 9.9e-03
Gm12070 1.0e-02
Snx3 1.0e-02
Mia3 1.0e-02
